# Insights into the conservation and diversification of the molecular functions of YTHDF proteins

**DOI:** 10.1101/2022.10.19.512826

**Authors:** Daniel Flores-Téllez, Mathias Due Tankmar, Sören von Bülow, Junyu Chen, Kresten Lindorff-Larsen, Peter Brodersen, Laura Arribas-Hernández

**Affiliations:** University of Copenhagen, Biology Department. Ole Maaløes Vej 5, 2200 Copenhagen N, Denmark; Universidad Francisco de Vitoria, Facultad de Ciencias Experimentales. Ctra. Pozuelo-Majadahonda km 1800, 28223 Pozuelo de Alarcón (Madrid), Spain

**Author notes:** Corresponding authors (P.B.) and (L.A.-H.). The authors declare that they have no conflict of interest.

## Abstract

YT521-B homology (YTH) domain proteins act as readers of *N6*-methyladenosine (m^6^A) in mRNA. Members of the YTHDF clade determine properties of m^6^A-containing mRNAs in the cytoplasm. Vertebrates encode three YTHDF proteins whose possible functional specialization is debated. In land plants, the YTHDF clade has expanded from one member in basal lineages to eleven so-called EVOLUTIONARILY CONSERVED C-TERMINAL REGION1-11 (ECT1-11) proteins in *Arabidopsis thaliana*, named after the conserved YTH domain placed behind a long N-terminal intrinsically disordered region (IDR). *ECT2*, *ECT3* and *ECT4* show genetic redundancy in stimulation of primed stem cell division, but the origin and implications of YTHDF expansion in higher plants are unknown, as it is unclear whether it involves acquisition of fundamentally different molecular properties, in particular of their divergent IDRs. Here, we use functional complementation of *ect2*/*ect3*/*ect4* mutants to test whether different YTHDF proteins can perform the same function when similarly expressed in leaf primordia. We show that stimulation of primordial cell division relies on an ancestral molecular function of the m^6^A-YTHDF axis in land plants that is present in bryophytes and is conserved over YTHDF diversification, as it appears in all major clades of YTHDF proteins in flowering plants. Importantly, although our results indicate that the YTH domains of all arabidopsis ECT proteins have m^6^A-binding capacity, lineage-specific neo-functionalization of ECT1, ECT9 and ECT11 happened after late duplication events, and involves altered properties of both the YTH domains, and, especially, of the IDRs. We also identify two biophysical properties recurrent in IDRs of YTHDF proteins able to complement *ect2 ect3 ect4* mutants, a clear phase separation propensity and a charge distribution that creates electric dipoles. Human and fly YTHDFs do not have IDRs with this combination of properties and cannot replace ECT2/3/4 function in arabidopsis, perhaps suggesting different molecular activities of YTHDF proteins between major taxa.

**Author Summary:** Regulation of gene expression is essential to life. It ensures correct balancing of cellular activities and the controlled proliferation and differentiation necessary for the development of multicellular organisms. Methylation of adenosines in mRNA (m^6^A) contributes to genetic control, and absence of m^6^A impairs embryo development in plants and vertebrates. m^6^A-dependent regulation can be exerted by a group of cytoplasmic proteins called YTHDFs. Higher plants have many more YTHDFs than animals, but it is unknown whether these many YTHDF proteins carry out fundamentally different or roughly the same molecular functions, as only a small fraction of them have been studied thus far. This work addresses the origin and reasons behind YTHDF expansion during land plant evolution and reveals that most YTHDFs in flowering plants have the same molecular functions that facilitate rapid division of differentiating stem cells. Remarkably, this molecular activity is present in the most basal lineage of land plants and functions similarly across 450 million years of land plant evolution. We also identified a few plant YTHDF proteins with divergent molecular function despite their ability to bind to m^6^A. Our work provides a firm basis for further advances on understanding molecular properties and biological contexts underlying YTHDF diversification.

## Introduction

*N6*-methyladenosine (m^6^A) is the most abundant modified nucleotide occurring internally in eukaryotic mRNA. It is of major importance in gene regulation as illustrated by the embryonic lethality of mutants in the dedicated mRNA adenosine methyltransferase in higher plants [1] and in mammals [2]. The presence of m^6^A in an mRNA may have multiple biochemical consequences. These include changes in the secondary structure [3–5] and the creation of binding sites for RNA-binding proteins specialized for m^6^A recognition [6, 7]. YT521-B homology (YTH) domain proteins [8] constitute the best studied class of m^6^A-binding proteins [6, 7, 9]. They achieve specificity for m^6^A via an aromatic pocket accommodating the *N6*-adenosine methyl group, such that the affinity of isolated YTH domains for m^6^A-containing RNA is 10-20-fold higher than for unmodified RNA [9–14].

Two different phylogenetic clades of YTH domains have been defined, YTHDC and YTHDF, sometimes referred to simply as DC and DF [6, 15]. Genetic studies show that major functions of m^6^A in development in both vertebrates and higher plants depend on the YTHDF clade of readers [16–22]. In all cases studied in detail thus far, plant and animal YTHDF proteins are cytoplasmic in unchallenged conditions [15, 16, 23–26], and contain a long N-terminal intrinsically disordered region (IDR) in addition to the C-terminal YTH domain [6, 8, 27]. While the YTH domain is necessary for specific binding to m^6^A in mRNA [9–14], the IDR is considered to be the effector part of the protein [9, 28–30]. Nonetheless, it has been proposed that the IDR may also participate in RNA binding, because the YTH domain alone has low affinity for mRNA [6]. Indeed, the IDR-dependent crosslinks between a YTHDF protein and mRNA detected upon UV-irradiation of living arabidopsis seedlings [31] experimentally supports such a mechanism, conceptually equivalent to the contribution of IDRs in transcription factors to specific DNA binding [32]. Additionally, the IDR may be involved in phase separation. Plant YTHDF proteins can form condensates *in vitro* [16] and, upon stress, localize *in vivo* to distinct foci [16, 26] identified as stress granules [26]. Similar properties have been reported for human YTHDFs, which engage in liquid-liquid phase separation when concentrated on polyvalent m^6^A-modified target RNA *in vitro* [33–36].

While yeast, flies and primitive land plants encode only one YTHDF protein [24–26, 37, 38], vertebrates have three closely related paralogs (YTHDF1-3) [6, 8, 15, 39] and higher plants encode an expanded family, with eleven members in *Arabidopsis thaliana* (*Ath*) referred to as EVOLUTIONARILY CONSERVED C-TERMINAL REGION1-11 (ECT1-11) [27, 40]. It is a question of fundamental importance for the understanding of how complex eukaryotic systems use the regulatory potential of m^6^A whether these many YTHDF proteins perform the same biochemical function, or whether their molecular properties are specialized. Such molecular specialization could, for instance, arise as a consequence of differential binding specificity to mRNA targets, distinct protein-protein interaction properties, or distinct biophysical properties of IDRs, such as those related to the propensity to phase separate [41, 42]. Alternatively, diversification of biological functions could be achieved with distinct expression patterns or induction by environmental cues, even if targets and molecular functions are the same. The studies on YTHDF specialization in vertebrate cells illustrate the importance of the topic, and, equally, the importance of addressing it rigorously [43]. Initial work on mammalian cell cultures advocated a model in which YTHDF1 would enhance translation of target mRNAs, YTHDF2 would promote mRNA decay, and YTHDF3 would be able to trigger either of the two [9, 44–46]. Nonetheless, subsequent research work in mouse, zebrafish and human cell culture involving single and combined *ythdf* knockouts and analysis of interacting mRNAs and proteins [18, 22, 39], and comparative studies of human YTHDF1/2/3 structure and molecular dynamics [47] do not support functional specialization, and propose a unified molecular function for all three vertebrate YTHDFs in accelerating mRNA decay. Whether metazoan YTHDF1/2/3 are molecularly redundant or not, and what their precise molecular functions may be, are topics that continue to be debated as of today [48].

The great expansion of YTHDF proteins in higher plants is unique among eukaryotes [8]. Phylogenetic analyses of plant YTHDF domains have established the existence of 3 clades in angiosperms, DF-A (comprising *Ath* ECT1, ECT2, ECT3, ECT4), DF-B (comprising *Ath* ECT5, ECT10, ECT9), and DF-C (comprising *Ath* ECT6, ECT7, ECT8 and ECT11) [26]. The fact that the eleven arabidopsis paralogs have the aromatic residues necessary for specific binding to m^6^A [49] suggests that they may all function as m^6^A readers. The unified model for YTHDF function recently proposed for the three vertebrate paralogs [18, 22, 39] is consistent with what had already been established for *Ath* ECT2, ECT3 and ECT4 in the plant DF-A clade [16], even though the three plant paralogs are more divergent in sequence than the highly similar mammalian YTHDF1-3 [8, 16, 39]. The two most highly expressed members in *A. thaliana*, ECT2 and ECT3, accumulate in dividing cells of organ primordia and exhibit genetic redundancy in the stimulation of stem cell proliferation during organogenesis [16, 17]. The two proteins probably act truly redundantly *in vivo* to control this process, because they associate with highly overlapping target sets in wild type plants, and each exhibits increased target mRNA occupancy in the absence of the other protein [23]. Simultaneous knockout of *ECT2* and *ECT3* causes a 2-day delay in the emergence of the first true leaves, aberrant leaf morphology, slow root growth and defective root growth directionality among other defects [16, 17] that resemble those of plants with diminished m^6^A deposition [50–52]. For the third DF-A clade member, *ECT4*, the genetic redundancy is only noticeable in some tissues as an exacerbation of *ect2/ect3* phenotypes upon additional mutation of *ECT4*, most conspicuously seen in leaf morphogenesis [16, 17]. Despite the strong evidence for redundant functions among these three YTHDF paralogs in the DF-A clade, the presence of many other YTHDF proteins in arabidopsis leaves open the question of whether substantial functional specialization of YTHDF proteins exists in plants.

In this study, we systematically define overlaps in the molecular functions of YTHDF proteins in land plants (embryophytes). Employing functional complementation of leaf formation in triple *ect2/ect3/ect4* (*te234*) mutants, we demonstrate that at least one member of all clades in arabidopsis, and the only YTHDF protein from a group of primitive land plants, can replace ECT2/3/4 functions. In contrast, a few late-diverging ECTs were not able to perform the molecular functions of ECT2/3/4. Based on these results, we propose an ancestral molecular role of land plant YTHDF proteins in stimulation of primordial cell proliferation, and sustained functional redundancy during the diversification process that started more than 400 <u>m</u>illion <u>y</u>ears <u>a</u>go (Mya). In addition, our results also support the functional divergence of a small subset of fast-evolving plant YTHDF proteins with contributions to specialization mainly from the IDRs, but also from the YTH domains.

## Results

### The phylogeny of YTHDF proteins comprises several clades in land plants

The adaptation of plants to terrestrial life was accompanied by the acquisition of morphological, physiological, and genomic complexity [53–55]. Knowing how diversification of YTHDF proteins came about during the course of plant evolution is relevant to understand their distinct functions and may hint at roles they might have played in this process. However, the species included in the so far most detailed phylogenetic study on plant YTHDF proteins jump from bryophytes (mosses, liverworts and hornworts [54, 56–60]) to angiosperms (flowering plants) [26]. To include the YTHDF repertoires of intermediately diverging clades of embryophytes such as lycophytes, ferns, and gymnosperms, we performed phylogenetic analyses using YTHDF proteins from 32 species widely spread across land plant evolution. We also included YTHDFs from 8 species of green algae (chlorophytes and charophytes) and, as outgroup, 3 microalgal species from eukaryotic taxa evolutionarily close to plants (Fig 1A). In the resulting phylogenetic tree, all embryophyte YTHDFs branch from a single stem that connects this monophyletic group to the more divergent green algal orthologs (Figs 1B and S1-S3), whose peculiarities are further considered in the discussion. Regarding land plants, the tree largely agrees with the DF-A/B/C clades defined on the basis of angiosperm YTHDF sequences [26], with some minor differences. We consider that the former DF-C group comprising *Ath* ECT6/7/8/11 [26] can be subdivided into two groups that diverged early during YTHDF radiation (Figs 1B and S1). Thus, we introduced the name ‘DF-D’ for the clade defined by one *Amborella trichopoda* (a basal angiosperm) YTHDF protein (*Atr* DFD) and *Ath* ECT6/7 (Figs 1B and S1).

**Fig 1.**
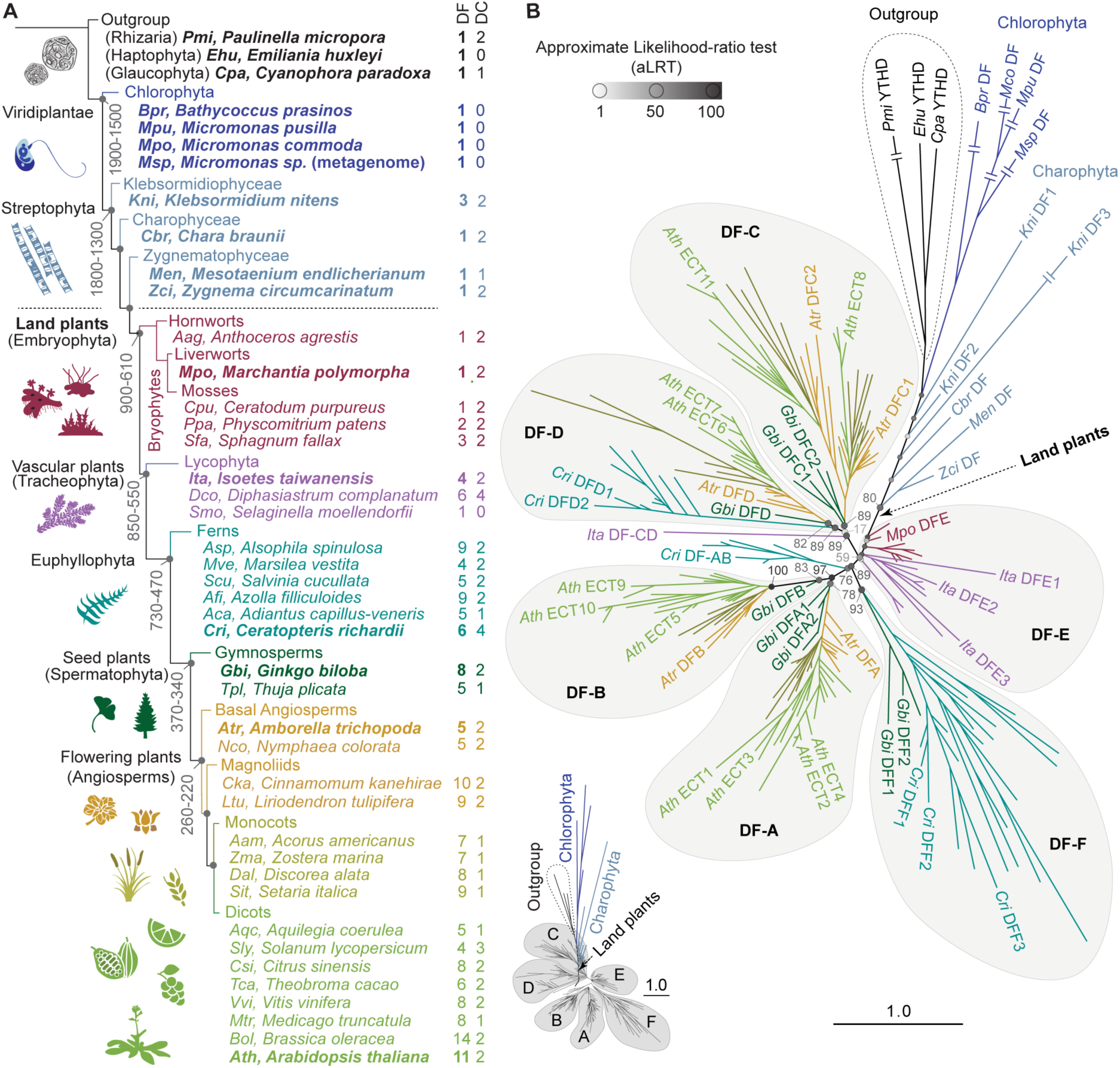
Phylogenetic analysis of YTHDF proteins in plants. **(A)** Schematic representation of Viridiplantae evolution and relative position of the species used in this study. The architecture of the diagram and the age (million years, My) indicated on some nodes are a simplified version of the trees from Su et al. [59] and Bowman [62]. The length of the branches has been adjusted for illustrative purposes. **(B)** Phylogenetic tree of YTHDF proteins in plants (S1-S4 Datasets), with color-coding and abbreviations of species names as in A. Three YTHDF proteins from taxa outside but evolutionarily close to Viridiplantae are included as outgroup. *Arabidopsis thaliana* (*Ath*) YTHDF proteins are named after the original nomenclature for proteins containing an Evolutionarily Conserved C-Terminal Region (ECT) established by Ok et al. [40]. Proteins from other plant species adhere to the nomenclature established by Scutenaire et al. [26], with small variations reflecting additional clades (DF-E, -F) and the split of the original DF-C clade into -C and -D. Statistical support is calculated as approximate likelihood-ratio test (aLRT), and indicated by grayscale-coded spheres and values on the most relevant nodes. For simplicity, only a subset of proteins is labelled–those from one representative species of the main taxa highlighted in bold in A–, but a comprehensive representation of the same tree with all protein names and aLRT values is available in S1 Fig. The length of the branches represents the evolutionary distance in number of amino acid substitutions per site relative to the scale bar. Double lines crossing the longest branches indicate that the branch has been collapsed due to space problems, but a schematic representation of their relative lengths is shown on the lower-left corner, and not-collapsed branches can be found in S1 Fig.

Furthermore, since the group comprising bryophyte YTHDFs did not receive a designation in previous studies, we named it DF-E (mnemotechnic for ‘Early’). It includes YTHDF proteins from bryophytes and some lycophyte orthologs that branched close to the moss subgroup (Figs 1B and S1). Finally, an additional group composed of YTHDF proteins from six species of ferns and two gymnosperms was named DF-F, as they did not fall into any of the other clades (Figs 1B and S1). Interestingly, the alignment revealed independent gains and losses of molecular traits during plant YTHDF evolution. For example, an aspartate-to-asparagine substitution in helix α1 that increases affinity for m^6^A [61], characteristic of DC and plant DF-B YTH domain proteins [26, 61], is also present in the fern proteins of the DF-F clade, but absent in DF-Es, DF-As and gymnosperm DF-Fs (Fig S4), suggesting independent losses or acquisitions. Taking these observations together, we conclude that land plant YTHDFs are phylogenetically more diverse than previously appreciated.

### YTHDF protein diversification occurred early during land plant evolution

Our phylogenetic analysis shows that plant YTHDF diversification started at least before the radiation of Euphyllophytes (plants with true leaves comprising ferns, gymnosperms and angiosperms) more than 410 Mya [59, 63]. This is because bryophytes possess one or two YTHDFs in the ‘early clade’ (DF-E), while all six fern species analyzed here have extended sets DF proteins (4-9 paralogs) that spread into clades DF-F, DF-D, and the branch sustaining the DF-A and DF-B groups (Fig 1B). Furthermore, because one DF-protein from the lycophyte *Isoetes taiwanensis* branches from the common stem between clades DF-C and DF-D (*Ita* DF-CD), it is likely that a first diversification event started in the ancestral vascular plants. Thus, our analysis reveals that YTHDF radiation started early in land plant evolution and coincided with the acquisition of morphological complexity and the adaptation to diverse environments.

### A functional complementation assay for plant YTHDF proteins

To address the degree of functional specialization among plant YTHDF proteins in a simple manner, we set up a functional complementation assay that scores the ability of each of the eleven arabidopsis YTHDF proteins (Fig 2A) to perform the molecular functions of ECT2/3/4 required for rapid cellular proliferation in leaf primordia. Initial attempts at using the *ECT2* promoter for ectopic *ECT* expression in primordial cells were unsuccessful, as pilot experiments with a genomic *ECT4* fragment revealed that its expression was substantially lower than that of a similar *ECT2* genomic fragment when driven by the *ECT2* promoter (S5 Fig), perhaps pointing to the presence of internal *cis*-regulatory elements. We therefore turned to the use of *ECT* cDNAs (c*ECT*s) under the control of the promoter of the ribosomal protein-encoding gene *RPS5A/uS7B/US7Y* (At3g11940 [64, 65]), henceforth *US7Yp*, active in dividing cells [66] in a pattern that resembles the *ECT2/3/4* expression domain [16, 17]. Transformation of the *US7Yp:cECT2-mCherry* construct in *te234* plants resulted in complementation frequencies of ∼40-50% among primary transformants, a percentage slightly lower but comparable to that obtained with the genomic *ECT2-mCherry* construct under the control of *ECT2* promoter and terminator regions (*ECT2p:gECT2-mCherry*) [16] (S6A Fig). Free mCherry expressed from a *US7Yp:mCherry* transgene showed no complementation (S6A Fig), as expected. Importantly, the leaf morphology and the pattern of mCherry fluorescence in *US7Yp:cECT2-mCherry* lines was indistinguishable from that of *ECT2p:gECT2-mCherry* (S6B Fig). Hence, we proceeded with expression of cDNAs encoding all of ECT1-11 fused to C-terminal mCherry under the control of the *US7Y* promoter (henceforth ECT1-11) in *te234* mutants (Fig 2B).

**Fig 2.**
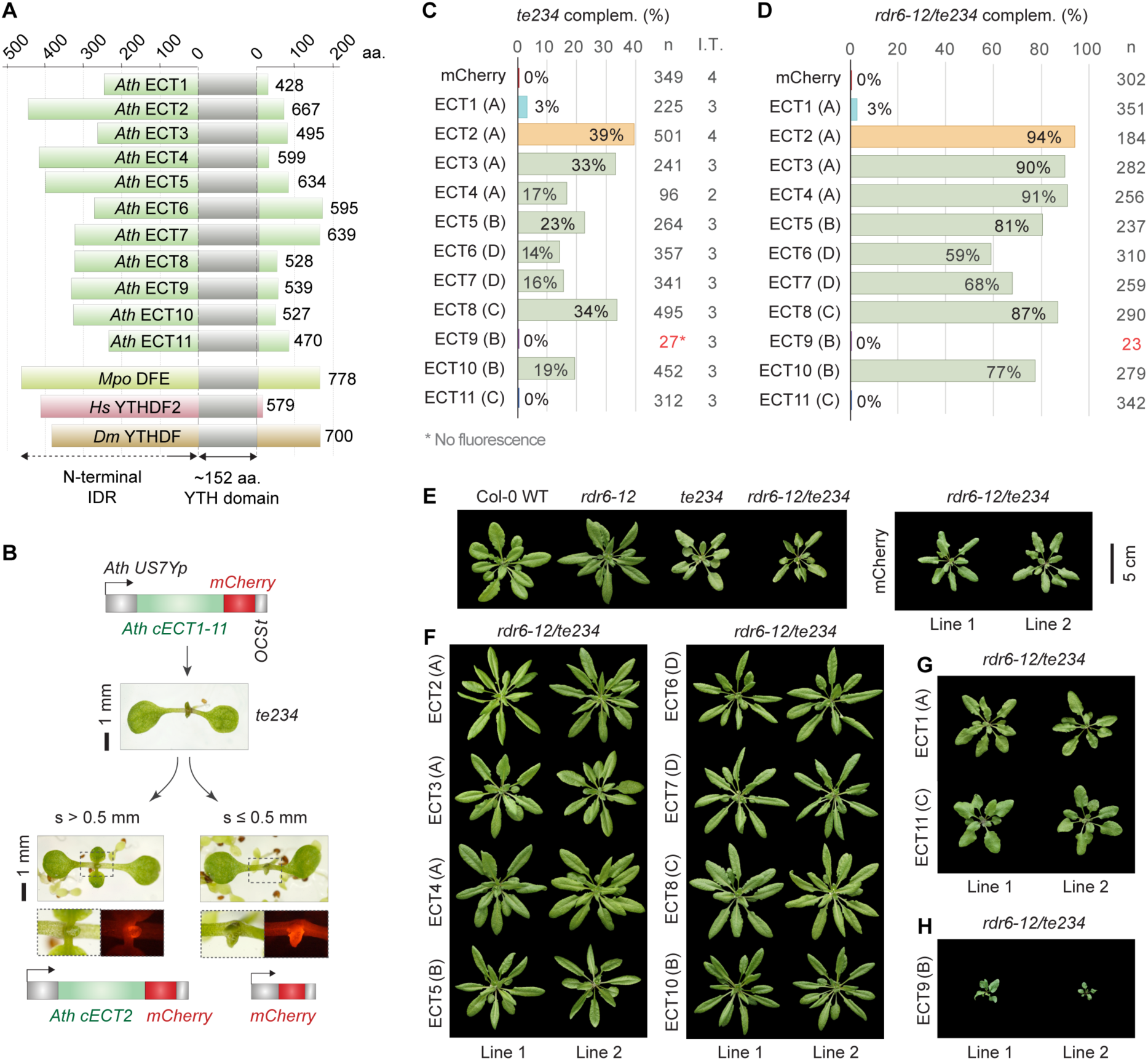
Most but not all arabidopsis YTHDF proteins can replace ECT2/3/4-function. **(A)** Diagram showing the relative length of the N-terminal IDRs of *Arabidopsis thaliana* (*Ath*) YTHDF proteins (ECT1-ECT11) together with *Marchantia polymorpha* (*Mpo*) DFE, *Homo sapiens* (*Hs*) YTHDF2, and *Drosophila melanogaster* (*Dm*) YTHDF. Numbers indicate the length of the proteins in amino acids (aa). **(B)** Strategy followed for the functional assay. *US7Yp:cECT(X)-mCherry* constructs are introduced in *ect2-1/ect3-1/ect4-2* (*te234*) plants, and complementation rates are estimated by the percentage of primary transformants (T1) whose first true leaves have a size (s) exceeding 0.5 mm after 10 days of growth. The *construct US7Yp:mCherry* is used as negative control. Examples of T1 seedlings expressing control and ECT2 constructs are shown. Dashed outlines are magnified below to show mCherry fluorescence in emerging leaves. **(C)** Weighed averages of the complementation percentages observed for each *US7Yp:cECT(X)-mCherry* construct in 2-5 independent transformations (I.T.). Letters between parenthesis indicate the DF-clade assigned to each protein. Colouring highlights ECT2 (reference) and proteins with low or negligible complementation capacity (ECT1/9/11) for fast identification along the manuscript. Low transformation efficiency (T.E.) for ECT9 can be inferred by the low number of transformants (n) highlighted in red, but detailed T.E. and raw complementation rates in each transformation can be found in S9 and S7A Figs respectively. Additional data regarding this assay is shown in S8 Fig. **(D)** Same as C on a single transformation using the *rdr6-12/te234* background. **(E-H)** Rosette phenotype of 32-day old T1s bearing *US7Yp:cECT(X)-mCherry* constructs in the *rdr6-12/te234* background, sorted according to their degree of complementation in three panels: F (high), G (residual) and H (non-complementation and enhancement). Different genetic backgrounds and control transformants (mCherry) are shown in *E* as a reference. The scalebar (5 cm) applies to all plants in the four panels.

### Percentage of complementation among primary transformants as a readout for functionality

To design the experimental approach in detail, we considered the fact that expression of transgenes involves severe variations in levels and patterns among transformants. This is due to positional effects of the T-DNA insertion and the propensity of transgenes to trigger silencing in plants [67], and explains why only a fraction of *ECT2-mCherry* lines rescues loss of ECT2 function (S6A Fig, [16]). Hence, many independent lines need to be analyzed before choosing stable and representative lines for further studies. Taking this into account, we decided to perform systematic and unbiased comparisons of ECT functionality by counting the fraction of primary transformants (henceforth T1s) able to complement the late leaf emergence of *te234* plants for each construct. We set a size threshold for the rapid assessment of large numbers of transformants considering that leaves longer than 0.5 mm after 10 days of growth constitute unambiguous sign of complementation, because this is the minimum size never reached by *te234* or control lines (S2B Fig). Hence, we calculated complementation percentages as the number of T1s with leaf size (s) larger than 0.5 mm relative to the total number of transformants, and averaged that score over several independent transformations (Figs 2C and S7A, S8A).

### Most arabidopsis YTHDF paralogs can perform the molecular function required for leaf development

The results of the comparative analysis reveal a high degree of functional overlap within the arabidopsis YTHDF family, because eight out of the eleven YTHDF proteins, namely ECT2/3/4/5/6/7/8/10, complement the delayed leaf emergence of *ect2/3/4* mutants with frequencies ranging from 14% (ECT6) to 39% (ECT2) when expressed in the same cells (Fig 2C). Importantly, complementation is indirect proof of m^6^A-binding ability, because the proteins must bind to the targets whose m^6^A-dependent regulation is necessary for correct leaf organogenesis in order to restore ECT2/3/4 function [16]. This conclusion is not trivial, as some eukaryotes lacking canonical m^6^A-methyltransferases encode YTH domain proteins [68] and, indeed, the YTH domain of fission yeast Mmi1 does not bind to m^6^A [69]. Thus, most, but not all, arabidopsis YTHDF proteins retain the molecular functions required to stimulate proliferation of primed stem cells. Of the three remaining proteins, ECT1 showed only residual complementation capacity, while no complementation was observed for ECT11 at this stage (Fig 2C). Finally, we could not conclude at this point whether ECT9 could replace ECT2/3/4 function or not, because this transgene produced very few transformants in the *te234* background with barely visible fluorescence (Figs 2C and S7-S9). All such transformants underwent complete silencing in the following generation (T2).

### Differences in complementation scores are not explained by a bias in expression levels

Our T1 scoring method relies on the assumption that the *US7Y* promoter drives similar ranges of expression of the different *ECT* cDNAs. However, different silencing propensities between the *ECT* cDNAs might introduce bias in the results. To take this possibility into account, we first performed western blotting from three lines per transgene selected as the best-complementing in their group (S8B Fig). The results showed that the higher complementation scores of e.g. ECT2, ECT3 and ECT8 are not explained by higher expression. Rather, the reverse was true. For example, while ECT2 could complement *te234* plants with the lowest expression among all ECTs, the few lines expressing ECT1 able to pass the complementation threshold had the highest expression of all ECTs, suggesting that ECT1 can only partially compensate for loss of ECT2/3/4 when overexpressed.

### The same functional classes of ECTs emerge from complementation of silencing-deficient te234

Next, we tested whether the same results could be reproduced in plants lacking *RNA-DEPENDENT RNA POLYMERASE 6* (*RDR6*), a gene essential for the biogenesis of transgene-derived short interfering RNAs (siRNAs) [70, 71]. Because siRNA-guided transgene silencing is impaired in *rdr6* mutants, higher and more homogenous ranges of expression among different transgenes can be expected in this background. Hence, we constructed *rdr6-12/te234* mutants that exhibited both the narrow, downward-curling leaf phenotype of *rdr6* mutants [72] and the characteristic rosette morphology of *te234* [16] (S10A Fig). Importantly, the delay in leaf emergence was identical in *te234* and *rdr6-12/te234* mutants (S10A Fig) and the complementation rates of *ECT2* transgenes were much higher in the silencing-deficient plants (S6A Fig), indicating that the <100% complementation frequencies in *te234* were indeed due to RDR6-dependent silencing. In the *rdr6*/*te234* background, complementation scores of the eight functionally equivalent ECTs (ECT2/3/4/5/6/7/8/10) ranged from 59% (ECT6) to 94% (ECT2), and the differences between these ECTs were less pronounced, but followed a trend largely unchanged compared to the *te234* background (Fig 2C,D). On the other hand, the difference between the eight ECTs with ECT2/3/4-like function and the group of ECTs with negligible complementation capacity (ECT1/9/11) was highlighted by the *rdr6* mutation: ECT1 and ECT11 did not increase their scores (Fig 2D), and absence of silencing resulted in a number of *ECT9* transformants with visible mCherry fluorescence, yet no complementation activity (Figs 2D and S10B), thus verifying the lack of ECT2/3/4-like function of ECT9.

### Rosette phenotypes corroborate the complementation scores

To verify the results of the complementation assay in the adult phenotype, we characterized rosettes of two independent lines for each transgene (Fig 2E-H). As expected, *rdr6-12/te234* plants expressing ECTs with high complementation scores resembled single *rdr6-12* mutants (Fig 2E,F), while the ECT1 and ECT11 transformants with the biggest first true leaves at 10 DAG produced plants only slightly bigger than the *rdr6-12/te234* background (Fig 2E,G), indicating low but residual ECT2/3/4-like function for these two proteins. Strikingly, *rdr6-12/te234* plants expressing ECT9 exhibited strong developmental phenotypes (Figs 2H and S10B) that resembled those of severely m^6^A-depleted plants [51, 52]. Such an effect was not observed for any other *Ath* ECT. This result suggests that ECT9 can bind to m^6^A targets, either exerting opposite molecular actions compared to ECT2/3/4, or simply precluding their possible regulation by remaining ECTs such as ECT5/6/7/8/10 by competitive binding. We tested whether ECT9 can outcompete endogenous ECT2/3/4 in the proliferating cells of the apex by expressing the ECT9 transgene in single *rdr6-12* mutant plants. We found no evidence for this, because the transformation produced many fluorescent T1s without obvious defects (S11 Fig). Altogether, our results indicate that arabidopsis YTHDF proteins may be divided into three functional classes, i) ECT2/3/4/5/6/7/8/10 with the molecular functionality of ECT2/3/4 required for correct and timely leaf formation; ii) ECT1/11, with only residual ECT2/3/4 functions, and iii) ECT9, unable to perform such functions and toxic when present in the expression domain of ECT2/3/4 in their absence.

### Validation of the functional assay by loss-of-function genetic analysis

One of the predictions on genetic interactions between *Ath ECT* genes that emerges from our results is particularly unexpected and, therefore, suitable for the validation of our complementation assay by classical genetic analysis: Mutation of *ECT1* should not exacerbate the developmental defects of *te234*, even if it is endogenously expressed in leaf primordia. *ECT1* lends itself well to validation for two reasons. First, ECT1 is the closest paralog of ECT2/3/4 (Fig 1B), intuitively suggesting some degree of genetic redundancy. Second, *ECT1* promoter-reporter fusions [40] and mRNA-seq data ([73], S12 Fig) suggest that expression levels and tissue specificity of *ECT1* are comparable to that of *ECT4*, whose activity is easily revealed as an enhancer mutant of *ect2/ect3* [16].

### ECT1 and ECT2/3/4 are cytoplasmic and are expressed in the same cell types

We first assessed the expression pattern and subcellular localization of ECT1 using stable transgenic lines expressing TFP translationally fused to the C-terminus of *ECT1* (*ECT1-TFP*, S13A Fig). The pattern of turquoise fluorescence was strongly reminiscent of that of fluorescent ECT2/3/4 fusions [16] with signal at the base of young leaves, in leaf primordia, main root tips and lateral root primordia at different stages (Fig 3A-D). In addition, confocal microscopy of meristematic root cells showed cytoplasmic ECT1-TFP signal with heterogenous texture similar to ECT2/3/4 [16] (Fig 3E). We noticed, however, that most root tips had a few cells containing distinct ECT1-TFP foci (Fig 3F) in the absence of stress. We found no trace of free TFP that could be causing imaging artifacts (S13A Fig) but note that the *ECT1-TFP* transgenic lines selected on the basis of visible and reproducible pattern of fluorescence may overexpress *ECT1* (S13B Fig). We conclude from the similar expression pattern and subcellular localization of ECT1 and ECT2/3/4 that assessment of the possible exacerbation of *te234* phenotypes by additional loss of *ECT1* function is a meaningful test of our functional assay.

**Fig 3.**
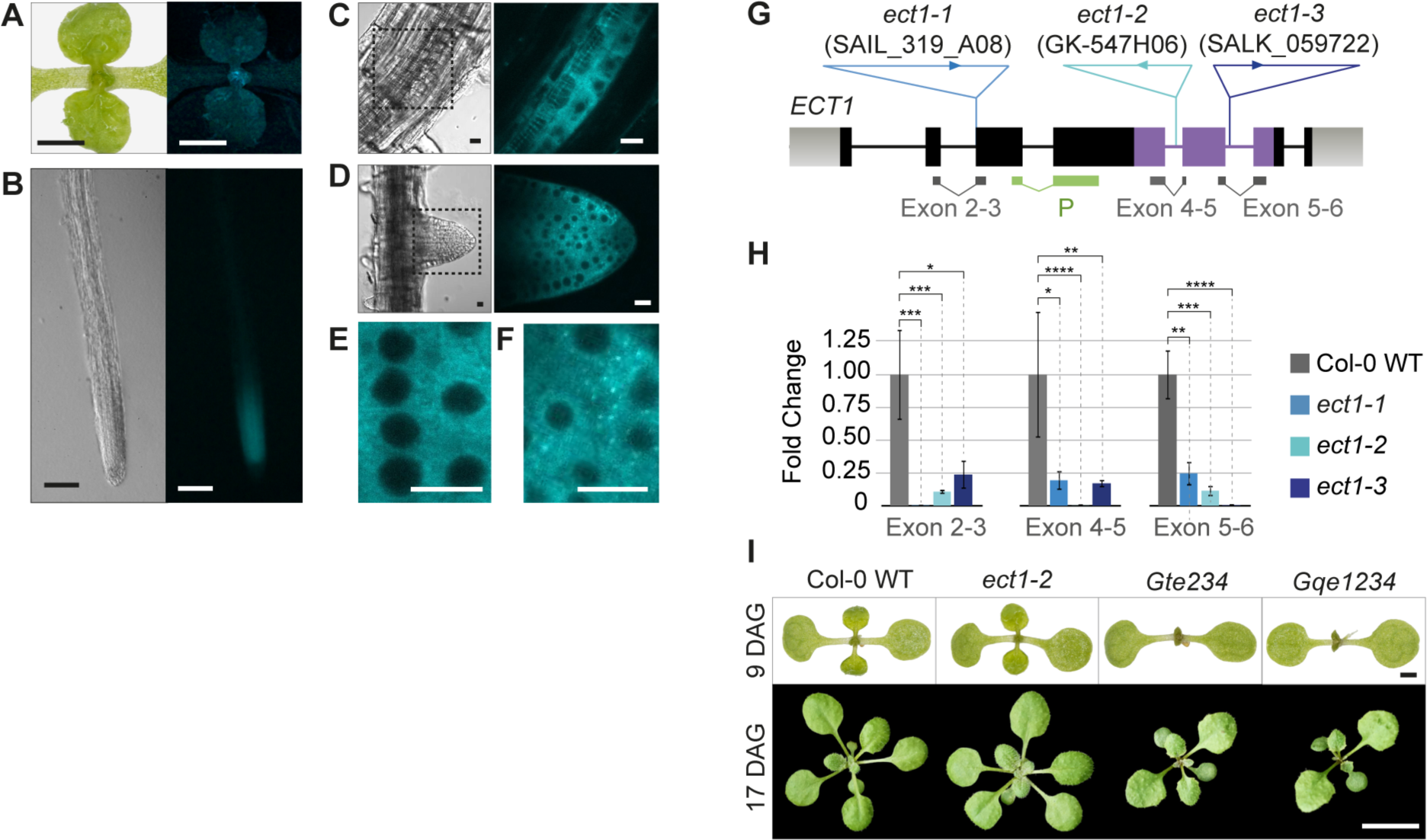
Endogenous ECT1 has negligible ECT2/3/4-like function, despite similar expression pattern and subcellular localization. **(A-D)** Expression pattern of *ECT1p:gECT1-TFP* (gDNA) in aerial organs (A), main root (B), lateral root primordia (C) and emerging lateral roots (D) of 10-day-old seedlings. The expression mimics the pattern observed for ECT2/3/4 (Arribas-Hernández et al., 2020). **(E-F)** Intracellular localization of ECT1-TFP in meristematic cells of root tips in unchallenged conditions. Although the cytoplasmic signal is largely homogenous I, sporadic foci (F) are frequently observed. S13A Fig shows the integrity of the fluorescently tagged protein in independent transgenic lines assessed by protein blot. **(G)** Schematic representation of the *ECT1* locus. Exons are represented as boxes and introns as lines. The positions and identifiers of the T-DNA insertions assigned to the *ect1-1*, *ect1-2* and *ect1-3* alleles are marked, and the location of qPCR amplicons and hybridization probes (P) for analyses is indicated below. Genotyping of *ect* alleles is shown in S13C-F Fig. **(H)** Expression analysis of *ECT1* mRNA in wild type and T-DNA insertion lines by qPCR. *P<0.01, **P<0.001, ***P<0.0001, and ****P<0.00001 in T-test pairwise comparisons (supporting data in S9 Dataset). Northern blot using the probe (P) marked in G detects *ECT1-TFP* mRNA, but the endogenous *ECT1* transcript is below detection limit (S13B Fig). **(I)** Morphological appearance of seedlings with or without *ECT1* in the different backgrounds indicated. Nomenclature and additional phenotyping of these and alternative allele combinations can be found in S14 Fig. DAG, days after germination. Scale bars are: 1 cm in A, 1 mm in B, 100 μm in C, 10 μm in D-F, 1 mm in upper panels of I, and 1 cm in the lower panels.

### Mutation of ECT1 does not exacerbate the phenotype of ect2/ect3/ect4 plants

We isolated three homozygous knockout lines caused by T-DNA insertions in the *ECT1* gene, *ect1-1*, *ect1-2*, and *ect1-3* (Figs 3G,H and S13C-F), and crossed two of them to different *ect2/ect3/ect4* triple mutants [16, 17] to obtain two independent allele combinations of <u>q</u>uadruple *<u>e</u>ct* mutants, *<u>qe</u>1234* and *G<u>qe</u>1234* (see S14A Fig for nomenclature). In both cases, the quadruple mutant plants were indistinguishable from the corresponding *te234* and *Gte234* triple mutant parentals (Figs 3I and S14B). Similarly, triple *ect1/ect2/ect3* mutants were identical to double *ect2/ect3* mutants, and single *ect1* mutants did not show any obvious defects in *ect2/3*-associated traits such as leaf or root organogenesis and trichome branching (Figs 3I and S14B-D). Thus, despite their similar expression pattern, there is no indication of redundancy between ECT1 and the other three *Ath* DF-A paralogs. This unexpected conclusion agrees with the poor ECT2/3/4-like activity of ECT1 in our functional assay, thereby validating the approach.

### The divergent function of ECT1 and ECT11 is caused mainly by properties of their IDRs

Having established that different molecular functions are represented in the arabidopsis YTHDF family, we went on to map the molecular regions responsible for this functional divergence. First, we investigated whether lack of *te234* complementation by ECT1/9/11 is due to altered target binding by the YTH domain, IDR-related effector functions, or a combination of both. Thus, we built chimeric IDR/YTH constructs between ECT2 and ECT1/9/11 (Fig 4A). We also tested an ECT<u>8</u>_IDR_/ECT<u>2</u>_YTH_ <u>c</u>himera (C-8/2) to assess whether hybrid constructs can at all be functional, choosing ECT8 as a positive control due to its robust complementation of seedling and adult phenotypes (Fig 2C-D,F). Expression of the constructs in *te234* mutants showed that the C-8/2 chimera was fully functional (Figs 4B and S7B, S15), thus validating the approach. The ECT1- and ECT11-derived hybrids showed that their YTH domains retain m^6^A-binding ability, because the complementation frequencies for the C-2/1 and C-2/11 chimeras were not zero (Fig 4B), contrary to an m^6^A-binding deficient point mutant of *ECT2* [16]. They also proved, however, that both the IDR and the YTH domain of these two proteins only retain a reduced degree of function compared to the equivalent regions in ECT2. In particular, the IDRs of both proteins scored lower than their YTH domains when combined with the other half from ECT2 (Fig 4B), pointing to divergence in the IDR as the main cause for different functionality. In wild type ECT1 and ECT11 proteins, the combination of the two poorly-performing parts is likely the cause of the minute ECT2/3/4-like activity.

**Fig 4.**
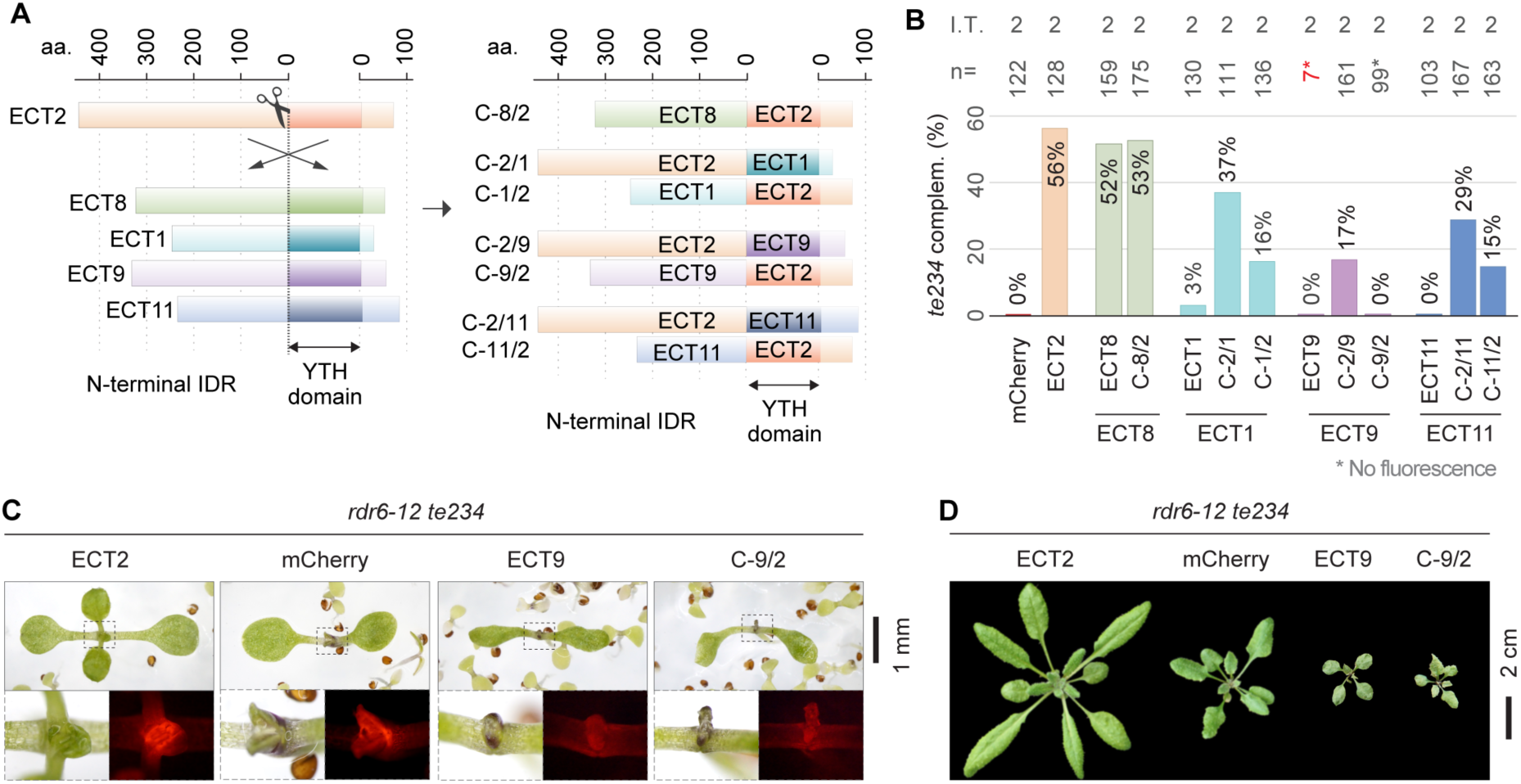
Dissection of ECT1/9/11 functional regions. **(A)** Schematic representation of the strategy followed to express chimeras with N-terminal IDR and YTH domains of different ECT proteins. **(B)** Weighed averages of the complementation rates observed for each chimeric construct measured as in Fig 2B,C. The raw complementation rates and transformation efficiencies of each independent transformation (I.T.) can be found in Figs S7B and S9 respectively, and photographs of the scored 10-day-old T1 seedlings with mCherry fluorescence are shown in S15 Fig. **(C)** 10-day-old primary transformants of *rdr6-12/te234* with the indicated transgenes. Dashed outlines are magnified below to show mCherry fluorescence. **(D)** Same genotypes as in C at 26 days after germination. Additional independent transgenic lines, developmental stages and controls for C and D can be found in S10 Fig. All transgenes are expressed from the *US7Y* promoter and contain mCherry fused to the C-terminus.

### The IDR of ECT9 is incapable of performing ECT2/3/4 molecular functions, but its YTH domain retains some ECT2/3/4-like function

The ECT<u>2</u>_IDR_/ECT<u>9</u>_YTH_ <u>c</u>himera (C-2/9) yielded normal transformation efficiencies and up to 17% of T1s had signs of complementation compared to 56% for ECT2 (Figs 4B and S7B, S9, S15), suggesting that the YTH domain of ECT9 retains at least partial functionality, including m^6^A-binding activity. However, the reverse construct (ECT<u>9</u>_IDR_/ECT<u>2</u>_YTH_, C-9/2) recapitulated the results obtained for ECT9: low transformation efficiency and low fluorescence in the few T1s recovered, and complete silencing in the next generation (Figs 4B and S7B, S9, S15). Hence, we introduced the C-9/2 chimera into the *rdr6-12/te234* background and observed plants with stronger fluorescence exhibiting aberrant morphology, again similar to what was obtained with ECT9 (Figs 4C,D and S7C, S10B). These results confirm that ECT9 is not functionally equivalent to ECT2/3/4, and point to the IDR as the main site of functional divergence of the protein.

### The molecular requirements for m^6^A-binding are conserved in the YTH domains of ECT1/9/11

To pinpoint molecular characteristics of ECT1/9/11 responsible for their different functionality, we performed a series of comparative analyses between ECT1/9/11 and other ECT proteins. We first analyzed the YTH domains, taking ECT2 as a reference and using the structurally well-characterized YTH domain of *Homo sapiens* (*Hs*) YTHDF1 in complex with RNA GGm^6^ACU [61] as a model. Amino acid sequence alignment of the YTH domains of all *Ath* ECTs (S16 Fig), AlphaFold-predictions [74, 75] of the apo structures and homology models [76] of the m^6^A-bound state (S17 Fig) confirmed a high degree of conservation of the key residues required for m^6^A-RNA binding and overall fold of the YTH domain of ECT1/9/11, in agreement with the partial functionality observed in our chimeric constructs (Fig 4B). However, we also detected a few non-conservative amino acid substitutions specific for ECT1/9/11 (Figs 5A and S16), preferentially located on the surface of the protein but far from the RNA-binding groove (S17 Fig). These non-conservative substitutions are prime candidates to explain the minor differences in functionality, a scenario that implies biological functions of YTH domains other than mere m^6^A-binding capacity.

**Fig 5.**
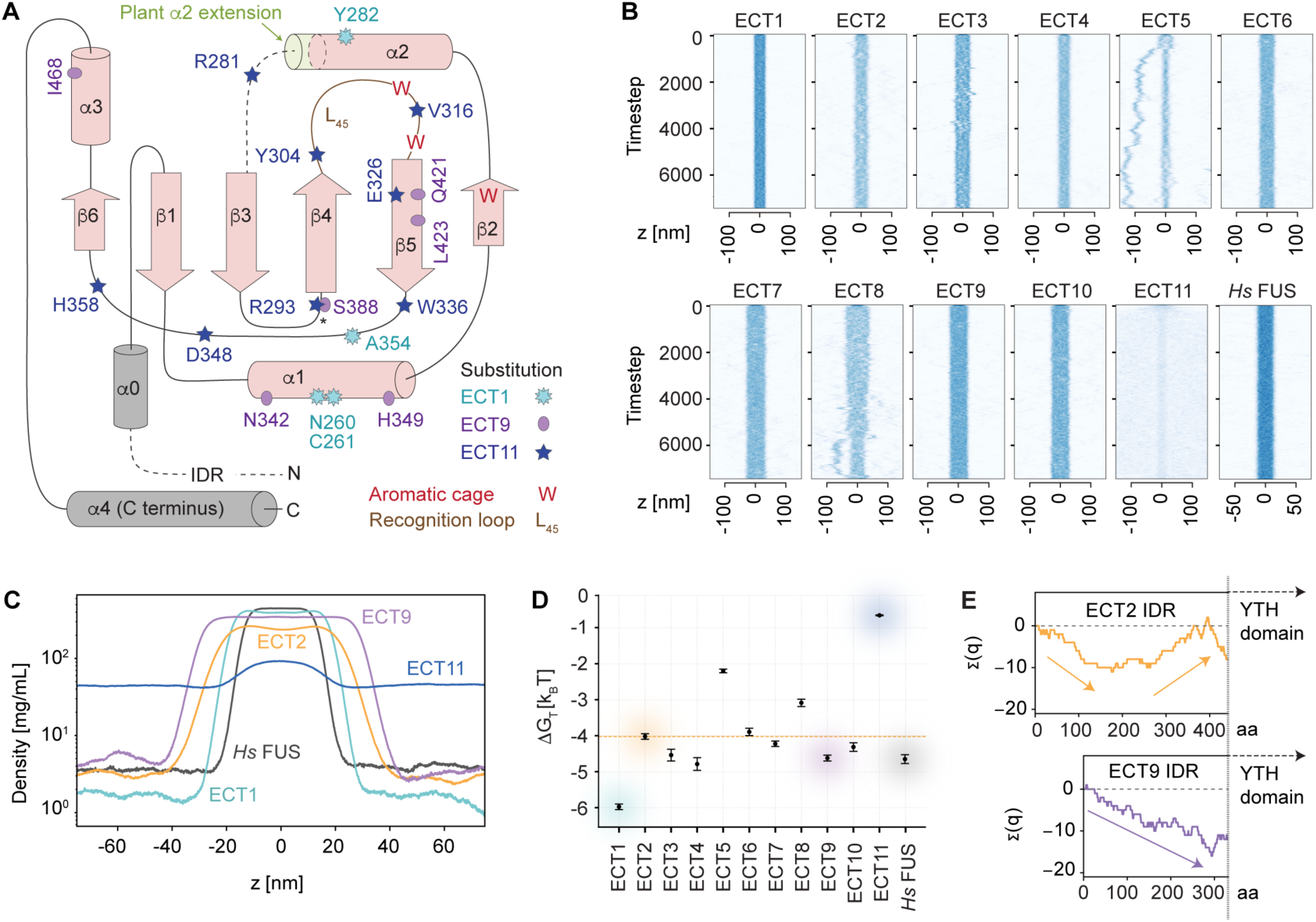
Analysis of ECT1/9/11 functional regions. **(A)** Topology of the YTH domain of *Homo sapiens* (*Hs*) YTHDF1 with secondary structure elements drawn in pink [61]. The two grey helices represent the only structural elements outside the YTH domain as defined by Stoilov et al. [8]: α0 immediately upstream, and α4 at the C-terminus of the protein. Superimposed on the diagrams are the non-conservative amino acid substitutions found in *Ath* ECT1, ECT9 and ECT11 compared to the majority of *Ath* ECTs (see alignment in S16 Fig). The localization of these amino acids in the predicted 3D structures, predominantly on the surface, and the spatial integrity of the aromatic cage in the homology models of ECT1/9/11, can be found in S17 Fig. **(B)** Slab simulations of the IDRs of the indicated proteins showing their tendency to remain in condensed phase or disperse in solution. Source videos of ECT2, ECT1 and ECT11 simulations are provided as S1-S3 Movies **(C)** Time-averaged density profiles for a subset of the proteins in B. The analyses of the remaining *Ath* ECTs are shown in S19A Fig. **(D)** Calculated free energy change for the transition between aqueous solution and condensed phase (ΔGtrans = RT ln [cdilute / cdense]) for the IDRs of the indicated proteins. **(E)** Cumulative charge distribution of the IDRs of the indicated proteins. The length in amino acids (aa) is at scale. The same analysis for all *Ath* ECTs can be found in S19D Fig.

### The IDRs of ECT1/9/11 have biophysical properties distinct from ECT2/3/4/5/6/7/8/10

We next focused on the N-terminal regions of *Ath* ECT1/9/11. Because protein disorder predictions using MobiDB [77] confirmed that they are all disordered (S18 Fig), we used approaches suitable for the study of IDRs. Prediction of short linear motifs (SLiMs) that may mediate interaction with effector proteins gave many possibilities that did not follow a clear pattern when compared between the different ECT classes (S18 Fig). Hence, we decided to study other properties known to be important in IDRs and focused on the propensity to form liquid condensates, the amino acid composition, and the distribution of charge. To determine whether the distinct functional classes of ECTs differed in their propensity to phase separate, we employed a recently developed coarse-grained model of IDRs with residue-specific parameters estimated from experimental observations of many IDRs [78] to simulate their tendency to remain in the condensed phase or disperse in solution. As control, we used the well-studied IDR of the human RNA-binding protein Fused in Sarcoma (FUS) [79–82]. The results showed that the IDR of ECT2 has a clear propensity to remain in the condensed phase (Fig 5B-C and S1 Movie) with a highly favorable Gibb’s free energy change for the transition from dilute aqueous solution to condensed phase (Fig 5D), comparable to, albeit not as pronounced as for the IDR of *Hs* FUS (Fig 5B-D). While the IDRs of most *Ath* ECT proteins behaved comparably to ECT2, the IDRs of ECT1 and ECT11 were clear outliers (Figs 5B-D and S19A). Compared to all other ECTs, the IDR of ECT1 showed a much stronger propensity to phase separate, consistent with the spontaneous granule formation of ECT1-TFP *in vivo* (Figs 3F and 5B-D, and S2 Movie). On the contrary, the IDR of ECT11 dispersed into solution much more readily (Fig 5B-D and S3 Movie). Thus, the IDRs of ECT1 and ECT11 have markedly different biophysical properties from those of ECT2/3/4/5/6/7/8/10, perhaps contributing to their different functionality *in vivo*. In contrast, the IDR of ECT9 did not stand out from the rest in the simulations of phase separation propensity. We also note that the different phase separation propensity among the ECTs is driven, in the main, by their different content of sticky residues as defined by the CALVADOS model [78] (S19B Fig). In this context, it is potentially of interest that ECT1, ECT9 and ECT11 all have a higher content of Phe residues than the nearly Phe-depleted IDRs of ECT2/3/4/5/6/7/8/10 (S19C Fig). For ECT1, whose Tyr and Trp enrichment is comparable to that of ECT2/3/4/5/6/7/8/10 (S19C Fig), the additional enrichment of the similarly sticky Phe may contribute to its more pronounced phase separation propensity.

We finally analyzed the charge distribution, as this may dictate properties such as compactness and shape of IDR conformations [83, 84]. We found that the IDRs of ECT2/3/4/5/6/7/8/10 show a recurrent pattern with accumulation of net negative charge towards the N-terminus, and recovery to near-neutrality closer to the YTH domain, resulting in creation of electric dipoles (Figs 5E and S19D). The IDRs of ECT1, ECT9 and ECT11 differ from this pattern, but in distinct ways: ECT1 does not show a pronounced patterning of the charged residues, while ECT9 increasingly accumulates negative charge (Fig 5E) and ECT11 increasingly accumulates positive charge (S19D Fig). It is an interesting possibility, therefore, that charge separation in the IDR is a requirement to fulfill the molecular functions of ECTs to accelerate growth in primordial cells.

### The ability to complement ect2/3/4 does not simply follow affiliation with phylogenetic clades

We next compared the functional groups of YTHDF proteins defined by our complementation assays with the YTHDF phylogeny. Strikingly, the ability to rescue defective timing of leaf emergence does not follow the defined phylogenetic clades, because both high and low/null complementation capacities are present in the same clades. For example, ECT2, ECT5 and ECT8 in clades DF-A, -B and -C respectively present high ECT2/3/4-like functionality, while ECT1, ECT9 and ECT11 in these same clades do not (Figs 1B and 2C-H). It is noteworthy, however, that there is a clear trend for non-complementors to be amongst the most highly divergent proteins in each group, as revealed by the length of the branches in the phylogenetic tree (Fig 1B). Indeed, some of the longest branches in both our tree and the one in the study by Scutenaire et al. [26] correspond to ECT1, ECT9, and ECT11. Accordingly, the three proteins have the highest number of amino acid substitutions compared to the only YTHDF protein of the liverwort *Marchantia polymorpha* (*Mpo* DFE) [85] (Fig 6A). These results suggest that the common ancestor of YTHDF proteins in land plants may have had a molecular function related to that of the modern *Ath* ECT2/3/4, and that this function has been conserved in the least-divergent members of the DF-A/B/C/D clades. Subsequent duplication and neofunctionalization may have driven the rapid evolution of highly-divergent YTHDF proteins (ECT11, ECT1, and ECT9 in arabidopsis) that have lost their primary function and perhaps fulfill different molecular roles in the plant m^6^A pathway.

**Fig 6.**
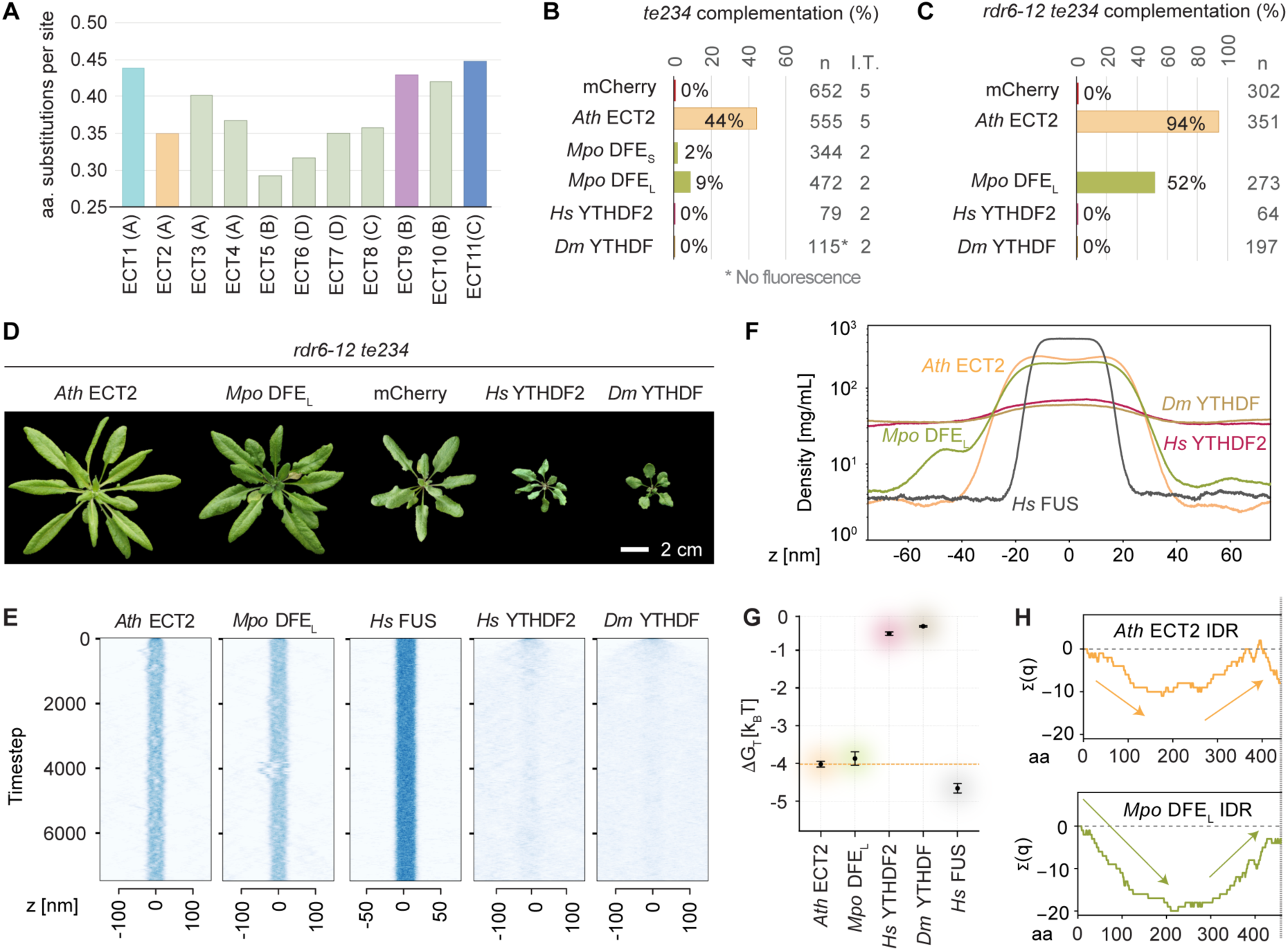
Heterologous expression of a bryophyte DFE rescues loss of ECT2/3/4 function in arabidopsis, while metazoan YTHDF proteins aggravates it. **(A)** Evolutionary divergence of *Arabidopsis thaliana* (*Ath*) ECT proteins estimated by pairwise comparisons between these and the only DF protein in *Marchantia polymorpha* (*Mpo* DFE) according to the alignment of YTHDFs from bryophytes and angiosperms in S7-S8 Datasets (see Methods). **(B-C)** Weighed averages of the complementation rates observed for *Mpo* DFE (S/L refer to the <u>S</u>hort and <u>L</u>ong splice forms found in meristematic tissues–apical notches and gemma cups–of *M. polymorpha* thalli), *Homo sapiens* (*Hs*) YTHDF2 and *Drosophila melanogaster* (*Dm*) YTHDF in the *te234* (B) or *rdr6-12/te234* (C) backgrounds, calculated as in Fig 2C-D. n, total number of transformants; I.T. number of independent transformations (only one in C). Raw T1 counts, complementation rates and transformation efficiencies can be found in the S6 Dataset, and S7 and S9 Figs. **(D)** 32-day-old primary transformants expressing the transgenes analysed in C. Additional controls and developmental stages are shown in S10B Fig. **(E)** Slab simulations of the IDRs of the indicated proteins as in Fig 5B. **(F)** Time-averaged density profiles of the slab simulations in E. **(G)** Calculated free energy change for the IDRs of the indicated proteins as in Fig 5D. **(H)** Cumulative charge distribution of the IDRs of the indicated proteins, as in Fig 5E. The same analysis for the two metazoan YTHDFs can be found in S19D Fig.

### The molecular function of ECT2/3/4 was present in the first land plants

To test whether the common ancestor of plant YTHDF proteins indeed had a molecular function similar to that of the modern *Ath* ECT2/3/4, we subjected *Mpo* DFE (Fig 2A) to our functional assay, as bryophytes are the earliest-diverging taxon of land plants (Fig 1A, [54, 56–60]). The result showed a partial, but clear capacity of two different splice forms of *Mpo* DFE to complement the delay in leaf emergence and morphology defects of arabidopsis *te234* plants (Fig 6B and S7D, S20A-B). Although the complementation scores were low (Fig 6B), a thorough inspection of the fluorescence intensity in all primary transformants revealed that the transgene was only expressed very weakly in a reduced subpopulation of plants, possibly due to the tendency to trigger silencing upon heterologous expression of a long cDNA from a distant plant species [86]. We therefore introduced the best-complementing isoform into *rdr6-12/te234* mutants. This approach yielded a much higher complementation score of 52% (Fig 6C), and plants very similar to those expressing ECT2 in the same background (Fig 6D and S6B). Thus, the *Ath* ECT2/3/4 activity needed to stimulate proliferation of primed stem cells was present in the first embryophytes, suggesting that this activity is an ancestral molecular function of YTHDF proteins in land plants. Gratifyingly, the simulations of the behavior of the IDR of *Mpo* DFE showed a dipole-like charge pattern and a phase separation propensity similar to those of the ECT2/3/4/5/6/7/8/10 group, further substantiating that these IDR characteristics correlate with the ability to fulfill ECT2/3/4 function in leaf formation (Fig 6E-H). We investigated this interesting possibility more deeply by analysing the IDRs of YTHDF proteins from species representing the main clades of land plants that diverged prior to flower evolution, a moss, a lycophyte, a fern and a gymnosperm, and from the sole species in the sister group to all flowering plants, *A. trichopoda* [87, 88]. This analysis showed that at least one YTHDF protein with ECT2-like charge distribution and phase separation propensity is present in all these taxa (S21A Fig). In particular, this included the only YTHDF protein in the moss *Ceratodum purpureus*. Interestingly, when the proteins from the different taxa were analysed together, a general trend for increased phase separation propensity that again correlated with higher ‘stickiness’ (λ) was apparent in proteins of clades -A, -B and -E compared to those of -C, -D and -F (S21B Fig). We also noticed that in the species that encode multiple YTHDF proteins, some paralogs exhibit divergent IDR features similar to those seen in *Ath* ECT1/9/11, but this phenomenon was not clade-specific (S21 Fig). This result suggests that the IDR-based functional diversification observed in arabidopsis may apply more generally. We conclude that the molecular function of YTHDF proteins required for stimulation of primordial cell proliferation is deeply conserved in land plants, and that the biophysical properties of the IDRs of *Mpo* DFE and the *Ath* DF proteins capable of performing this function are recurrent in all the major plant taxa.

### Heterologous expression of animal YTHDFs enhances the phenotype caused by loss of ECT2/3/4 in Arabidopsis

Finally, we subjected *Hs* YTHDF2 and *Drosophila melanogaster* (*Dm*) YTHDF (Fig 2A) to our functional assay to test whether they could function molecularly like plant YTHDFs. While we did not observe complementation of the delayed leaf emergence of *te234* plants in any of the transformants (Figs 6B and S7D), several *Hs* YTHDF2 lines showed more acute developmental defects compared to *te234* (S20B Fig). Although these defects were not observed among *Dm* YTHDF lines, no expression could be detected in these plants either (S7D Fig). Hence, we repeated the transformation in the *rdr6-12/te234* background, this time obtaining numerous transformants of both *Dm* YTHDF and *Hs* YTHDF2 with strong fluorescence (Figs 6C and S10B). These plants showed aberrant morphology comparable to the effect caused by expression of ECT9 and C-9/2, with resemblance to arabidopsis mutants with severe depletion of m^6^A [51, 52] (Figs 6D and S10B). Thus, expression of metazoan YTHDFs in arabidopsis *ECT2/3/4*-deficient plants exacerbates their developmental phenotype. This result is in agreement with the generally accepted idea that m^6^A-YTHDF2 destabilizes mRNA in mammals [9, 89–92] while the m^6^A/ECT2/3/4 axis does the opposite in plants [23, 51, 93–97]. However, an analogous molecular mechanism between plant and animal YTHDFs cannot be completely ruled out, because a different cellular context may be hampering the molecular activity of metazoan YTHDFs in plant cells, and simple competition for targets with remaining ECTs could be the cause of the enhanced phenotype. Nonetheless, we note that the IDRs of *Hs* YTHDF2 and *Dm* YTHDF had much weaker phase separation propensity (Fig 6E-G) and different charge distribution (S19D Fig) compared to ECT2, perhaps contributing to the lack of ECT2/3-like function *in planta*.

## Discussion

Our study combines thorough phylogenetic analyses with functional complementation to assess the extent to which molecular functions of YTHDF proteins have diverged over the course of land plant evolution. The key observation is that 8 of the 11 *Ath* ECTs and, remarkably, the sole YTHDF of the liverwort *M. polymorpha*, have the molecular functions required for leaf development in arabidopsis. Along with the refined phylogenetic relationships between plant YTHDFs that we present, these findings have implications for our understanding of three distinct, if obviously related, areas that we discuss in more detail below: (1) the molecular properties of land plant YTHDF proteins, (2) the ancestral YTHDF functions and the evolutionary paths towards YTHDF diversification, and (3) the biology underlying the recurrent employment of expanded YTHDF families in higher plants.

### 1. Inferences on molecular properties of land plant YTHDF proteins

Our study provides at least four new insights into molecular properties of land plants YTHDF proteins, all of which are important for future, more detailed mechanistic studies of eukaryotic m^6^A-YTHDF function. First, the functional assays with *Ath* ECT proteins and the chimeras combining the YTH domain of ECT1/9/11 with the IDR of ECT2 allow us to infer that all 11 arabidopsis YTHDFs have m^6^A-binding capacity, and suggest that some, if not all, of their *in vivo* functions involve m^6^A binding. Second, the complementation analyses of the reverse chimeras show that the IDRs of *Ath* ECTs with divergent functions contribute substantially to their different molecular properties. This finding implies a key role of IDRs in defining the molecular activities of YTHDF proteins, and is in line with the conclusions of a recent study of mammalian YTHDFs [48], with the functional importance of the IDR of *Hs* YTHDF2 inferred from early tethering studies [9], and with the isolation of IDR-dependent interactors proposed to be important for *Hs* YTHDF2 and *Ath* ECT2 function [28–30, 96, 98]. Third, the ability to direct correct leaf formation correlates with a combination of two biophysical properties of the IDRs of plant YTHDFs: a clear propensity to phase separate, and an organization of charged residues that creates an electric dipole. Future studies are needed to address whether a causal relation underlies this correlation, and, if so, what its molecular basis is. Fourth, amino acid substitutions in YTH domains inferred to be capable of binding to m^6^A also contribute, to a lower extent, to functional specialization. Because most affected amino acids are surface-exposed but not predicted to contact RNA, this observation indicates that YTH domains should not be seen as inert adaptors for m^6^A association, as they may also participate in effector functions. It is an important goal of future studies to identify such functions and define how they contribute to functional diversification.

### 2. The ancestral function and evolution of YTHDF proteins

It is a key finding of our study that the sole *M. polymorpha* YTHDF protein possesses the molecular functions necessary for leaf formation in flowering plants. This remarkable result demonstrates deep functional conservation, across at least 450 My of evolution, and suggests that stimulating cell division in organ primordia may be the ancestral function of land plant YTHDFs. It is now a question of profound interest to determine how the molecular activity of the ancient YTHDF proteins evolved and diversified as the main eukaryotic linages came into existence. Our work provides several important new insights to guide the understanding of this process. In particular, since it is a valid question whether the main evolutionary event that gave rise to the ancestral function of land plant YTHDFs occurred prior to their terrestrialization, we open this section of the discussion with considerations on how the available information on algal YTHDF proteins informs the debate. We close with the analysis of *Ath* ECT1 divergence as a case study on evolution of specialized YTHDF function.

### *2a.* YTHDF proteins in chlorophyte algae are either lost or highly divergent

The genomes of most of the chlorophyte species sequenced thus far do not contain *YTHDF* genes. Indeed, the single YTH domain protein in the model green alga C*hlamydomonas reinhardtii* belongs to the DC clade [26, 68], and the same is true for all the species that we inspected in this and five additional orders (S2 Fig). We only found YTHDF-encoding genes in a subset of species in the basal group Mamiellophyceae (S2 Fig). Like land plant, metazoan and yeast YTHDF proteins, these chlorophyte YTHDFs have long N-terminal IDRs and no folded domains other than the YTH domain. Nevertheless, YTHDFs have diverged remarkably fast in Chlorophyta, as revealed by their long branches in the phylogenetic tree, well surpassing that of YTHDF proteins in streptophytes (land plants and charophyte algae) and even the microalgal species of Glaucophyta, Rhizaria and Haptophyta used as outgroup [99, 100] (left bottom corner in Fig 1B, and S1 Fig). Furthermore, the amino acid substitutions and insertions found in some of the proteins affect the aromatic cage (S3 Fig), likely compromising their m^6^A-binding capacity. Hence, YTHDF proteins are poorly represented in chlorophytes and, as for the only YTH-domain protein of fission yeast Mmi1 [69], they may have lost m^6^A-related functions altogether.

### *2b.* YTHDF proteins are widespread and variably divergent in the sister clade of land plants

Another informative group of green algae is the charophytes, or streptophyte algae, photosynthetic organisms that thrive in fresh water or humid terrestrial habitats. Their ancestors split from chlorophytes over a billion years ago and gave rise to all land plants and extant charophytes, [55, 62, 101]. The genomes of most charophyte species sequenced thus far encode YTHDF proteins (Fig S2) with intact aromatic cages and overall higher conservation than those of chlorophytes (Fig S3), but some of them also differ considerably from their closest land plant relatives (Figs 1B and S1, S3), and may include additional folded and even transmembrane domains (S22 Fig).

### *2c.* Possible evolutionary scenarios

In the light of the observations regarding YTHDFs in green algae, we consider three possible scenarios for the acquisition of ECT2-like molecular functions in land plants:

1. The acquisition occurred during the terrestrialization of the common ancestor of all land plants or, at the latest, within the time window between terrestrialization and bryophyte divergence. In that case, charophyte YTHDFs should not exhibit ECT2-like functionality.
2. The acquisition occurred early during streptophyte algae evolution. We consider this the most likely scenario, because such acquisition could have served as exaptation (pre-adaptation) for the successful colonization of dry land, perhaps by contributing to the appearance of apical meristems [55, 102]. The prediction in that case would be that charophytes from late-diverging clades, but not from basal groups, encode YTHDFs with ECT2-like functions.
3. The key evolutionary event occurred much earlier, perhaps upon chloroplast acquisition by the last common ancestor of Viridiplantae, Rhodophyta and Glaucophyta, i.e. the first Archaeoplastida, or even before that. We have, however, not been able to find any YTH domain protein in rhodophytes (red algae), and the only glaucophyte sequenced so far [103] contains a single YTHDF (S2 Fig). These observations, together with the scarcity and great divergence of chlorophyte YTHDFs, suggest that if the functional acquisition indeed happened in ancestral Archaeoplastida or earlier, the function and even the proteins altogether were lost in most green and red algae, but kept in the streptophyte lineage that gave rise to land plants. Future functional studies of YTHDFs from Charophyta and Glaucophyta, and taxa close to Archaeoplastida such as Haptophyta, Rhizaria or Stramenopiles (S2 Fig, [99, 100]) will be key for the dissection of the evolutionary origin of the ECT2-like molecular properties required for plant growth [104].

### *2d.* ECT1: a newly evolved gene in a Brassicaceae lineage as a case study

In addition to the fundamental question on evolution of the ancestral molecular function of land plant YTHDFs, it is also of interest to consider what characterizes the evolution of late YTHDF specialization. We examine *Ath* ECT1 as a case study in this regard. A thorough look at phylogenetic analyses from Scutenaire *et al.* [26] reveals that *ECT1* is present only in a few species of Brassicaceae such as *A. thaliana* and *Capsella rubella* (*Cru*). *Brassica oleracea* in the same family has a clear homolog of *Ath/Cru ECT3* but not of *ECT1*, other dicots encode *ECT3/1* orthologs in a separate cluster, and monocot DF-As arose from an entirely separate branch (S1 Fig, [26]). Therefore, *ECT1* is the product of a recent duplication of the *ECT3/1* gene in a small subgroup of dicots, that diverged more rapidly than any other DF-A clade member in arabidopsis (Fig 1B, [26]). This observation agrees with our overall conclusion that most YTHDFs in early angiosperms possessed *Ath* ECT2-like molecular functions, and only late events resulted in neofunctionalization of specific paralogs. We note that such rapid divergence appears to result primarily from changes in the IDR. We were able to identify two biophysical properties of the IDR of ECT1—its stronger predicted propensity to phase separate and its charge distribution—that are distinct from the ECT2/3/4/5/6/7/8/10 group, but note that additional changes, e.g. in SLiMs mediating interaction with regulatory factors, may also contribute to the different functionality [98].

### 3. Functional versus biological redundancy and the need for many YTHDFs

Our study reveals that most YTHDF proteins in higher plants can perform, essentially, the same molecular functions that allow ECT2/3/4 to promote cell proliferation in organ primordia and control organogenesis. We anticipate that this molecular function may be used in distinct biological contexts by ECT5/6/7/8/10 due to different expression patterns and response to stimuli, thereby achieving biological specialization. Such variability in expression patterns can be inferred from available RNA-Seq data in the arabidopsis eFP Browser (BAR.toronto.ca) that show, for example, that ECT8 is primarily expressed during senescence-associated processes, perhaps in connection to re-growth after reversible proliferative arrest [105, 106]. Clearly, however, higher plants do not only use many YTHDF-encoding genes to provide the same molecular function(s) in distinct expression domains, as shown by the specialized molecular activity of *Ath* ECT1, ECT11, and most prominently ECT9. Whether these divergent proteins act to counteract the ECT2/3/4-like function by competing for m^6^A-containing transcripts, perhaps halting growth upon stress, or whether they perform entirely independent functions in different tissues or cell types, is open for new investigations.

An outstanding question not yet resolved by our study is why higher plants seem to need many YTHDF proteins. Indeed, the number of paralogs in vascular plants well surpasses the three YTHDFs present in most vertebrates [6] and the even smaller sets in other eukaryotes (S2 Fig, [8, 68]). The need for many YTHDFs of considerable variability is all the more puzzling given our demonstration here that the majority of them retains the same molecular functions. Robustness is a commonly given explanation for redundancy, but the same argument could apply to YTHDCs, typically encoded by 0-2 genes in most eukaryotes including land plants [8, 68] (S2 Fig), or methyltransferase subunits, for which only single copies have been maintained during evolution [68]. Therefore, the answer should probably be sought elsewhere. It is clear now that the m^6^A/YTHDF axis controls cellular proliferation and organ growth in plants [17, 97], and a major difference between land plants and other complex organisms in that regard is the extreme plasticity of plant development. Because growth and stress responses are also tightly intertwined, it is possible that plants use a plethora m^6^A-YTHDF growth-stimulators in different organs, responsive to different stresses and developmental cues, to shape plant architecture in response to stimuli. The answer to the paradox may be in the details: plant YTHDFs responsive to different environmental cues may fine-tune growth programs through slightly different performance and target-binding capacity in partly overlapping expression domains, where they are known to compete for targets [23].

## Materials and Methods

### Plant material and growth conditions

All the lines used in this study are in the *A. thaliana* Columbia-0 (Col-0) ecotype. The T-DNA insertion lines *rdr6-12* [72], *ect1-1* (SAIL_319_A08), *ect1-2* (GK-547H06), and *ect1-3* (SALK_059722) were obtained from Nottingham Arabidopsis Stock Centre. The mutants *ect2-3*, *ect3-2*, *ect2-3/ect3-2* (*Gde23*), *ect2-3/ect3-2/ect4-2* (*Gte234*) and *ect2-1/ect3-1/ect4-2* (*te234*) used for genetic crosses or as background to produce transgenic lines, and the primers for genotyping the corresponding alleles, have been previously described [16, 17]. The standard growing conditions used are detailed in S1 Appendix.

### Phylogenetic analysis

Amino acid sequences of all YTHDF proteins found in 40 species distributed among the main taxa of Viridiplantae were downloaded from UniProt (Apweiler et al., 2004), TAIR (www.arabidopsis.org), Phytozome [107], GinkgoDB [108], FernBase [109] and PhycoCosm [110, 111] (S1 Dataset). As outgroup, we selected three species from the eukaryotic taxa Glaucophyta (*Cyanophora paradoxa, Cpa*), Haptophyta (*Emiliania huxleyi, Ehu*) and Rhizaria (*Paulinella microscpora, Pmi*), as these are evolutionarily closer to Viridiplantae than e.g. Opisthokonta such as animals and fungi [99, 100]. All sequences were aligned with MUSCLE [112, 113] using default parameters (S2 Dataset). The alignment was trimmed from both ends to discard regions with poor sequence conservation according to the quality and conservation scores provided by Jalview [114] (S3 Dataset and S23 Fig). The retained part of the alignment, comparable to the region used for phylogenetic analysis in a previous study [26], contains the entire canonical YTH domain [8] and, on both sides, short stretches with high conservation scores (S23 Fig). Evolutionary analyses were conducted using the maximum likelihood algorithm IQ-Tree 2 [115, 116] from the CIPRES Science Gateway v. 3.3 (https://www.phylo.org/index.php/) to infer a phylogenetic tree. The model based on Q matrix [117] estimated for plants [118], ‘Q.plant+R8’, was determined as best-fit using the built-in tool ModelFinder [119] according to Bayesian Information Criterion (BIC) scores, and used for the analysis. Measures of branch supports by approximate Likelihood Ratio Test [aLRT] (SH-aLRT, %) [120] were also calculated. The resulting tree (S4 Dataset) was rooted on *Pmi* YTHDF, annotated adding SH-aLRT values, and color-coded using FigTree (https://github.com/rambaut/figtree/). An additional alignment performed with a subset of these proteins with the purpose to illustrate the conservation of the YTH domain in green algae and non-flowering plants was calculated and trimmed as described above (S5 Dataset). The graphical representations of this alignment (S3 and S4 Figs) were obtained using Jalview [114].

Estimates of evolutionary divergence between *A. thaliana* and *M. polymorpha* YTHDF proteins were obtained computing pairwise distances from the full length alignment of homologs in a subset of land plant species (bryophytes, basal angiosperms, magnoliids and dicots) (S6 Dataset). These taxa were chosen based on the observation that inclusion of divergent proteins from green algae, ferns, gymnosperms and monocots causes loss of information due to more frequent gaps, while proteins from species closely related to *M. polymorpha* (bryophytes) and *A. thaliana* (dicots and basal angiosperms) maximize the accuracy of the alignment. The analysis was done as described above, and we used the Equal Input model [121] in MEGA11 [122] to calculate the distances. All positions containing gaps and missing data were eliminated (complete deletion option), resulting a total of 171 positions in the final dataset. The complete matrix as well can be found in S7 Dataset.

### Cloning, plant transformation and line selection

We employed the scar-free USER cloning method [123] to produce *US7Yp:cYTHDF(X)-mCherry-OCSter* and *ECT1p:gECT1-FLAG-TFP-ECT1ter* constructs by gluing PCR-amplified DNA fragments into the pCAMBIA3300-U plasmid [124] that provides resistance to glufosinate in plants. Cloning of the trial *ECT2p:gECT1-mCherry-ECT2ter* and *ECT2p:gECT4-mCherry-ECT2ter* constructs was done with GreenGate [125]. The detailed cloning, transformation and line selection strategy for each construct, as well as the specific details of T1 selections for the complementation assay, are explained in S1 Appendix.

### IDR modeling and simulations

We carried out coarse-grained simulations of protein IDRs using the implicit-solvent CALVADOS 2 forcefield [78]. In CALVADOS, amino-acid residues are mapped to single beads corresponding to their amino-acid type. Neighboring residues are linked using harmonic bonds with an equilibrium distance of 0.38 nm and force constant of 8033 kJ mol^-1^nm^-2^. Nonbonded interactions are modeled with a hydropathy amino-acid “stickiness” model using 𝜆-parameters from CALVADOS 2 [78, 126]. Salt-screened electrostatic interactions are accounted for with a truncated (r_c_ = 4 nm) and shifted Debye-Hückel potential; non-electrostatic interactions were truncated at 2 nm [126]. We used a Langevin integrator with friction coefficient 𝛾 = 0.01 ps^-1^ to propagate the simulation in 0.01 ps timesteps. All simulations were carried out at T = 293 K.

We performed slab simulations of IDRs of the following proteins: *Ath* ECT1-11, *Dm* YTHDF1, *Hs* YTHDF2 and FUS, *Mpo* DFE, *Cpu* DFE, *Ita* DFE1-3, *Cri* DFF1-3, DFAB and DFD1-2, *Gbi* DFF1-2, DFA1-2, DFB, DFC1-2 and DFD, and *Atr* DFA1, DFB, DFC1-2 and DFD, using a previously described procedure [126, 127]. IDR sequences of all proteins are detailed in S1 Dataset. Briefly, we simulated 100 copies of each IDR (one type of IDR at a time) in a simulation slab of dimensions 17×17×300 nm for *Ath* ECT1/3/6-11, *Gbi* DFB and DFF1 and *Atr* DFC2, 15×15×150 nm for *Hs* FUS, or 25×25×300 nm for all other proteins, accounting for the different IDR sequence lengths. We simulated each system for 7500 ns and recorded configurations at 1-ns intervals. The first 2000 ns of each simulation were considered as equilibration and not used for analysis. We centered the protein condensates in the z-dimension of the simulation box during post-processing. The concentrations of the dense phase and dilute phase were calculated by fitting the concentration profiles to a hyperbolic function, as described by Tesei *et al.* [78]. Excess transfer free energies were calculated from the dilute and dense phase equilibrium concentrations via ΔG_trans_ = RT ln (c_dilute_ / c_dense_) [128]. All simulation data and analysis scripts are available at the Zenodo repository (https://doi.org/10.5281/zenodo.8269719).

Charges (q) were added along the IDR sequences to obtain cumulative charge plots. Arg and Lys residues were assigned charges of q = +1; Glu and Asp residues were assigned charges of q = -1; other residue types, including His, were assigned charges of q = 0. The average ‘stickiness’ of a given IDR sequence was determined by calculating the mean hydrophobicity 𝜆̅ of the IDR amino acid composition, using the CALVADOS 2 hydrophobicity scale reported in [126].

### Additional methods

S1 Appendix describes additional standard methods including DNA and cDNA obtention, genotyping (see S13C-F Fig and S2 Table for additional details), fluorescence microscopy, qPCR (see S9 Dataset for the raw data), RNA and protein blotting, and the characterization of YTH domains.

## Acknowledgements

We thank Lena Bjørn Johansson, Sergio D’Anna, Jakub Najbar and Tobias Lahti for technical assistance, Theo Bølsterli and his team for plant care, and the Danish National Life Science Supercomputing Center Computerome and the core facility for biocomputing at the Department of Biology (University of Copenhagen) for the use of computational resources. We acknowledge the constructive criticism of Rupert Fray as a reviewer on previous publications, as his suggestion to use the *ect2/ect3/ect4* triple mutant phenotype to test ECT molecular function by systematic ectopic expression in the ECT2 expression domain inspired us to embark on the present project.

## Supporting information captions

**S1 Appendix. Extended Materials and Methods.**

**S1 Table. Isoforms contained in cYTHDF constructs, and templates used for PCR**.

**S2 Table. Constructs and DNA Oligonucleotides.** All sequences are 5’ to 3’.

**S1 Dataset. Protein Sequences.** Amino acid sequences of proteins used in this study with their identifiers and taxonomic classification. Abbreviations of species names match with those used in Figs 1 and S1. For YTHDF proteins, the canonical YTH domain is colored in blue, and long internal insertions within the YTH domain are highlighted in a paler shade of blue. Additional folded domains are marked in other colors. The sequence of the IDR used for biophysical simulations of some of the proteins is the entire sequence upstream to the YTH domain. For the human fused in sarcoma (FUS) RNA-binding protein used as control for the analyses of IDR properties, the IDR sequence is highlighted in bold.

**S2 Dataset. Full length alignment of Viridiplantae YTHDF proteins.** Full-length amino acid sequence alignment of the YTHDF proteins used for the phylogenetic analyses in Figs 1 and S1. The alignment was obtained using MUSCLE (Multiple Sequence Comparison by Log-Expectation [112, 113]), and is given in clustal format.

**S3 Dataset. Trimmed alignment of Viridiplantae YTHDF proteins.** Amino acid sequence alignment of the YTHDF proteins used for the phylogenetic analyses in Figs 1 and S1 after trimming the poorly conserved regions (IDRs) at both ends, according to the score given by Jalview [114] (S23 Fig), from the alignment provided in S2 Dataset.

**S4 Dataset. Phylogenetic tree of Viridiplantae YTHDF proteins.** Phylogenetic tree (Maximum likelihood phylogenetic analysis) in Nexus/Newick format obtained from the trimmed alignment provided in S3 Dataset. Values of nodes are SH-aLRT (%) support.

**S5 Dataset. Full length alignment of YTHDF proteins in Viridiplantae focused on early-diverging clades**. Full-length amino acid sequence alignment of selected YTHDF proteins, with focus on early-diverging taxa within Viridiplantae, used for S3 and S4 Figs. The names of the proteins are preceded by a ‘tag’ related to the DF clade and organism taxon to facilitate a meaningful sorting of the alignment for illustration purposes. The alignment was obtained using MUSCLE (Multiple Sequence Comparison by Log-Expectation [112, 113]), and is given in clustal format.

**S6 Dataset. Raw T1 complementation counts.** Numerical data and calculations supporting the histograms in Figs 2C-D, 4B, 6B-C, and S7.

**S7 Dataset. Full length alignment of YTHDF proteins for the calculation of evolutionary divergence of *Ath* ECTs.** Full-length amino acid sequence alignment of 101 YTHDF proteins from selected taxa (bryophytes, basal angiosperms, magnoliids and dicots) used for pairwise comparisons to calculate evolutionary distances between *Mpo* DFE and *Ath* ECTs (S8 Dataset, Fig 6A). All the sequences with their identifiers and the taxonomic classification of the organisms can be found in S1 Dataset. The alignment was computed with MUSCLE (Multiple Sequence Comparison by Log-Expectation [112, 113]), and is given in clustal format.

**S8 Dataset. Evolutionary divergence of *Ath* ECTs.** Matrix with estimates of evolutionary divergence (number of amino acid substitutions per site) between 101 land plant proteins from selected taxa (bryophytes, basal angiosperms, magnoliids and dicots) according to the alignment in S7 Dataset. The analysis was conducted using the Equal Input model [121] in MEGA11 [122, 135]. All positions containing gaps and missing data were eliminated (complete deletion option). There were a total of 171 positions in the final dataset. A complete matrix (‘Matrix output’) as well as a selection of the comparisons regarding *Ath* ECTs vs. *Mpo* DFE, is extracted from the matrix in a separate sheet (‘Selection’). Color-coding follows the rest of the figures in the article. The graphic representation of the values is shown in Fig 6A.

**S9 Dataset. Raw data and calculations supporting qPCR data in Fig 3H**

**S1 Movie. ECT2 simulation.** Coarse-grained molecular dynamics simulation of 100 ECT2 IDRs using the CALVADOS 2 forcefield (7.5 microseconds simulation time) [126]. The proteins are colored differently to distinguish individual chains.

**S2 Movie. ECT1 simulation.** Coarse-grained molecular dynamics simulation of 100 ECT1 IDRs using the CALVADOS 2 forcefield (7.5 microseconds simulation time) [126]. The proteins are colored differently to distinguish individual chains.

**S3 Movie. ECT11 simulation.** Coarse-grained molecular dynamics simulation of 100 ECT11 IDRs using the CALVADOS 2 forcefield (7.5 microseconds simulation time) [126]. The proteins are colored differently to distinguish individual chains.

## References

1. Zhong S, Li H, Bodi Z, Button J, Vespa L, Herzog M, et al. MTA is an Arabidopsis messenger RNA adenosine methylase and interacts with a homolog of a sex-specific splicing factor. Plant Cell. 2008;20(5):1278–88. doi: 10.1105/tpc.108.058883. PubMed PMID: 18505803; PubMed Central PMCID: PMC2438467.

2. Geula S, Moshitch-Moshkovitz S, Dominissini D, Mansour AA, Kol N, Salmon-Divon M, et al. m6A mRNA methylation facilitates resolution of naïve pluripotency toward differentiation. Science. 2015;347(6225):1002-6. doi: 10.1126/science.1261417.

3. Liu N, Dai Q, Zheng G, He C, Parisien M, Pan T. N6-methyladenosine-dependent RNA structural switches regulate RNA-protein interactions. Nature. 2015;518(7540):560-4. doi: 10.1038/nature14234. PubMed PMID: 25719671; PubMed Central PMCID: PMC4355918.

4. Roost C, Lynch SR, Batista PJ, Qu K, Chang HY, Kool ET. Structure and Thermodynamics of N6-Methyladenosine in RNA: A Spring-Loaded Base Modification. J Am Chem Soc. 2015;137(5):2107–15. doi: 10.1021/ja513080v.

5. Spitale RC, Flynn RA, Zhang QC, Crisalli P, Lee B, Jung J-W, et al. Structural imprints in vivo decode RNA regulatory mechanisms. Nature. 2015;519(7544):486–90. Epub 2015/03/18. doi: 10.1038/nature14263. PubMed PMID: 25799993.

6. Patil DP, Pickering BF, Jaffrey SR. Reading m6A in the Transcriptome: m6A-Binding Proteins. Trends Cell Biol. 2018;28(2):113–27. doi: 10.1016/j.tcb.2017.10.001.

7. Dominissini D, Moshitch-Moshkovitz S, Schwartz S, Salmon-Divon M, Ungar L, Osenberg S, et al. Topology of the human and mouse m6A RNA methylomes revealed by m6A-seq. Nature. 2012;485(7397):201-6. doi: 10.1038/nature11112. PubMed PMID: 22575960.

8. Stoilov P, Rafalska I, Stamm S. YTH: a new domain in nuclear proteins. Trends Biochem Sci. 2002;27(10):495–7. doi: 10.1016/S0968-0004(02)02189-8.

9. Wang X, Lu Z, Gomez A, Hon GC, Yue Y, Han D, et al. N6-methyladenosine-dependent regulation of messenger RNA stability. Nature. 2014;505(7481):117-20. doi: 10.1038/nature12730. PubMed PMID: 24284625.

10. Li F, Zhao D, Wu J, Shi Y. Structure of the YTH domain of human YTHDF2 in complex with an m6A mononucleotide reveals an aromatic cage for m6A recognition. Cell Res. 2014;24(12):1490–2. doi: 10.1038/cr.2014.153.

11. Luo S, Tong L. Molecular basis for the recognition of methylated adenines in RNA by the eukaryotic YTH domain. Proc Natl Acad Sci USA. 2014;111(38):13834–9. doi: 10.1073/pnas.1412742111.

12. Theler D, Dominguez C, Blatter M, Boudet J, Allain FH. Solution structure of the YTH domain in complex with N6-methyladenosine RNA: a reader of methylated RNA. Nucleic Acids Res. 2014;42(22):13911–9. doi: 10.1093/nar/gku1116. PubMed PMID: 25389274; PubMed Central PMCID: PMC4267619.

13. Xu C, Wang X, Liu K, Roundtree IA, Tempel W, Li Y, et al. Structural basis for selective binding of m6A RNA by the YTHDC1 YTH domain. Nat Chem Biol. 2014;10(11):927–9. doi: 10.1038/nchembio.1654.

14. Zhu T, Roundtree IA, Wang P, Wang X, Wang L, Sun C, et al. Crystal structure of the YTH domain of YTHDF2 reveals mechanism for recognition of N6-methyladenosine. Cell Res. 2014;24(12):1493–6. doi: 10.1038/cr.2014.152.

15. Zhang Z, Theler D, Kaminska KH, Hiller M, de la Grange P, Pudimat R, et al. The YTH domain is a novel RNA binding domain. The Journal of biological chemistry. 2010;285(19):14701–10. Epub 02/18. doi: 10.1074/jbc.M110.104711. PubMed PMID: 20167602.

16. Arribas-Hernández L, Bressendorff S, Hansen MH, Poulsen C, Erdmann S, Brodersen P. An m6A-YTH Module Controls Developmental Timing and Morphogenesis in Arabidopsis. Plant Cell. 2018;30(5):952–67.

17. Arribas-Hernández L, Simonini S, Hansen MH, Paredes EB, Bressendorff S, Dong Y, et al. Recurrent requirement for the m6A-ECT2/ECT3/ECT4 axis in the control of cell proliferation during plant organogenesis. Development. 2020;147(14):dev189134. doi: 10.1242/dev.189134.

18. Lasman L, Krupalnik V, Viukov S, Mor N, Aguilera-Castrejon A, Schneir D, et al. Context-dependent functional compensation between Ythdf m6A reader proteins. Genes Dev. 2020;34(19-20):1373–91.

19. Ivanova I, Much C, Di Giacomo M, Azzi C, Morgan M, Moreira PN, et al. The RNA m6A reader YTHDF2 is essential for the post-transcriptional regulation of the maternal transcriptome and oocyte competence. Mol Cell. 2017;67(6):1059–67. doi: 10.1016/j.molcel.2017.08.003.

20. Li M, Zhao X, Wang W, Shi H, Pan Q, Lu Z, et al. Ythdf2-mediated m6A mRNA clearance modulates neural development in mice. Genome biology. 2018;19(1):69. doi: 10.1186/s13059-018-1436-y.

21. Shi H, Zhang X, Weng Y-L, Lu Z, Liu Y, Lu Z, et al. m6A facilitates hippocampus-dependent learning and memory through YTHDF1. Nature. 2018;563(7730):249-53. doi: 10.1038/s41586-018-0666-1.

22. Kontur C, Jeong M, Cifuentes D, Giraldez AJ. Ythdf m6A Readers Function Redundantly during Zebrafish Development. Cell Rep. 2020;33(13):108598. doi: 10.1016/j.celrep.2020.108598.

23. Arribas-Hernández L, Rennie S, Schon M, Porcelli C, Enugutti B, Andersson R, et al. The YTHDF proteins ECT2 and ECT3 bind largely overlapping target sets and influence target mRNA abundance, not alternative polyadenylation. eLife. 2021;10:e72377. doi: 10.7554/eLife.72377.

24. Lence T, Akhtar J, Bayer M, Schmid K, Spindler L, Ho CH, et al. m6A modulates neuronal functions and sex determination in Drosophila. Nature. 2016;540:242. doi: 10.1038/nature20568.

25. Haussmann IU, Bodi Z, Sanchez-Moran E, Mongan NP, Archer N, Fray RG, et al. m6A potentiates Sxl alternative pre-mRNA splicing for robust Drosophila sex determination. Nature. 2016;540:301. doi: 10.1038/nature20577.

26. Scutenaire J, Deragon J-M, Jean V, Benhamed M, Raynaud C, Favory J-J, et al. The YTH Domain Protein ECT2 Is an m6A Reader Required for Normal Trichome Branching in Arabidopsis. Plant Cell. 2018;30(5):986–1005. doi: 10.1105/tpc.17.00854.

27. Li D, Zhang H, Hong Y, Huang L, Li X, Zhang Y, et al. Genome-wide identification, biochemical characterization, and expression analyses of the YTH domain-containing RNA-binding protein family in Arabidopsis and rice. Plant Mol Biol Report. 2014;32(6):1169–86. doi: 10.1007/s11105-014-0724-2.

28. Du H, Zhao Y, He J, Zhang Y, Xi H, Liu M, et al. YTHDF2 destabilizes m6A-containing RNA through direct recruitment of the CCR4–NOT deadenylase complex. Nat Commun. 2016;7:12626. doi: 10.1038/ncomms12626.

29. Park OH, Ha H, Lee Y, Boo SH, Kwon DH, Song HK, et al. Endoribonucleolytic Cleavage of m6A-Containing RNAs by RNase P/MRP Complex. Mol Cell. 2019;74(3):494–507.e8. doi: 10.1016/j.molcel.2019.02.034.

30. Boo SH, Ha H, Lee Y, Shin M-K, Lee S, Kim YK. UPF1 promotes rapid degradation of m6A-containing RNAs. Cell Rep. 2022;39(8):110861. doi: 10.1016/j.celrep.2022.110861.

31. Arribas-Hernández L, Rennie S, Köster T, Porcelli C, Lewinski M, Staiger D, et al. Principles of mRNA targeting via the Arabidopsis m6A-binding protein ECT2. eLife. 2021;10:e72375. doi: 10.7554/eLife.72375.

32. Brodsky S, Jana T, Mittelman K, Chapal M, Kumar DK, Carmi M, et al. Intrinsically Disordered Regions Direct Transcription Factor In Vivo Binding Specificity. Mol Cell. 2020;79(3):459–71.e4. doi: 10.1016/j.molcel.2020.05.032.

33. Wang J, Wang L, Diao J, Shi YG, Shi Y, Ma H, et al. Binding to m6A RNA promotes YTHDF2-mediated phase separation. Protein & Cell. 2020;11(4):304–7. doi: 10.1007/s13238-019-00660-2.

34. Gao Y, Pei G, Li D, Li R, Shao Y, Zhang QC, et al. Multivalent m6A motifs promote phase separation of YTHDF proteins. Cell Res. 2019;29(9):767–9. doi: 10.1038/s41422-019-0210-3.

35. Ries RJ, Zaccara S, Klein P, Olarerin-George A, Namkoong S, Pickering BF, et al. m6A enhances the phase separation potential of mRNA. Nature. 2019;571(7765):424-8. doi: 10.1038/s41586-019-1374-1.

36. Fu Y, Zhuang X. m6A-binding YTHDF proteins promote stress granule formation. Nat Chem Biol. 2020;16(9):955–63. doi: 10.1038/s41589-020-0524-y.

37. Scutenaire J, Plassard D, Matelot M, Villa T, Zumsteg J, Libri D, et al. The S. cerevisiae m6A-reader Pho92 promotes timely meiotic recombination by controlling key methylated transcripts. Nucleic Acids Res. 2022:gkac640. doi: 10.1093/nar/gkac640.

38. Varier RA, Sideri T, Capitanchik C, Manova Z, Calvani E, Rossi A, et al. N6-methyladenosine (m6A) reader Pho92 is recruited co-transcriptionally and couples translation to mRNA decay to promote meiotic fitness in yeast. eLife. 2022;11:e84034. doi: 10.7554/eLife.84034.

39. Zaccara S, Jaffrey SR. A Unified Model for the Function of YTHDF Proteins in Regulating m(6)A-Modified mRNA. Cell. 2020;181(7):1582–95.e18. Epub 2020/06/04. doi: 10.1016/j.cell.2020.05.012. PubMed PMID: 32492408.

40. Ok SH, Jeong HJ, Bae JM, Shin JS, Luan S, Kim KN. Novel CIPK1-associated proteins in Arabidopsis contain an evolutionarily conserved C-terminal region that mediates nuclear localization. Plant Physiol. 2005;139(1):138–50. doi: 10.1104/pp.105.065649. PubMed PMID: 16113215.

41. Bergeron-Sandoval L-P, Safaee N, Michnick Stephen W. Mechanisms and Consequences of Macromolecular Phase Separation. Cell. 2016;165(5):1067–79. doi: 10.1016/j.cell.2016.05.026.

42. Borcherds W, Bremer A, Borgia MB, Mittag T. How do intrinsically disordered protein regions encode a driving force for liquid–liquid phase separation? Curr Opin Struct Biol. 2021;67:41–50. doi: 10.1016/j.sbi.2020.09.004.

43. Murakami S, Jaffrey SR. Hidden codes in mRNA: Control of gene expression by m6A. Mol Cell. 2022;82(12):2236–51. doi: 10.1016/j.molcel.2022.05.029.

44. Wang X, Zhao Boxuan S, Roundtree Ian A, Lu Z, Han D, Ma H, et al. N6-methyladenosine modulates messenger RNA translation efficiency. Cell. 2015;161(6):1388–99. doi: 10.1016/j.cell.2015.05.014.

45. Shi H, Wang X, Lu Z, Zhao BS, Ma H, Hsu PJ, et al. YTHDF3 facilitates translation and decay of N6-methyladenosine-modified RNA. Cell Res. 2017;27:315–28. doi: 10.1038/cr.2017.15.

46. Li A, Chen Y-S, Ping X-L, Yang X, Xiao W, Yang Y, et al. Cytoplasmic m6A reader YTHDF3 promotes mRNA translation. Cell Res. 2017;27:444. doi: 10.1038/cr.2017.10.

47. Li Y, Bedi RK, Moroz-Omori EV, Caflisch A. Structural and Dynamic Insights into Redundant Function of YTHDF Proteins. Journal of Chemical Information and Modeling. 2020;60(12):5932–5. doi: 10.1021/acs.jcim.0c01029.

48. Zou Z, Sepich-Poore C, Zhou X, Wei J, He C. The mechanism underlying redundant functions of the YTHDF proteins. Genome biology. 2023;24(1):17. doi: 10.1186/s13059-023-02862-8.

49. Fray RG, Simpson GG. The Arabidopsis epitranscriptome. Curr Opin Plant Biol. 2015;27:17–21. doi: 10.1016/j.pbi.2015.05.015.

50. Bodi Z, Zhong S, Mehra S, Song J, Graham N, Li H, et al. Adenosine methylation in Arabidopsis mRNA is associated with the 3’ end and reduced levels cause developmental defects. Front Plant Sci. 2012;3:48. doi: 10.3389/fpls.2012.00048. PubMed PMID: 22639649.

51. Shen L, Liang Z, Gu X, Chen Y, Teo Zhi Wei N, Hou X, et al. N6-methyladenosine RNA modification regulates shoot stem cell fate in Arabidopsis. Dev Cell. 2016;38(2):186–200. doi: 10.1016/j.devcel.2016.06.008.

52. Růžička K, Zhang M, Campilho A, Bodi Z, Kashif M, Saleh M, et al. Identification of factors required for m6A mRNA methylation in Arabidopsis reveals a role for the conserved E3 ubiquitin ligase HAKAI. New Phytol. 2017;215(1):157–72. doi: 10.1111/nph.14586.

53. Pires ND, Dolan L. Morphological evolution in land plants: new designs with old genes. Philosophical Transactions of the Royal Society B: Biological Sciences. 2012;367(1588):508-18. doi: 10.1098/rstb.2011.0252.

54. 54. One Thousand Plant Transcriptomes I. One thousand plant transcriptomes and the phylogenomics of green plants. Nature. 2019;574(7780):679–85. Epub 2019/10/23. doi: 10.1038/s41586-019-1693-2. PubMed PMID: 31645766.

55. de Vries J, Archibald JM. Plant evolution: landmarks on the path to terrestrial life. New Phytol. 2018;217(4):1428–34. doi: 10.1111/nph.14975.

56. Harris BJ, Harrison CJ, Hetherington AM, Williams TA. Phylogenomic Evidence for the Monophyly of Bryophytes and the Reductive Evolution of Stomata. Current Biology. 2020;30(11):2001–12.e2. doi: 10.1016/j.cub.2020.03.048.

57. Li F-W, Nishiyama T, Waller M, Frangedakis E, Keller J, Li Z, et al. Anthoceros genomes illuminate the origin of land plants and the unique biology of hornworts. Nature plants. 2020;6(3):259–72. Epub 2020/03/13. doi: 10.1038/s41477-020-0618-2. PubMed PMID: 32170292.

58. Puttick MN, Morris JL, Williams TA, Cox CJ, Edwards D, Kenrick P, et al. The Interrelationships of Land Plants and the Nature of the Ancestral Embryophyte. Current Biology. 2018;28(5):733–45.e2. doi: 10.1016/j.cub.2018.01.063.

59. Su D, Yang L, Shi X, Ma X, Zhou X, Hedges SB, et al. Large-Scale Phylogenomic Analyses Reveal the Monophyly of Bryophytes and Neoproterozoic Origin of Land Plants. Mol Biol Evol. 2021;38(8):3332–44. doi: 10.1093/molbev/msab106. PubMed PMID: 33871608.

60. Zhang J, Fu X-X, Li R-Q, Zhao X, Liu Y, Li M-H, et al. The hornwort genome and early land plant evolution. Nature plants. 2020;6(2):107–18. Epub 2020/02/10. doi: 10.1038/s41477-019-0588-4. PubMed PMID: 32042158.

61. Xu C, Liu K, Ahmed H, Loppnau P, Schapira M, Min J. Structural Basis for the Discriminative Recognition of N6-Methyladenosine RNA by the Human YT521-B Homology Domain Family of Proteins. Journal of Biological Chemistry. 2015;290(41):24902–13. doi: 10.1074/jbc.M115.680389.

62. Bowman JL. The origin of a land flora. Nature Plants. 2022;8(12):1352–69. doi: 10.1038/s41477-022-01283-y.

63. Magallón S, Hilu KW, Quandt D. Land plant evolutionary timeline: Gene effects are secondary to fossil constraints in relaxed clock estimation of age and substitution rates. Am J Bot. 2013;100(3):556–73. doi: 10.3732/ajb.1200416.

64. Lan T, Xiong W, Chen X, Mo B, Tang G. Plant cytoplasmic ribosomal proteins: an update on classification, nomenclature, evolution and resources. Plant J. 2022;110(1):292–318. doi: 10.1111/tpj.15667.

65. Scarpin MR, Busche M, Martinez RE, Harper LC, Reiser L, Szakonyi D, et al. An updated nomenclature for plant ribosomal protein genes. Plant Cell. 2023;35(2):640–3. doi: 10.1093/plcell/koac333.

66. Weijers D, Franke-van Dijk M, Vencken R-J, Quint A, Hooykaas P, Offringa R. An Arabidopsis Minute-like phenotype caused by a semi-dominant mutation in a RIBOSOMAL PROTEIN S5 gene. Development. 2001;128(21):4289–99. doi: 10.1242/dev.128.21.4289.

67. Fagard M, Vaucheret H. (TRANS)GENE SILENCING IN PLANTS: How Many Mechanisms? Annu Rev Plant Physiol Plant Mol Biol. 2000;51(1):167–94. doi: 10.1146/annurev.arplant.51.1.167.

68. Balacco DL, Soller M. The m6A Writer: Rise of a Machine for Growing Tasks. Biochemistry. 2019;58(5):363–78. doi: 10.1021/acs.biochem.8b01166.

69. Wang C, Zhu Y, Bao H, Jiang Y, Xu C, Wu J, et al. A novel RNA-binding mode of the YTH domain reveals the mechanism for recognition of determinant of selective removal by Mmi1. Nucleic Acids Res. 2016;44(2):969–82. doi: 10.1093/nar/gkv1382.

70. Dalmay T, Hamilton A, Rudd S, Angell S, Baulcombe DC. An RNA-dependent RNA polymerase gene in Arabidopsis is required for posttranscriptional gene silencing mediated by a transgene but not by a virus. Cell. 2000;101(5):543–53. PubMed PMID: 10850496.

71. Mourrain P, Beclin C, Elmayan T, Feuerbach F, Godon C, Morel J-B, et al. *Arabidopsis SGS2* and *SGS3* genes are required for posttranscriptional gene silencing and natural virus resistance. Cell. 2000;101:533–42.

72. Peragine A, Yoshikawa M, Wu G, Albrecht HL, Poethig RS. SGS3 and SGS2/SDE1/RDR6 are required for juvenile development and the production of trans-acting siRNAs in Arabidopsis. Genes Dev. 2004;18(19):2368–79. PubMed PMID: 15466488.

73. Zhang H, Zhang F, Yu Y, Feng L, Jia J, Liu B, et al. A Comprehensive Online Database for Exploring ∼20,000 Public Arabidopsis RNA-Seq Libraries. Molecular Plant. 2020;13(9):1231–3. doi: 10.1016/j.molp.2020.08.001.

74. Jumper J, Evans R, Pritzel A, Green T, Figurnov M, Ronneberger O, et al. Highly accurate protein structure prediction with AlphaFold. Nature. 2021;596(7873):583-9. doi: 10.1038/s41586-021- 03819-2.

75. Varadi M, Anyango S, Deshpande M, Nair S, Natassia C, Yordanova G, et al. AlphaFold Protein Structure Database: massively expanding the structural coverage of protein-sequence space with high-accuracy models. Nucleic Acids Res. 2022;50(D1):D439–D44. doi: 10.1093/nar/gkab1061.

76. Biasini M, Bienert S, Waterhouse A, Arnold K, Studer G, Schmidt T, et al. SWISS-MODEL: modelling protein tertiary and quaternary structure using evolutionary information. Nucleic Acids Res. 2014;42(W1):W252–W8. doi: 10.1093/nar/gku340.

77. Piovesan D, Del Conte A, Clementel D, Monzon Alexander M, Bevilacqua M, Aspromonte Maria C, et al. MobiDB: 10 years of intrinsically disordered proteins. Nucleic Acids Res. 2023;51(D1):D438–D44. doi: 10.1093/nar/gkac1065.

78. Tesei G, Schulze TK, Crehuet R, Lindorff-Larsen K. Accurate model of liquid-liquid phase behavior of intrinsically disordered proteins from optimization of single-chain properties. Proc Natl Acad Sci USA. 2021;118(44):e2111696118. doi: 10.1073/pnas.2111696118. PubMed PMID: 34716273.

79. Kato M, McKnight SL. The low-complexity domain of the FUS RNA binding protein self-assembles via the mutually exclusive use of two distinct cross-β cores. Proc Natl Acad Sci USA. 2021;118(42):e2114412118. doi: 10.1073/pnas.2114412118.

80. Kato M, Han TW, Xie S, Shi K, Du X, Wu LC, et al. Cell-free formation of RNA granules: low complexity sequence domains form dynamic fibers within hydrogels. Cell. 2012;149(4):753–67. doi: 10.1016/j.cell.2012.04.017. PubMed PMID: 22579281.

81. Patel A, Lee Hyun O, Jawerth L, Maharana S, Jahnel M, Hein Marco Y, et al. A Liquid-to-Solid Phase Transition of the ALS Protein FUS Accelerated by Disease Mutation. Cell. 2015;162(5):1066–77. doi: 10.1016/j.cell.2015.07.047.

82. Schwartz Jacob C, Wang X, Podell Elaine R, Cech Thomas R. RNA Seeds Higher-Order Assembly of FUS Protein. Cell Rep. 2013;5(4):918–25. doi: 10.1016/j.celrep.2013.11.017.

83. Bianchi G, Mangiagalli M, Barbiroli A, Longhi S, Grandori R, Santambrogio C, et al. Distribution of Charged Residues Affects the Average Size and Shape of Intrinsically Disordered Proteins. Biomolecules. 2022;12(4). doi: 10.3390/biom12040561.

84. Das RK, Huang Y, Phillips AH, Kriwacki RW, Pappu RV. Cryptic sequence features within the disordered protein p27Kip1 regulate cell cycle signaling. Proc Natl Acad Sci USA. 2016;113(20):5616–21. doi: 10.1073/pnas.1516277113.

85. Bowman JL, Kohchi T, Yamato KT, Jenkins J, Shu S, Ishizaki K, et al. Insights into Land Plant Evolution Garnered from the Marchantia polymorpha Genome. Cell. 2017;171(2):287–304.e15. doi: 10.1016/j.cell.2017.09.030.

86. Wang L, Roossinck MJ. Comparative analysis of expressed sequences reveals a conserved pattern of optimal codon usage in plants. Plant Mol Biol. 2006;61(4):699–710. doi: 10.1007/s11103-006-0041-8.

87. Soltis PS, Soltis DE, Chase MW. Angiosperm phylogeny inferred from multiple genes as a tool for comparative biology. Nature. 1999;402(6760):402-4. doi: 10.1038/46528.

88. Qiu Y-L, Lee J, Bernasconi-Quadroni F, Soltis DE, Soltis PS, Zanis M, et al. The earliest angiosperms: evidence from mitochondrial, plastid and nuclear genomes. Nature. 1999;402(6760):404-7. doi: 10.1038/46536.

89. Sommer S, Lavi U, Darnell JE. The absolute frequency of labeled N-6-methyladenosine in HeLa cell messenger RNA decreases with label time. Journal of Molecular Biology. 1978;124(3):487–99. doi: 10.1016/0022-2836(78)90183-3.

90. Wang Y, Li Y, Toth JI, Petroski MD, Zhang Z, Zhao JC. N6-methyladenosine modification destabilizes developmental regulators in embryonic stem cells. Nature cell biology. 2014;16(2):191–8. doi: 10.1038/ncb2902. PubMed PMID: 24394384.

91. Ke S, Pandya-Jones A, Saito Y, Fak JJ, Vågbø CB, Geula S, et al. m6A mRNA modifications are deposited in nascent pre-mRNA and are not required for splicing but do specify cytoplasmic turnover. Genes Dev. 2017;31(10):990–1006.

92. Herzog VA, Reichholf B, Neumann T, Rescheneder P, Bhat P, Burkard TR, et al. Thiol-linked alkylation of RNA to assess expression dynamics. Nat Meth. 2017;14:1198. doi: 10.1038/nmeth.4435.

93. Anderson SJ, Kramer MC, Gosai SJ, Yu X, Vandivier LE, Nelson ADL, et al. N(6)-Methyladenosine Inhibits Local Ribonucleolytic Cleavage to Stabilize mRNAs in Arabidopsis. Cell Rep. 2018;25(5):1146–57. doi: 10.1016/j.celrep.2018.10.020. PubMed PMID: 30380407.

94. Wei L-H, Song P, Wang Y, Lu Z, Tang Q, Yu Q, et al. The m6A Reader ECT2 Controls Trichome Morphology by Affecting mRNA Stability in Arabidopsis. Plant Cell. 2018;30(5):968–85. doi: 10.1105/tpc.17.00934.

95. Parker MT, Knop K, Sherwood AV, Schurch NJ, Mackinnon K, Gould PD, et al. Nanopore direct RNA sequencing maps the complexity of Arabidopsis mRNA processing and m6A modification. eLife. 2020;9:e49658. doi: 10.7554/eLife.49658.

96. Song P, Wei L, Chen Z, Cai Z, Lu Q, Wang C, et al. m6A readers ECT2/ECT3/ECT4 enhance mRNA stability through direct recruitment of the poly(A) binding proteins in Arabidopsis. Genome biology. 2023;24(1):103. doi: 10.1186/s13059-023-02947-4.

97. Arribas-Hernández L, Brodersen P. Occurrence and functions of m6A and other covalent modifications in plant mRNA. Plant Physiol. 2020;182(1):79–96. doi: 10.1104/pp.19.01156.

98. Tankmar MD, Reichel M, Arribas-Hernández L, Brodersen P. A YTHDF-PABP axis is required for m6A-mediated organogenesis in plants. bioRxiv. 2023:2023.07.03.547513. doi: 10.1101/2023.07.03.547513.

99. Al Jewari C, Baldauf SL. An excavate root for the eukaryote tree of life. Science Advances. 9(17):eade4973. doi: 10.1126/sciadv.ade4973.

100. Burki F, Roger AJ, Brown MW, Simpson AGB. The New Tree of Eukaryotes. Trends in Ecology & Evolution. 2020;35(1):43–55. doi: 10.1016/j.tree.2019.08.008.

101. McCourt RM, Lewis LA, Strother PK, Delwiche CF, Wickett NJ, de Vries J, et al. Green land: Multiple perspectives on green algal evolution and the earliest land plants. Am J Bot. 2023;110(5):e16175. doi: 10.1002/ajb2.16175.

102. Graham LE, Cook ME, Busse JS. The origin of plants: Body plan changes contributing to a major evolutionary radiation. Proc Natl Acad Sci USA. 2000;97(9):4535–40. doi: 10.1073/pnas.97.9.4535.

103. Price DC, Goodenough UW, Roth R, Lee J-H, Kariyawasam T, Mutwil M, et al. Analysis of an improved Cyanophora paradoxa genome assembly. DNA Research. 2019;26(4):287–99. doi: 10.1093/dnares/dsz009.

104. Van Etten J, Benites LF, Stephens TG, Yoon HS, Bhattacharya D. Algae obscura: The potential of rare species as model systems. J Phycol. 2023;59(2):293–300. doi: 10.1111/jpy.13321.

105. Gan S. Mitotic and Postmitotic Senescence in Plants. Science of Aging Knowledge Environment. 2003;2003(38):re7–re. doi: 10.1126/sageke.2003.38.re7.

106. Wuest SE, Philipp MA, Guthörl D, Schmid B, Grossniklaus U. Seed Production Affects Maternal Growth and Senescence in Arabidopsis Plant Physiol. 2016;171(1):392–404. doi: 10.1104/pp.15.01995.

107. Goodstein DM, Shu S, Howson R, Neupane R, Hayes RD, Fazo J, et al. Phytozome: a comparative platform for green plant genomics. Nucleic Acids Res. 2012;40(D1):D1178–D86. doi: 10.1093/nar/gkr944.

108. Gu K-J, Lin C-F, Wu J-J, Zhao Y-P. GinkgoDB: an ecological genome database for the living fossil, Ginkgo biloba. Database. 2022;2022:baac046. doi: 10.1093/database/baac046.

109. Li F-W, Brouwer P, Carretero-Paulet L, Cheng S, de Vries J, Delaux P-M, et al. Fern genomes elucidate land plant evolution and cyanobacterial symbioses. Nature Plants. 2018;4(7):460–72. doi: 10.1038/s41477-018-0188-8.

110. Grigoriev IV, Nordberg H, Shabalov I, Aerts A, Cantor M, Goodstein D, et al. The Genome Portal of the Department of Energy Joint Genome Institute. Nucleic Acids Res. 2012;40(D1):D26–D32. doi: 10.1093/nar/gkr947.

111. Nordberg H, Cantor M, Dusheyko S, Hua S, Poliakov A, Shabalov I, et al. The genome portal of the Department of Energy Joint Genome Institute: 2014 updates. Nucleic Acids Res. 2014;42(D1):D26–D31. doi: 10.1093/nar/gkt1069.

112. Edgar RC. MUSCLE: multiple sequence alignment with high accuracy and high throughput. Nucleic Acids Res. 2004;32(5):1792–7. doi: 10.1093/nar/gkh340. PubMed PMID: 15034147.

113. Edgar RC. MUSCLE: a multiple sequence alignment method with reduced time and space complexity. BMC Bioinformatics. 2004;5(1):113. doi: 10.1186/1471-2105-5-113.

114. Waterhouse AM, Procter JB, Martin DMA, Clamp M, Barton GJ. Jalview Version 2—a multiple sequence alignment editor and analysis workbench. Bioinformatics. 2009;25(9):1189–91. doi: 10.1093/bioinformatics/btp033.

115. Nguyen L-T, Schmidt HA, von Haeseler A, Minh BQ. IQ-TREE: A Fast and Effective Stochastic Algorithm for Estimating Maximum-Likelihood Phylogenies. Mol Biol Evol. 2015;32(1):268–74. doi: 10.1093/molbev/msu300.

116. Minh BQ, Schmidt HA, Chernomor O, Schrempf D, Woodhams MD, von Haeseler A, et al. IQ-TREE 2: New Models and Efficient Methods for Phylogenetic Inference in the Genomic Era. Mol Biol Evol. 2020;37(5):1530–4. doi: 10.1093/molbev/msaa015.

117. Minh BQ, Dang CC, Vinh LS, Lanfear R. QMaker: Fast and Accurate Method to Estimate Empirical Models of Protein Evolution. Syst Biol. 2021;70(5):1046–60. doi: 10.1093/sysbio/syab010.

118. Ran J-H, Shen T-T, Wang M-M, Wang X-Q. Phylogenomics resolves the deep phylogeny of seed plants and indicates partial convergent or homoplastic evolution between Gnetales and angiosperms. Proceedings of the Royal Society B: Biological Sciences. 2018;285(1881):20181012. doi: 10.1098/rspb.2018.1012.

119. Kalyaanamoorthy S, Minh BQ, Wong TKF, von Haeseler A, Jermiin LS. ModelFinder: fast model selection for accurate phylogenetic estimates. Nat Meth. 2017;14(6):587–9. Epub 2017/05/08. doi: 10.1038/nmeth.4285. PubMed PMID: 28481363.

120. Shimodaira H, Hasegawa M. Multiple Comparisons of Log-Likelihoods with Applications to Phylogenetic Inference. Mol Biol Evol. 1999;16(8):1114-. doi: 10.1093/oxfordjournals.molbev.a026201.

121. Tajima F, Nei M. Estimation of evolutionary distance between nucleotide sequences. Mol Biol Evol. 1984;1(3):269–85. doi: 10.1093/oxfordjournals.molbev.a040317.

122. Tamura K, Stecher G, Kumar S. MEGA11: Molecular Evolutionary Genetics Analysis Version 11. Mol Biol Evol. 2021;38(7):3022–7. doi: 10.1093/molbev/msab120. PubMed PMID: 33892491.

123. Bitinaite J, Nichols NM. DNA cloning and engineering by uracil excision. Curr Protoc Mol Biol. 2009;Chapter 3:Unit 3 21. doi: 10.1002/0471142727.mb0321s86. PubMed PMID: 19343708.

124. Nour-Eldin HH, Hansen BG, Norholm MH, Jensen JK, Halkier BA. Advancing uracil-excision based cloning towards an ideal technique for cloning PCR fragments. Nucleic Acids Res. 2006;34(18):e122. doi: 10.1093/nar/gkl635. PubMed PMID: 17000637.

125. Lampropoulos A, Sutikovic Z, Wenzl C, Maegele I, Lohmann JU, Forner J. GreenGate - A Novel, Versatile, and Efficient Cloning System for Plant Transgenesis. PLOS ONE. 2013;8(12):e83043. doi: 10.1371/journal.pone.0083043.

126. Tesei G, Lindorff-Larsen K. Improved predictions of phase behaviour of intrinsically disordered proteins by tuning the interaction range [version 2; peer review: 2 approved]. Open Research Europe. 2023;2(94). doi: 10.12688/openreseurope.14967.2.

127. Dignon GL, Zheng W, Kim YC, Best RB, Mittal J. Sequence determinants of protein phase behavior from a coarse-grained model. PLOS Computational Biology. 2018;14(1):e1005941. doi: 10.1371/journal.pcbi.1005941.

128. Benayad Z, von Bülow S, Stelzl LS, Hummer G. Simulation of FUS Protein Condensates with an Adapted Coarse-Grained Model. Journal of Chemical Theory and Computation. 2021;17(1):525–37. doi: 10.1021/acs.jctc.0c01064.

129. Kleinboelting N, Huep G, Kloetgen A, Viehoever P, Weisshaar B. GABI-Kat SimpleSearch: new features of the Arabidopsis thaliana T-DNA mutant database. Nucleic Acids Res. 2012;40(Database issue):D1211–D5. Epub 2011/11/12. doi: 10.1093/nar/gkr1047. PubMed PMID: 22080561.

130. Robert X, Gouet P. Deciphering key features in protein structures with the new ENDscript server. Nucleic Acids Res. 2014;42(Web Server issue):W320–4. doi: 10.1093/nar/gku316. PubMed PMID: 24753421; PubMed Central PMCID: PMC4086106.

131. Jurrus E, Engel D, Star K, Monson K, Brandi J, Felberg LE, et al. Improvements to the APBS biomolecular solvation software suite. Protein Sci. 2018;27(1):112–28. doi: 10.1002/pro.3280.

132. Akdel M, Pires DEV, Pardo EP, Jänes J, Zalevsky AO, Mészáros B, et al. A structural biology community assessment of AlphaFold2 applications. Nat Struct Mol Biol. 2022;29(11):1056–67. doi: 10.1038/s41594-022-00849-w.

133. Paysan-Lafosse T, Blum M, Chuguransky S, Grego T, Pinto BL, Salazar Gustavo A, et al. InterPro in 2022. Nucleic Acids Res. 2023;51(D1):D418–D27. doi: 10.1093/nar/gkac993.

134. Käll L, Krogh A, Sonnhammer ELL. A Combined Transmembrane Topology and Signal Peptide Prediction Method. Journal of Molecular Biology. 2004;338(5):1027–36. doi: 10.1016/j.jmb.2004.03.016.

135. Stecher G, Tamura K, Kumar S. Molecular Evolutionary Genetics Analysis (MEGA) for macOS. Mol Biol Evol. 2020;37(4):1237–9. doi: 10.1093/molbev/msz312.

