## Supplemental figures for "Insights into the conservation and diversification of the molecular functions of YTHDF proteins"

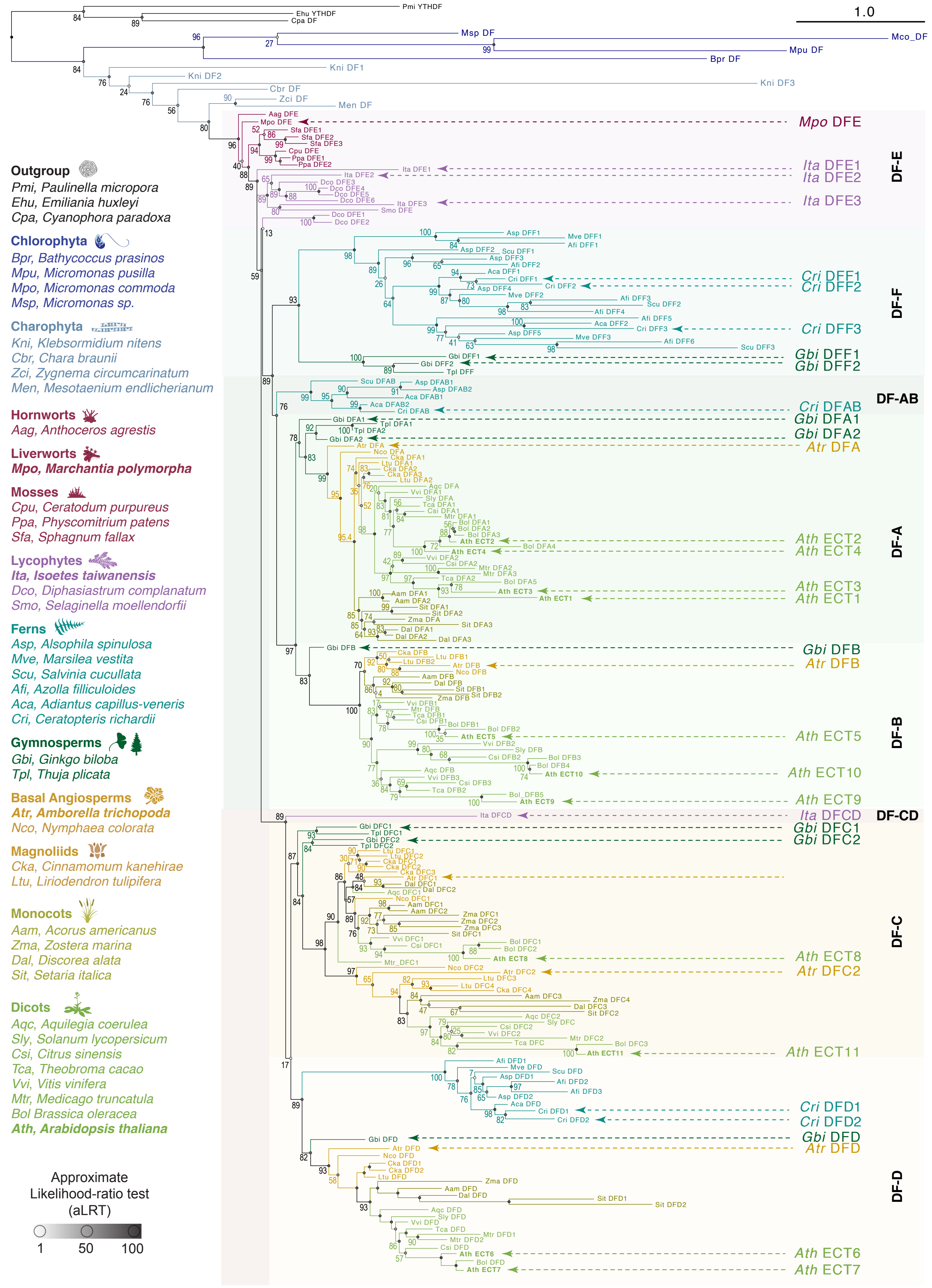


**S1 Fig. Fully annotated phylogenetic tree.** Same phylogenetic analysis of Fig 1, but with rectangular layout, and indicating the protein and species name of all YTHDFs, and aLRT values (%) for all nodes. The representative proteins labeled in Fig 1B are highlighted with larger typography on the right side of this figure as a reference.

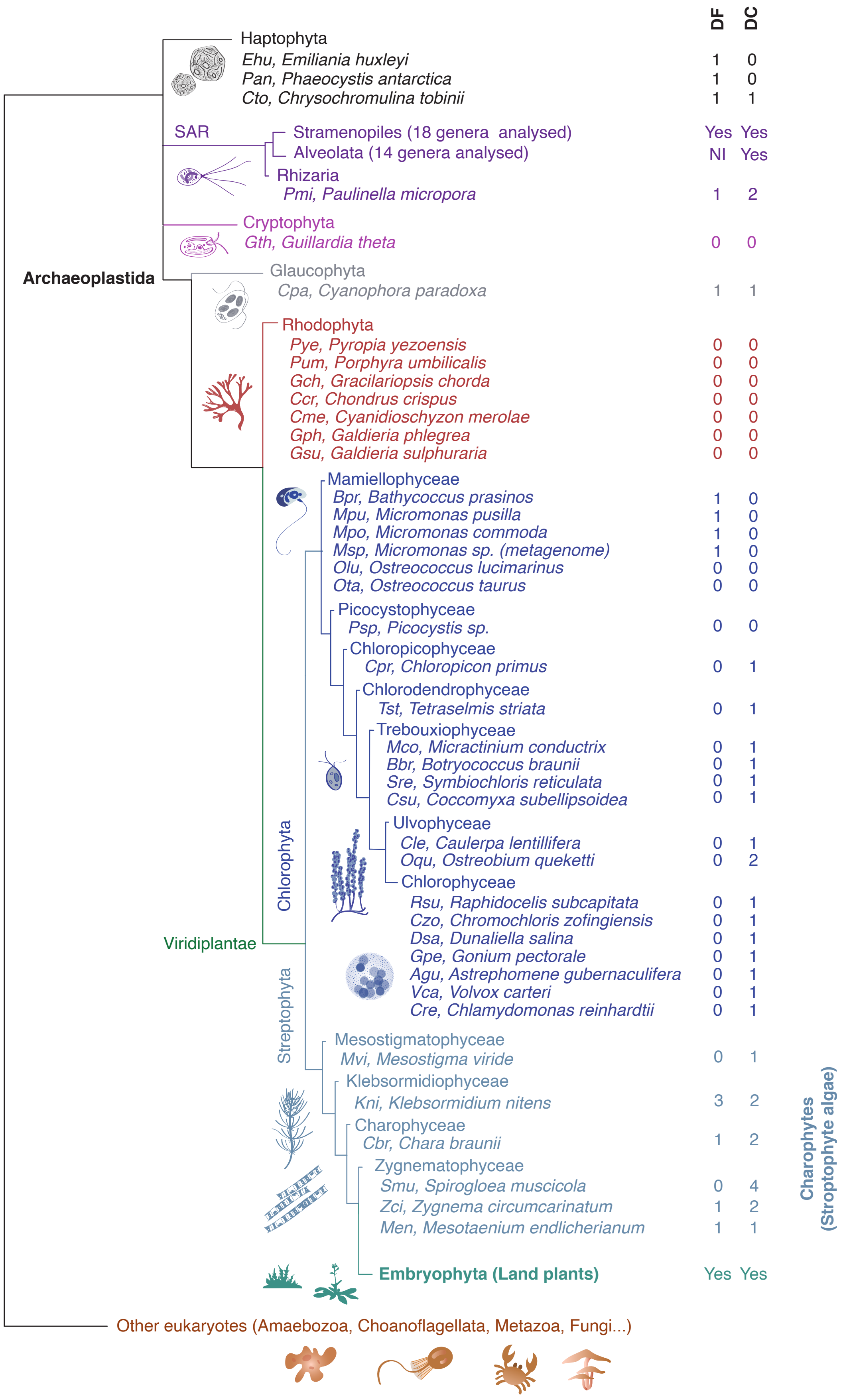


**S2 Fig. YTH domain proteins in Archaeoplastida.** Number of genes encoding YTHDF and YTHDC proteins in several species of Archaeoplastida (see Figs 1 and S1 for detailed data on Embryophyta), and closely related taxa. NI, Not Identified. Phylogenetic relationships between groups according to [55, 97-99] are indicated. Among chlorophytes, YTHDF proteins were found only in species of the *Micromonas* and *Bathycoccus* genera of the basal group *Mamiellophyceae*, although losses in other genera within this group have also occurred (e.g. *Ostreococcus* [67]). Regarding charophytes, the only sequenced species of their earliest-branching group, *Mesostigma viride*, has lost YTHDF proteins altogether, but YTHDF-encoding genes are found in four species of later-diverging groups: *Klebsormidium nitens*, *Chara braunii, Zygnema circumcarinatu*, and *Mesotaenium endlicherianum*.

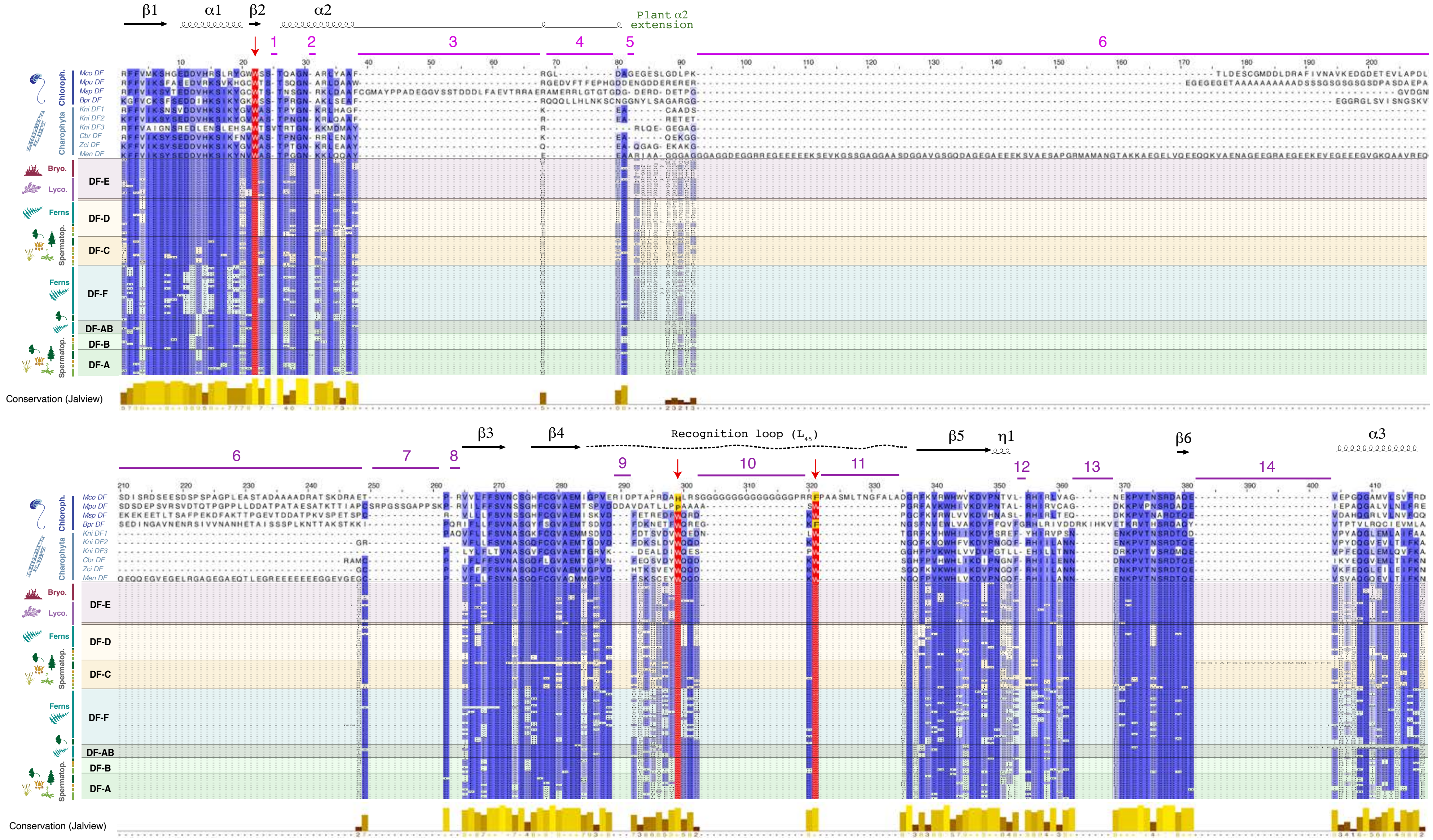


**S3 Fig. Conservation of YTHDF proteins in green algae.** Amino acid sequence alignment of the YTHDF proteins of the Viridiplantae species used for the phylogenetic analysis in Fig 1B, highlighting the family members found in green algae (see species abbreviations in S2 Fig). The frequent amino acid substitutions in structural elements of the green algal orthologs are revealed by the absence of the blue shading that indicates percentage identity (Jalview [112]). In some chlorophyte DFs, these substitutions affect the m⁶A-recognition aromatic cage (red arrows and red/yellow shading), because the two embryophyte-invariant tryptophan residues in the recognition loop (Fig 5A) exhibit changes to proline, histidine or phenylalanine. Chlorophyte DF proteins also contain insertions in their YTH domains, here marked with magenta lines and numbers above the sequences that correspond to collapsed gaps in S4 Fig. The insertions have variable lengths, reaching up to 77 amino acids, and can disrupt the m⁶A-recognition loop, e.g. the poly-Glycine stretch of *Micromonas commoda* DF (insertion 10, GGGGGGGGGGGGGGGPR). In the charophyte DF orthologs, the YTH domains have intact aromatic cages and overall higher conservation than those of chlorophytes, but they also differ considerably from the land plant relatives. For example, the sequences of the β1 and α1 structural elements (Fig 5A) have degenerated in one of the three *Klebsormidium nitens* DF proteins (*Kni* DF3), and a 156-amino acid insertion protruding from the plant-specific α2-extension is apparent in the only DF protein from *Mesotaenium endlicherianum* (*Men* DF), a species that belongs to the sister clade of all embryophytes (S2 Fig).

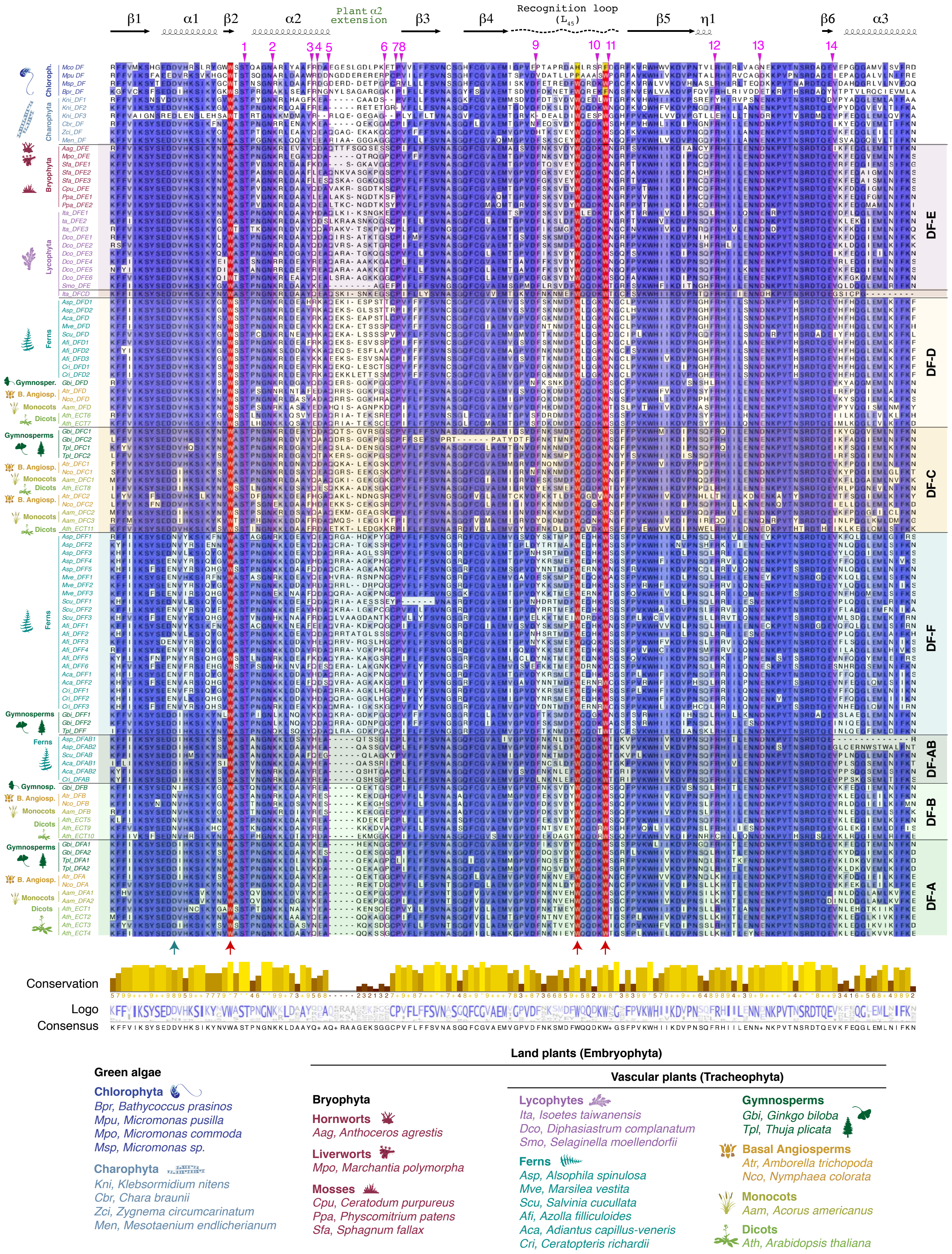


**S4 Fig. Conservation of YTHDF proteins in cryptogams and gymnosperms.** Amino acid sequence alignment of the YTH domains of DF proteins in plant taxa that diverged before the evolution of flowers. A few angiosperm YTHDFs are included in the analysis for comparison. Magenta-colored numbers at the top indicate the positions of amino acid insertions in one or few proteins. Because these insertions create gaps in the aligned sequences of the other homologs, they have been hidden to gain space and clarity, but the corresponding sequences can be found in S3 Fig and S5 Dataset. Red arrows under the alignment point to the conserved aromatic residues that contact m$^6$A, and a green arrow marks the Asp-to-Asn substitution, present in YTHDCs, that increases the affinity for m$^6$A by 15-fold [61]. This substitution is also found in the plant DF-B clade [26], and here revealed in fern DF-F proteins as well. Blue-white coloring and the conservation and consensus tracks at the bottom are computed by Jalview [112] as in S3 Fig.

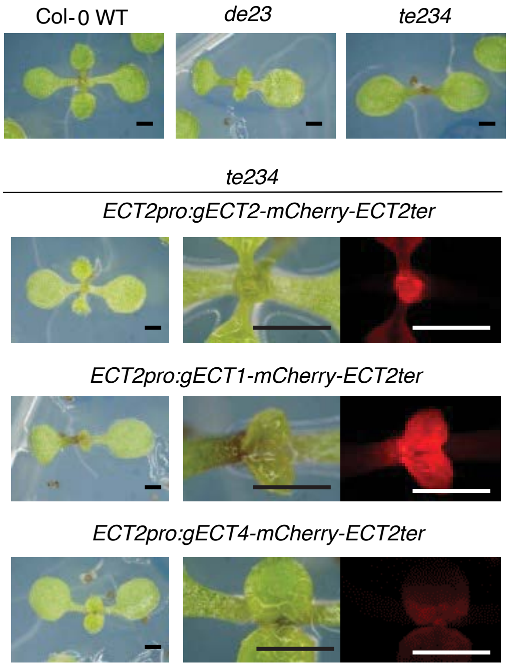


**S5 Fig. Expression of gDNA constructs of *ECT1, ECT2* and *ECT4* under the control of the *ECT2* promoter.** 9-day-old primary transformants expressing ECT1, ECT2 or ECT4 fused to mCherry in the *te234* background. All transgenes and expressed from gDNA under the control of the *ECT2* promoter. mCherry fluorescence on the right panels show the variability in transgene expression levels with this setup. Additional genotypes in the top panels are shown as a reference. Scale bars are 0.5 mm.

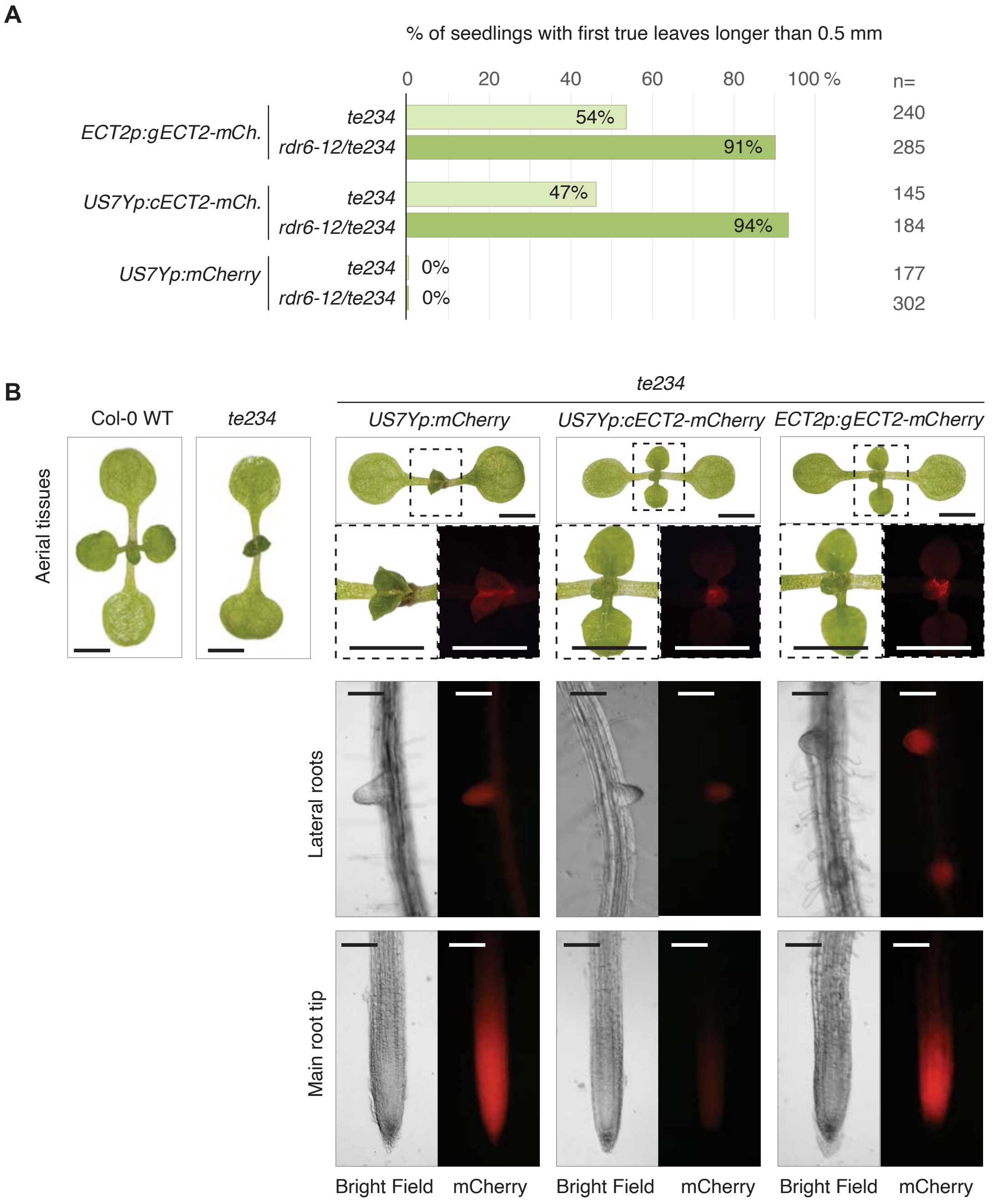


**S6 Fig. Expression of *cECT2-mCherry* driven by the ribosomal protein *uS7Y* promoter can rescue late leaf emergence in *ect2-1/ect3-1/ect4-2* (*te234*) seedlings. (A).** Raw percentages of seedlings with first true leaves >0.5 mm at 10 days after germination among primary transformants of *te234* or *rdr6-12/te234* plants transformed with *uS7Yp:cECT2-mCherry-OCSt* (cDNA) or *uS7Yp:mCherry-OCSt* (control) compared to *ECT2p:gECT2-mCherry-ECT2t* (gDNA). **(B)** Expression pattern in 9-day-old seedlings of the indicated genotypes (T2 for *US7Yp*-driven constructs, and T5 for *ECT2p:gECT2-mCherry-ECT2t*). Fluorescence and protein abundance is typically higher in plants expressing free mCherry (*US7Yp:mCherry-OCSt*) than in fusions of mCherry with ECTs (see S8 and S15 Figs). Although the expression pattern of *US7Yp:cECT2-mCherry-OCSt* is identical to that of *ECT2p:gECT2-mCherry-ECT2t*, the fluorescence level and protein abundance observed is typically lower among cDNA lines in the *te234* background. Scale bars are 1 mm for images of aerial tissues (upper panels), and 0.1 mm for roots (lower panels).

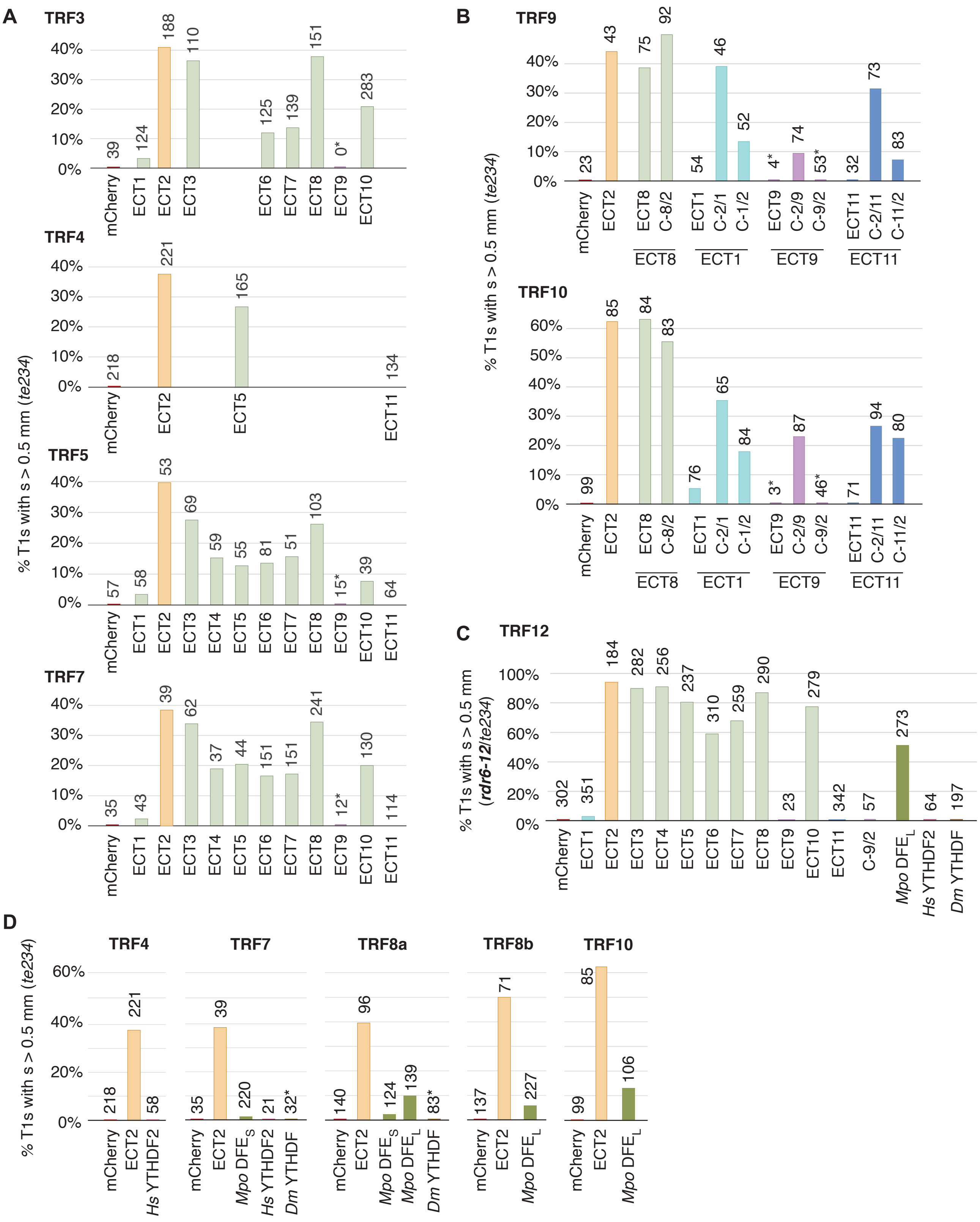


**S7 Fig. Complementation rates (raw data)**. **(A-D)** Absolute complementation rates in each independent transformation in *te234* (A,B,D) or *rdr6-12/te234* (C) plants given by the percentage of primary transformants (T1s) with first true leaves bigger than 0.5 mm at 10 days after germination (DAG) for *uS7Bp:cECT(X)-mCherry-OCSt* (A, C), *uS7Bp:cECT(X)*$_{IDR}$*/cECT(Y)*$_{YTH}$*-mCherry-OCSt* chimeras (B, C), or *uS7Bp:cYTHDF(X)-mCherry-OCSt* heterologous constructs (C,D). Numbers over the bars indicate the total number of transformants. Asterisks indicate absence of mCherry fluorescence (expression) among all retrieved transformants, or very faint fluorescence that underwent silencing in the next generation (T2). S6 Dataset contains detailed T1 counts and the calculation of complementation rates.

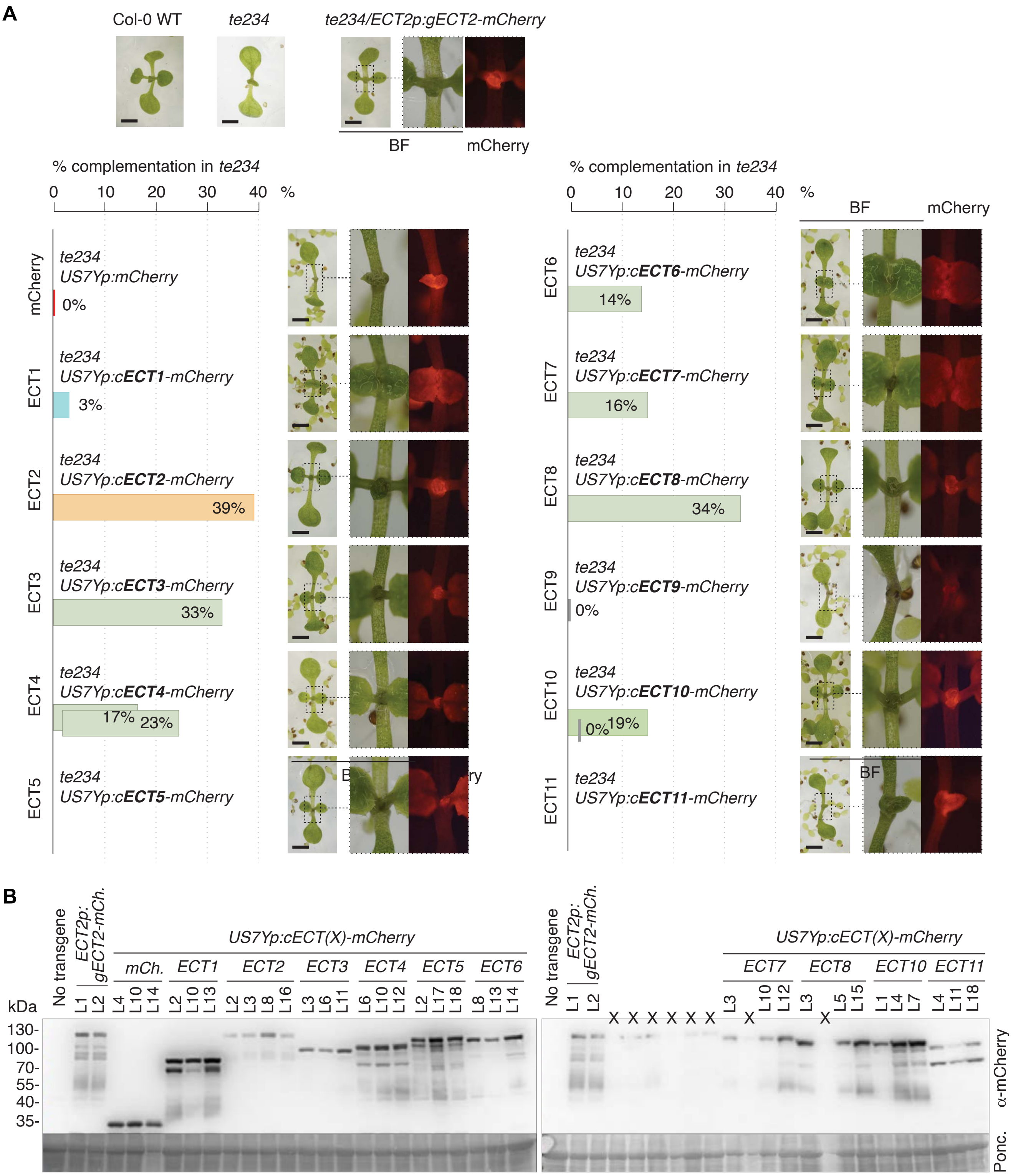


**S8 Fig. Expression of *US7Yp:cECT(X)-mCherry-OCSt* constructs in *te234* plants. (A)** Representative 10-day-old *te234/US7Yp:cECT(X)-mCherry-OCSt* T1 seedlings with their fluorescent signal and genetic background controls. Bars represent weighed averages of complementation over 2-5 transformations (Fig 2C). Scale bars are 1 mm. **(B)** Levels of protein expression from *US7Yp:cECT(X)-mCherry-OCSt* constructs in independent lines (L) of 9-day-old T2 seedlings assessed by α-mCherry western blot. All genotypes are in the *te234* background. Lines with single insertions and the highest complementation capacity (largest true leaves at 9 days after germination in T2) were selected for the analysis. ECT9 is not included because we could not observe fluorescence in the T2 generation for any of the few lines obtained due to silencing. *te234 ECT2p:gECT2-mCherry-ECT2t* lines [16] are included in both membranes as a reference. Lanes marked with X in the lower membrane correspond to either lines irrelevant for this study, or left empty due to defects in the wells. Ponceau-staining is shown as loading control.

| | *te234* | | | | | | | | | *rdr6-12 te234* |
|---|---|---|---|---|---|---|---|---|---|---|
| **T1s per plate (~2000 seeds)** | **TRF1** | **TRF3** | **TRF4** | **TRF5** | **TRF7** | **TRF8a** | **TRF8b** | **TRF9** | **TRF10** | **TRF12** |
| **mCherry** | 35.0 | 6.5 | 36.3 | 9.5 | 5.8 | 23.3 | 22.8 | 3.8 | 16.5 | 50.3 |
| **ECT1** | | 20.7 | | 9.7 | 7.2 | | | 9.0 | 12.7 | 58.5 |
| **ECT2** | 37.0 | 31.3 | 36.8 | 8.8 | 6.5 | 16.0 | 11.8 | 7.2 | 14.2 | 30.7 |
| **ECT3** | | 18.3 | | 11.5 | 10.3 | | | | | 56.4 |
| **ECT4** | | | | 9.8 | 6.2 | | | | | 42.7 |
| **ECT5** | | | 27.5 | 9.2 | 7.3 | | | | | 39.5 |
| **ECT6** | | 20.8 | | 13.5 | 25.2 | | | | | 51.7 |
| **ECT7** | | 23.2 | | 8.5 | 25.2 | | | | | 51.8 |
| **ECT8** | | 25.2 | | 17.2 | 40.2 | | | 12.5 | 14.0 | 48.3 |
| **ECT9** | | 0.0 | | 2.5 | 2.0 | | | 0.7 | 0.5 | 4.6 |
| **ECT10** | | 47.2 | | 6.5 | 21.7 | | | | | 69.8 |
| **ECT11** | | | 22.3 | 10.7 | 19.0 | | | 5.3 | 11.8 | 57.0 |
| **C-1/2** | | | | | | | | 8.7 | 14.0 | |
| **C-2/1** | | | | | | | | 7.7 | 10.8 | |
| **C-9/2** | | | | | | | | 8.8 | 7.7 | 20.5 |
| **C-2/9** | | | | | | | | 12.3 | 14.5 | |
| **C-11/2** | | | | | | | | 13.8 | 13.3 | |
| **C-2/11** | | | | | | | | 12.2 | 15.7 | |
| **C-8/2** | | | | | | | | 15.3 | 13.8 | |
| ***Mpo* DFE$_S$** | | | | | 36.7 | 20.7 | | | | |
| ***Mpo* DFE$_L$** | | | | | | 23.2 | 18.9 | | | 45.5 |
| ***Hs* YTHDF2** | | | 9.7 | | 3.5 | | | | | 10.7 |
| ***Dm* YTHDF** | | | | | 5.3 | 13.8 | | | | 32.8 |

Transformation efficiency (average) *te234:* 15.3 T1s / plate
*rdr6-12/te234:* 41.9 T1s / plate

**S9 Fig. Transformation Efficiency.** Estimated transformation efficienty (average of 6 plates) of each construct for all the independent transformation batches of *te234* (TRF1-10) or *rdr6-12/te234* (TRF12) plants used in this study. TRF8 was screened twice (TRF8a and TRF8b). Red-to-blue colouring in the *te234* background reflect the variability in transformation efficiencies across constructs and batches, to highlight the consistently low recovery of transformants for *Ath* ECT9 and *Hs* YTHDF2.

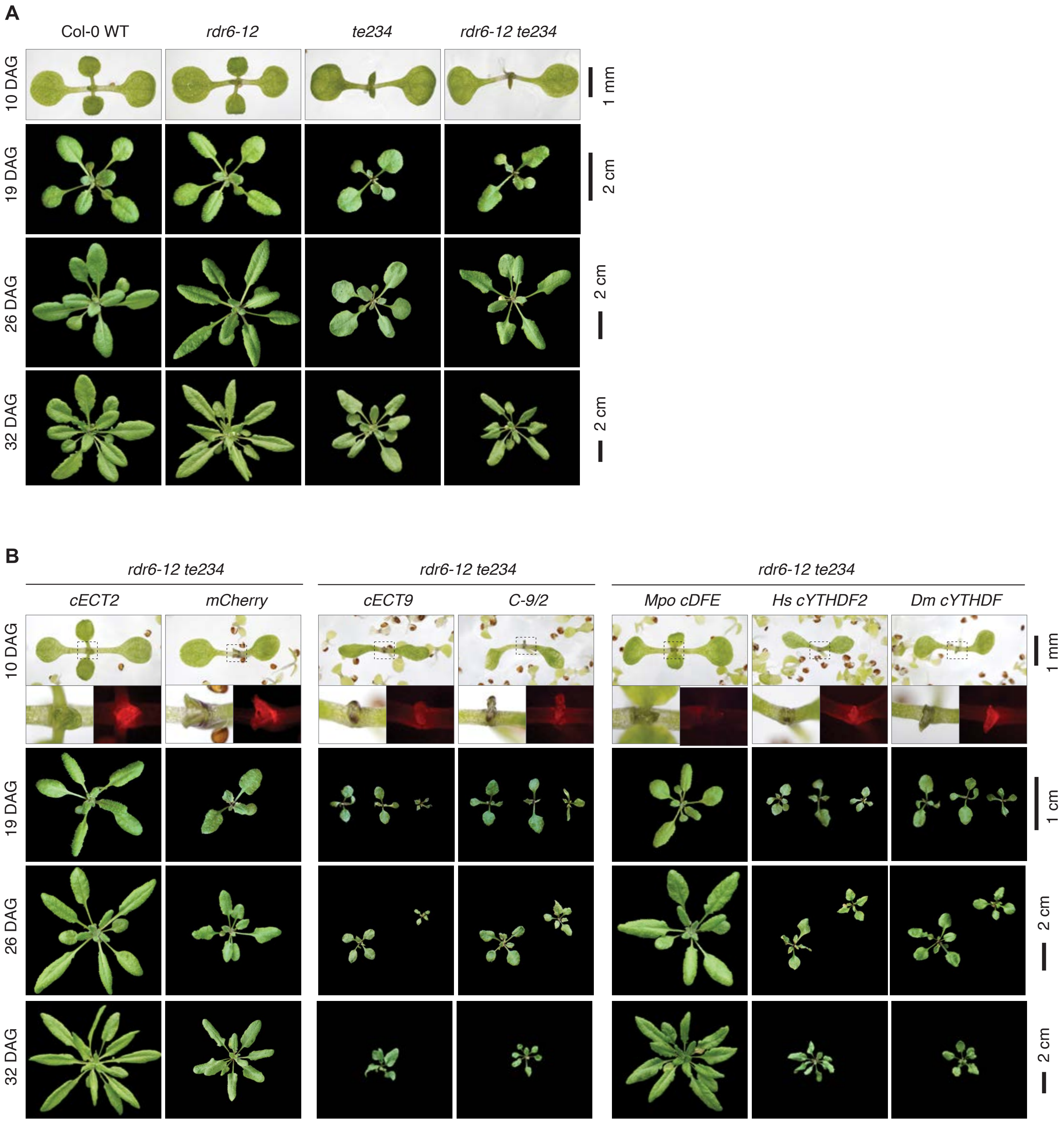


**S10 Fig. Phenotypic characterization of selected transgenic lines in *rdr6-12/te234* and relevant genetic backgrounds. (A)** Developmental stages of plants of the indicated genotypes. DAG, days after germination. **(B)** Same as in A for primary transformants (T1s) of the indicating transgenes, all expressed from the *US7Y* promoter with mCherry fused at the C-terminus. Dashed outlines at 10 DAG are magnified below each panel to show mCherry fluorescence. Several different independent lines of the same age are shown for the genotypes exhibiting the strongest developmental defects. Plants in A and B were grown in parallel.

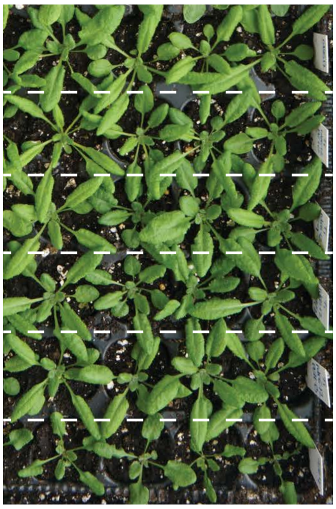


**S11 Fig. Expression of *US7Yp:cECT9-mCherry-OCSt* in the *rdr6-12* background has no obvious phenotypic effect.** Rosettes of 24-day-old primary transformants expressing the indicated transgenes in the *rdr6-12* background, grown in parallel with *rdr6-12* and *rdr6-12/te234* controls.

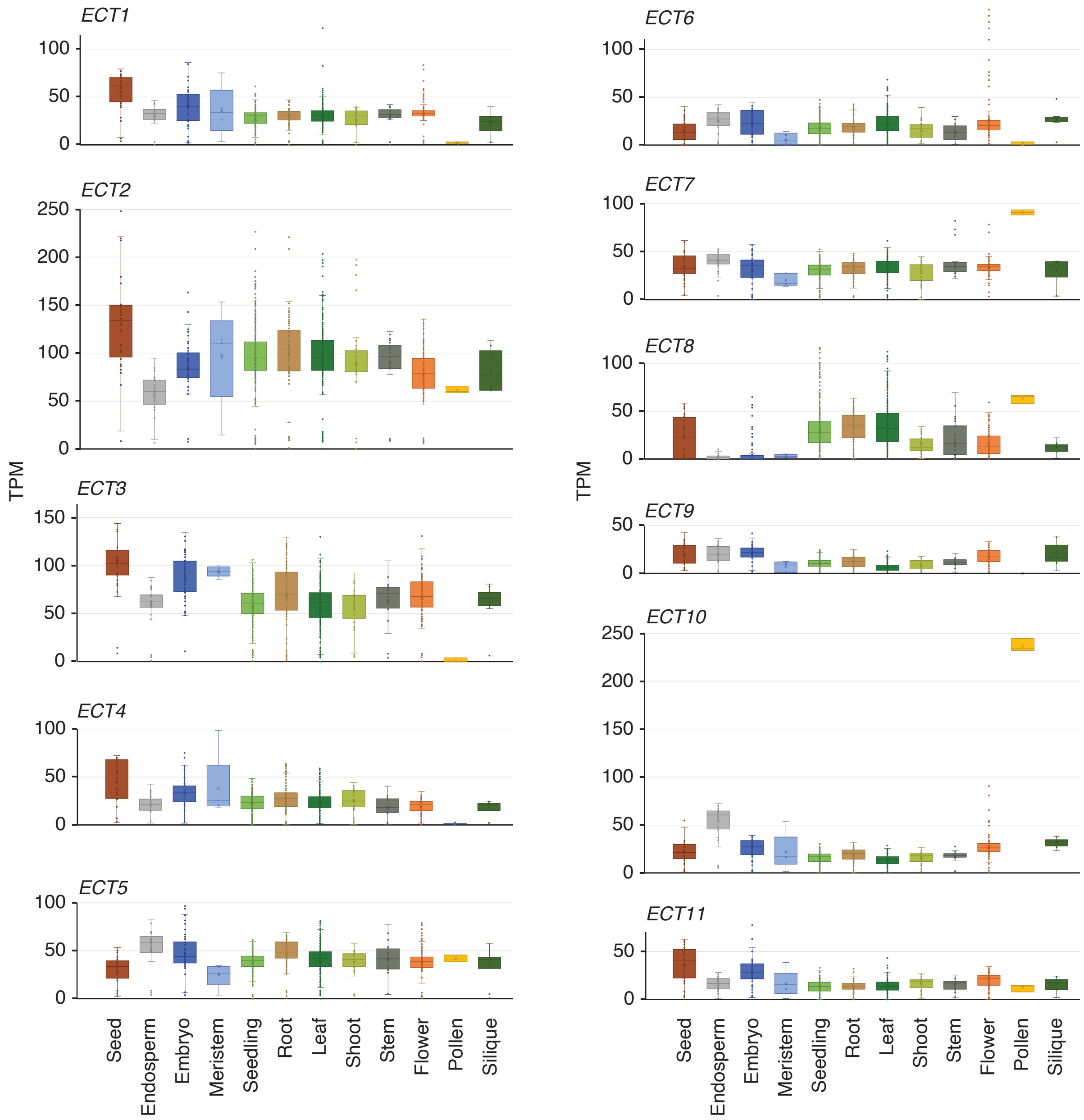


**S12 Fig. Expression levels of *ECT* paralogs in tissues of *Arabidopsis thaliana*.** Expression levels of *ECT1-ECT11* across different tissues according to public mRNA-Seq data [72]. TPM, transcripts per million.

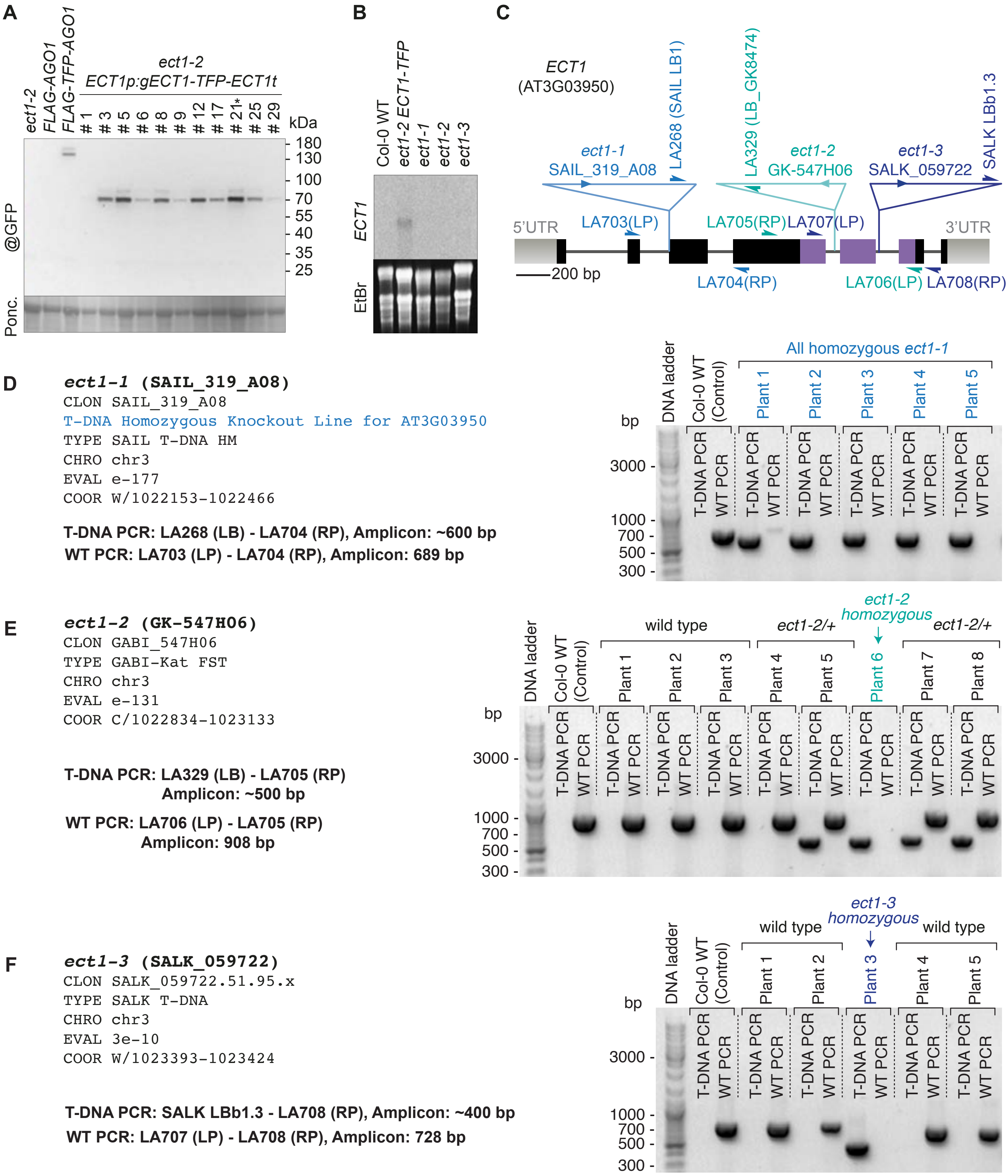


**S13 Fig. Isolation of ECT1-TFP transgenic lines and ect1 T-DNA insertion alleles. (A)** Western blot using antibodies against GFP (that recognize TFP) in different *ECT1p:gECT1-TFP-ECT1t* independent lines. Ponceau (Ponc.) staining of the membrane is used as loading control. **(B)** Northern blot using the probe (P) specified in Fig 3G to detect *ECT1* mRNA. Although the probe recognizes specifically *ECT1* in the *ECT1p:gECT1-TFP-ECT1t* Line #21 (marked with an asterisk in A), the endogenous expression levels of *ECT1* in Col-0 wild type are below detection limit by northern blot. **(C)** Schematic representation of the *Ath ECT1* locus (At3g03950). Exons are represented as boxes and introns as lines. Untranslated regions (UTRs) are coloured grey, the sequence encoding the YTH domain is purple, and the rest of the *ECT1* coding sequence is black. The IDs and positions of the T-DNA insertions assigned to *ect1-1*, *ect1-2* and *ect1-3* alleles are marked, and so is the location of primers used for their genotyping. **(D-F)** 1% agarose gels showing EtBr-stained PCR fragments corresponding to the genotyping of *ect1-1* (D), *ect1-2* (E) and *ect1-3* (F) in plants germinated from seeds provided by the Nottingham Arabidopsis Stock Center (NASC) as indicated. The primer set for each PCR ('T-DNA' detects the insertion, and 'WT' detects the wild type allele) and the length of the resulting amplicons are indicated to the left of each gel. The sequence of all primers can be found in S2 Table. The progeny of plants homozygous for each T-DNA insertion, highlighted in shades of blue, was selected for crosses and further characterization.

**A**

| Short name | Allele combination | First described |
|---|---|---|
| *Gde12* | *ect1-2 ect2-3* | This work |
| *Gde23* | *ect2-3 ect3-2* | Arribas-Hernández et al., 2018. The Plant Cell |
| *Gte123* | *ect1-2 ect2-3 ect3-2* | This work |
| *Gte234* | *ect2-3 ect3-2 ect4-2* | Arribas-Hernández et al., 2020. Development |
| *Gqe1234* | *ect1-2 ect2-3 ect3-2 ect4-2* | This work |
| *te234* | *ect2-1 ect3-1 ect4-2* | Arribas-Hernández et al., 2018. The Plant Cell |
| *qe1234* | *ect1-1 ect2-3 ect3-2 ect4-2* | This work |

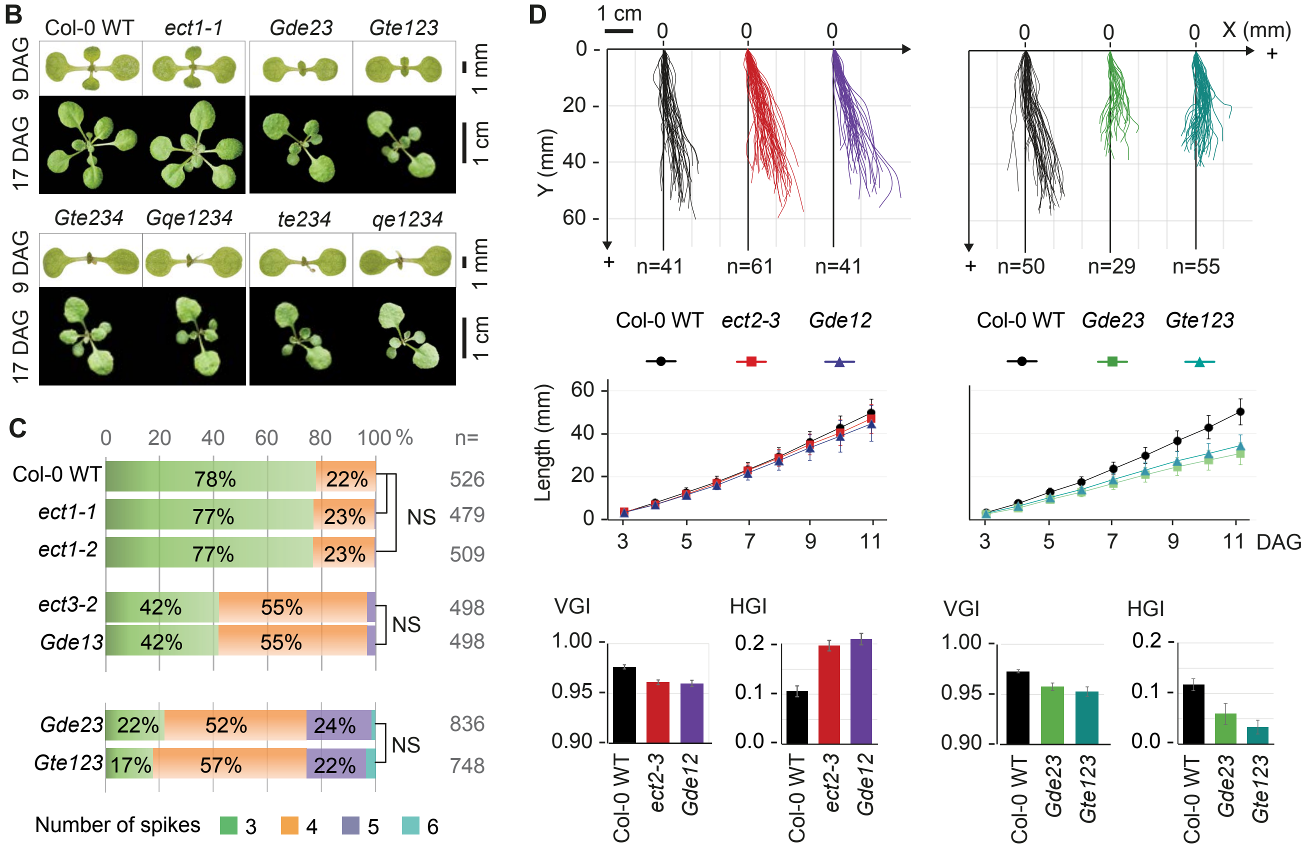


**S14 Fig. Knockout of ECT1 does not affect arabidopsis plant development. (A)** Abbreviations for combinations of double (d), triple (t), or quadruple (q) ect (e) mutants following the nomenclature proposed by Arribas-Hernández et al. [17]. The prefix 'G' identifies allele combinations having only T-DNA insertions belonging to the GABI-KAT collection [127]. **(B)** Morphological appearance of seedlings with or without *ECT1* in the different backgrounds indicated at 9 or 17 days after germination (DAG). **(C)** Analyses of the percentage of trichomes with 3, 4, 5 or 6 spikes in plants with or without *ECT1* in the different backgrounds indicated. n, number of trichomes assessed for each genotype. NS, non-significant differences according to statistical analysis performed as in [16]. **(D)** Analysis of root growth rate and directionality upon mutation of *ECT1* in the *ect2-3* and *ect2-3/ect3-2* (*Gde23*) knockout backgrounds. Upper panels represent the overlayed silhouettes of actual roots as they grow on vertically disposed MS-agar plates. n, number of plants assessed for each genotype. VGI, vertical growth index; HGI, horizontal growth index. Root phenotypic analyses were conducted as in [17].

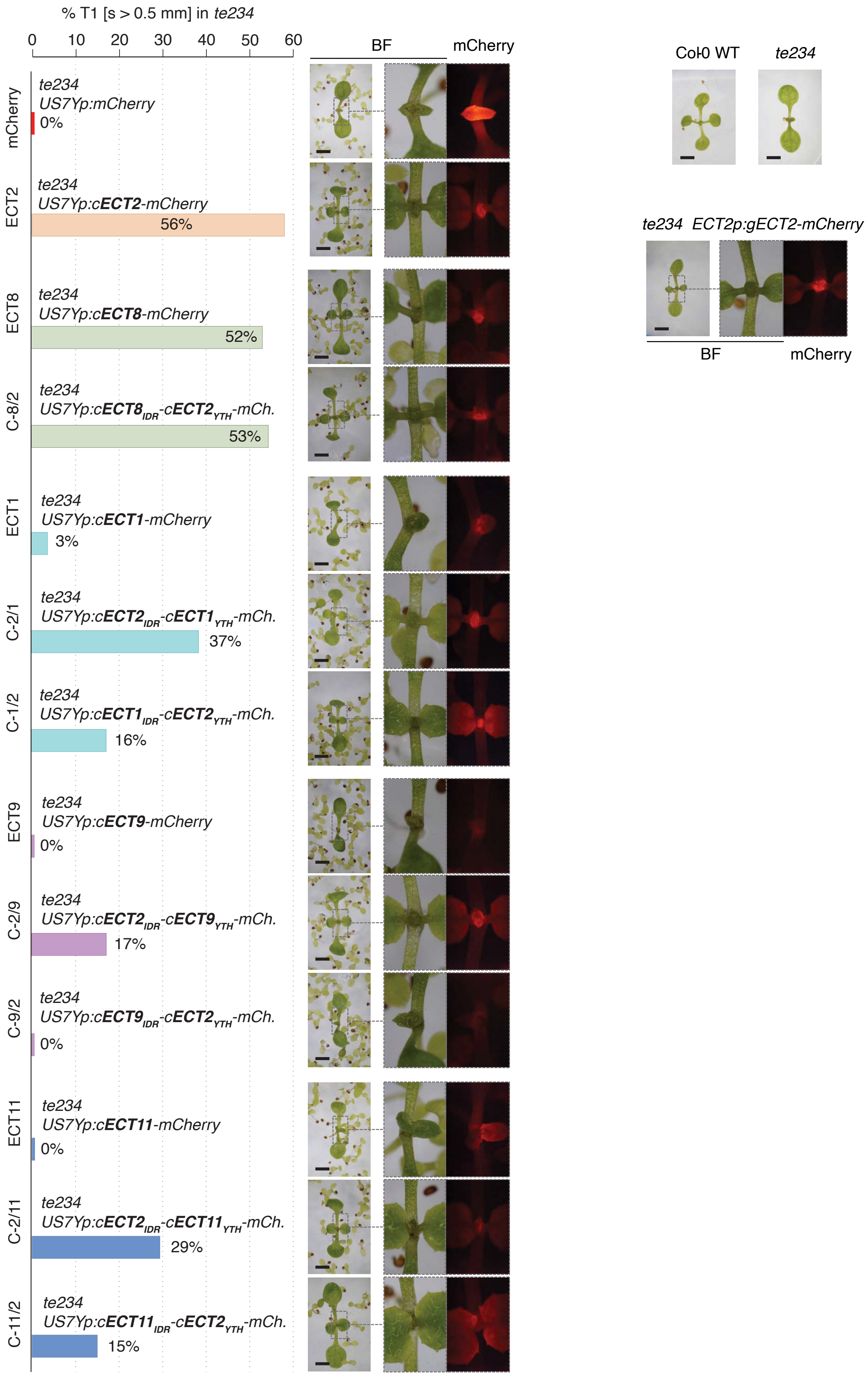


**S15 Fig. Expression of chimeric *ECT(X)*$_{IDR}$*-ECT(Y)*$_{YTH}$ constructs in *te234* plants.** Representative 10-day-old *te234/uS7Yp:cECT(X)*$_{IDR}$*-cECT(Y)*$_{YTH}$*-mCherry-OCSt* T1 seedlings with their fluorescent signal. Bars represent weighed averages of complementation. Scale bars are 1 mm.

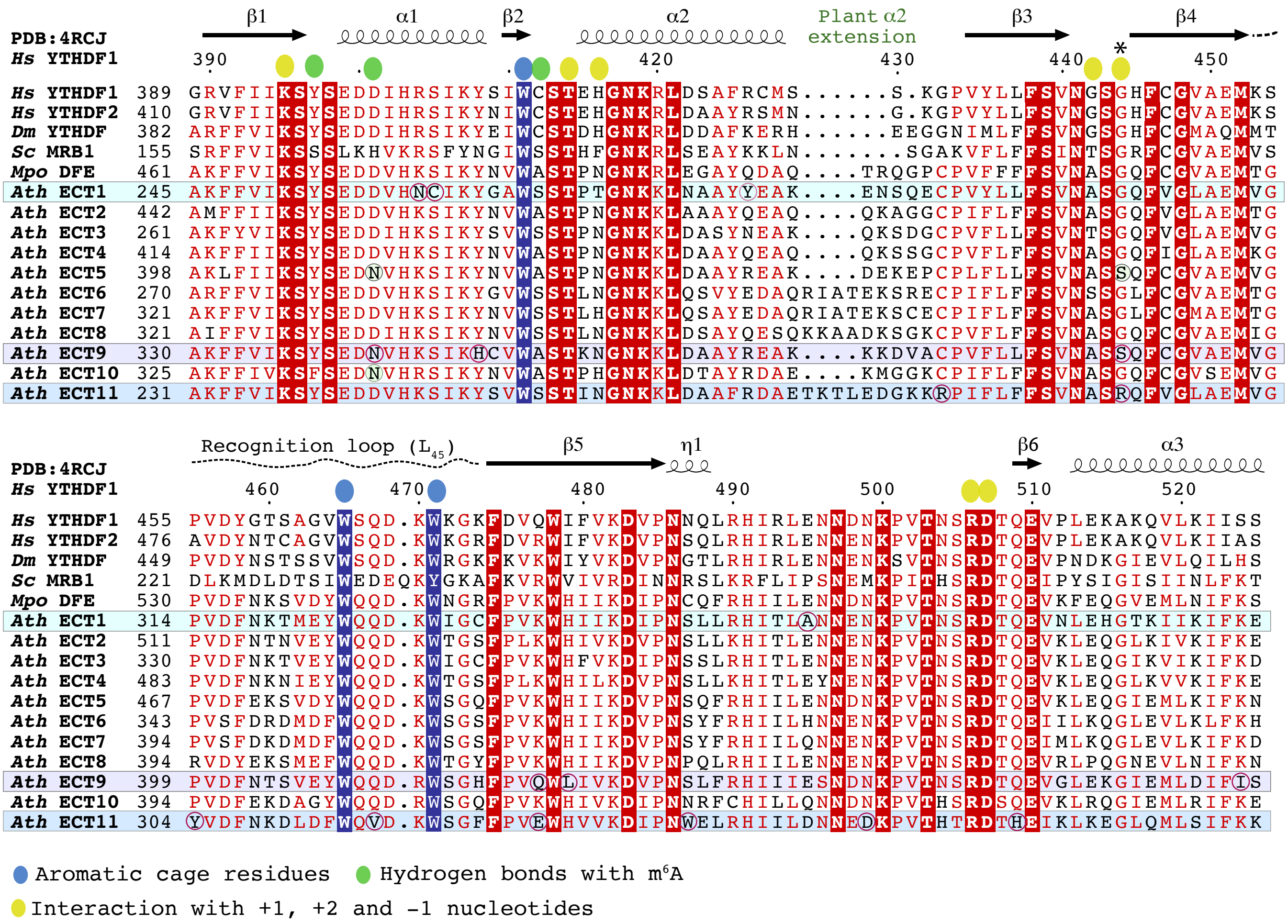


**S16 Fig. Conservation of the YTH domain at the sequence level.** Sequence alignment of the YTH domains of *Arabidopsis thaliana* (*Ath*) ECT1-11, *Marchantia polymorpha* (*Mpo*) DFE, *Saccharomyces cerevisiae* (*Sc*) MRB1/Pho92, *Drosophila melanogaster* (*Dm*) YTHDF and *Homo sapiens* (*Hs*) YTHDF2 and YTHDF1. The alignment is colored and annotated jusing ESPript 3 [127], according to the crystal structure of *Hs* YTHDF1 in complex with RNA GGm6ACU [61] as template for structural elements. The residues of Hs YTHDF1 that have contacts with RNA are marked above the sequences.

Positions in *Ath* ECT1, ECT9 and ECT11 with non-conservative changes compared to the majority of ECTs are highlighted. Most of these substitutions are located on the protein surface according to 3D homology models of the ECT1/9/11 YTH domains (S17 Fig) except for the following residues: For **ECT1**, N260 and C261 (α1) are in close proximity to the aromatic cage. For **ECT9**, N432 (α1) forms hydrogen bonds with $m^6A$. The D to N substitution at that position in all members of the DF-B clade (marked also in ECT5 and ECT10) [26], fern DF-Fs (S4 Fig) and YTHDC proteins [13, 61] increases the affinity for $m^6A$ by 15-fold [61] and is likely the product of convergent evolution. The remaining ECT9 substitutions are on the surface of the protein in areas not known to interact with RNA except for S388 (*), also a serine in the DF-B member ECT5 (marked) despite the presence of glycine in this position in most YTHDC and YTHDF proteins of plants, animals and fungi. The backbone NH group of this glycine in *Hs* YTHDF1 (Gly444) and *Zygosaccharomyces rouxii* (*Zro*) MRB1 (Gly233) forms hydrogen bonds, through a water molecule, with the phosphate backbone of the RNA [11, 61]. Remarkably, **ECT11** contains an arginine (R293) in the same position. Other ECT11 substitutions are on the surface and far from the RNA-binding groove.

Of note, plant YTHDF proteins have a small insertion between α2 and β3 that extends α2 further out than in the metazoan orthologs. This insertion is slightly longer the in the angiosperm DF-D and -C clades, which include *Ath* ECT11 (S17 Fig), as well as in all members of the fern/gymnosperm-exclusive DF-F clade (S4 Fig), and in many bryophyte and lycophyte DF-Es (S4 Fig).

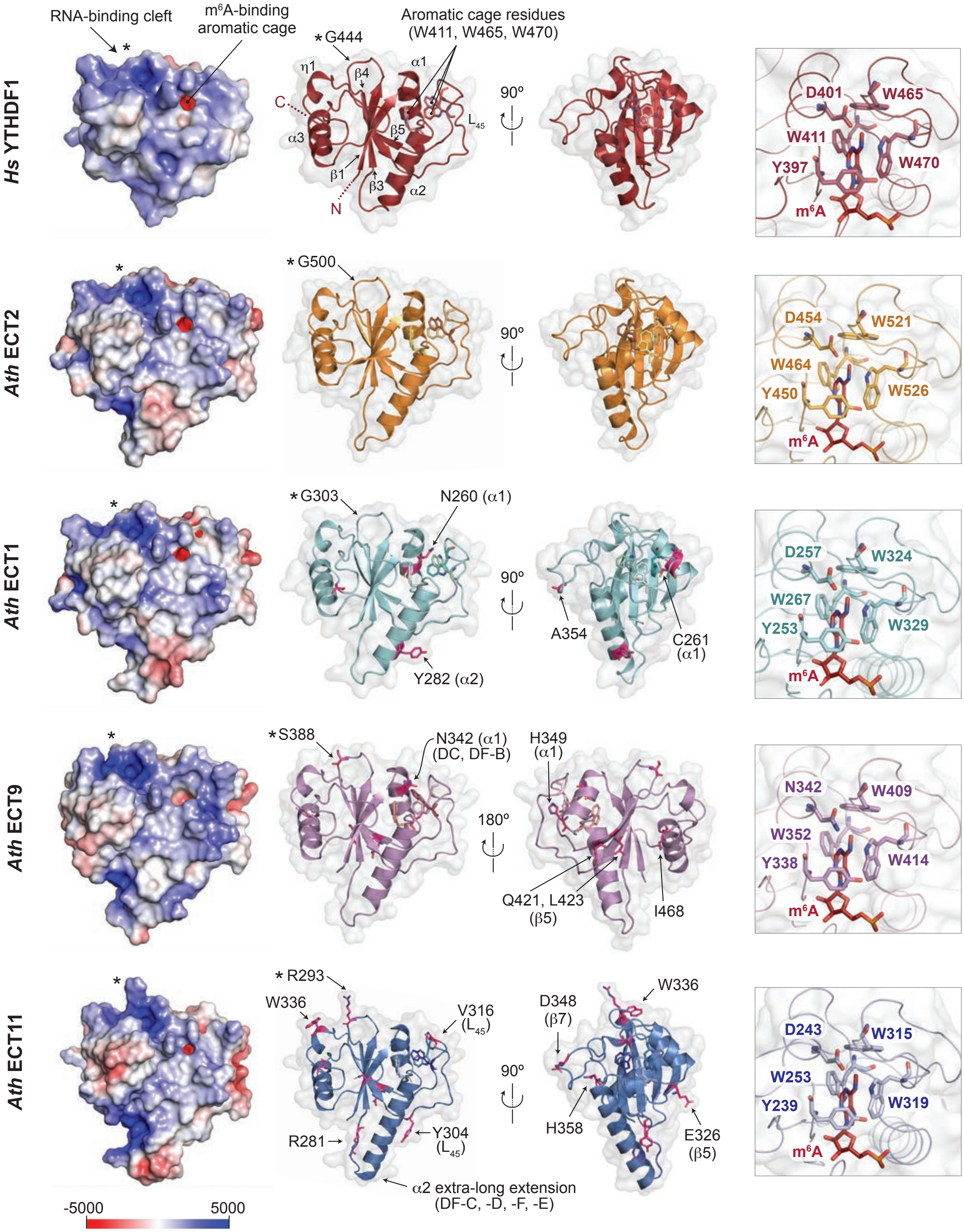


**S17 Fig. Conservation of the YTH domain at the structural level.** Three left panels, AlphaFold-predicted [73, 74] apo structures of YTH domains of *Arabidopsis thaliana* (*Ath*) ECT1/9/11 in comparison to that of *Ath* ECT2, and of *Homo sapiens* (*Hs*) YTHDF1. Red/blue coloring in the left-most panel indicates the electrostatic potential of the surface calculated using the APBS PyMOL plugin [128] with the parameters described in Methods. The two middle panels show the similarity of the secondary structure of all proteins and highlight the few non-conservative amino acid substitutions identified in *Ath* ECT1/9/11 (S16 Fig). Right panel, hydrophobic cage of the same proteins modelled on the crystal structure of the YTH domain of *Hs* YTHDF1 in complex with RNA GGm$^6$ACU [61] (Protein Data Bank entry 4RCJ). The key residues for m$^6$A binding are marked, including the aspartic to asparagine substitution in the DF-B clade member *Ath* ECT9 (N342).

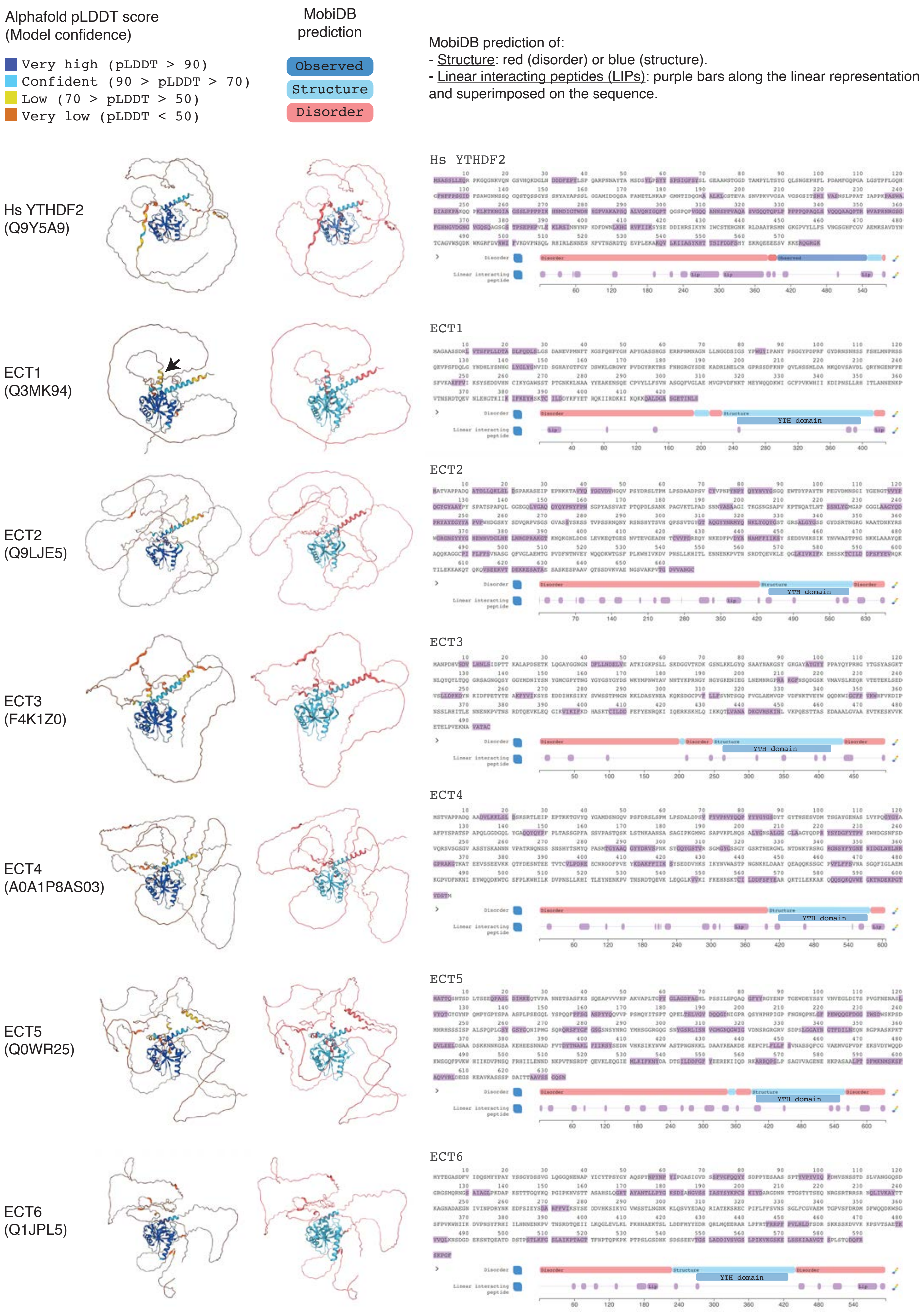


S18 Fig, continues on the next page

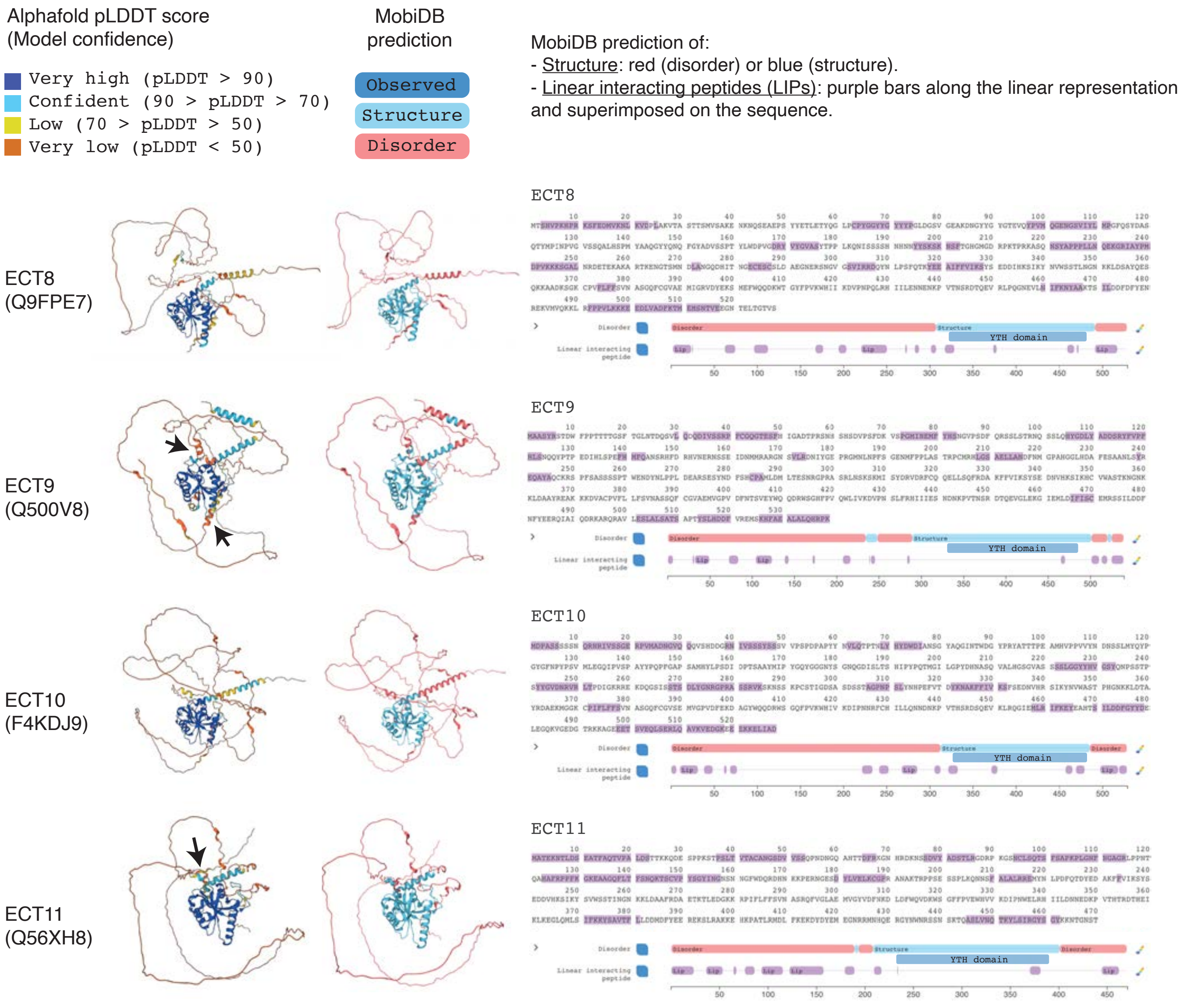


**S18 Fig. Predicted structure and short linear motifs (SLiMs) of arabidopsis ECTs compared to human YTHDF2. Left columns:** Structure of *Arabidopsis thaliana* (*Ath*) ECT proteins predicted by AlphaFold [73, 74] colored according to a confidence score (pLDDT) given per-residue between 0 and 100. Regions below 50 pLDDT may be unstructured in isolation [74, 129]. The vast majority of N-terminal regions in ECT proteins score below 50 pLDDT (dark orange). A few short stretches with low-confidence prediction of secondary structure are marked with black arrows for ECT1/9/11. Middle column: Disorder prediction by MobiDB [76] depicted by red (disorder) or light blue (structure) coloring of the AlphaFold-predicted structure. The N-terminal regions of all ECT proteins are predicted to be largely disordered (red). **Right columns**, sequences and linear representation of the proteins indicating MobiDB [76] disorder predictions ('Disorder' track) with the same colors as in the middle column. The known structure of the YTH domain of *Homo sapiens* (*Hs*) YTHDF2 is marked in dark blue, and the YTH domains of *Ath* ECTs, with no experimentally resolved structures to date, have been manually added also in dark blue. Purple bars superimposed on the amino acid sequences of the proteins and purple boxes positioned along the linear representations on the 'Linear interacting peptide' (LIP) tracks highlight residues predicted to interact with another molecule that preserve structural linearity in the bound state. In addition to LIPs, they are generally called short linear motifs (SLiMs), molecular recognition features (MoRFs) or protean segments (ProS). The UniProt ID of all proteins is given below their names.

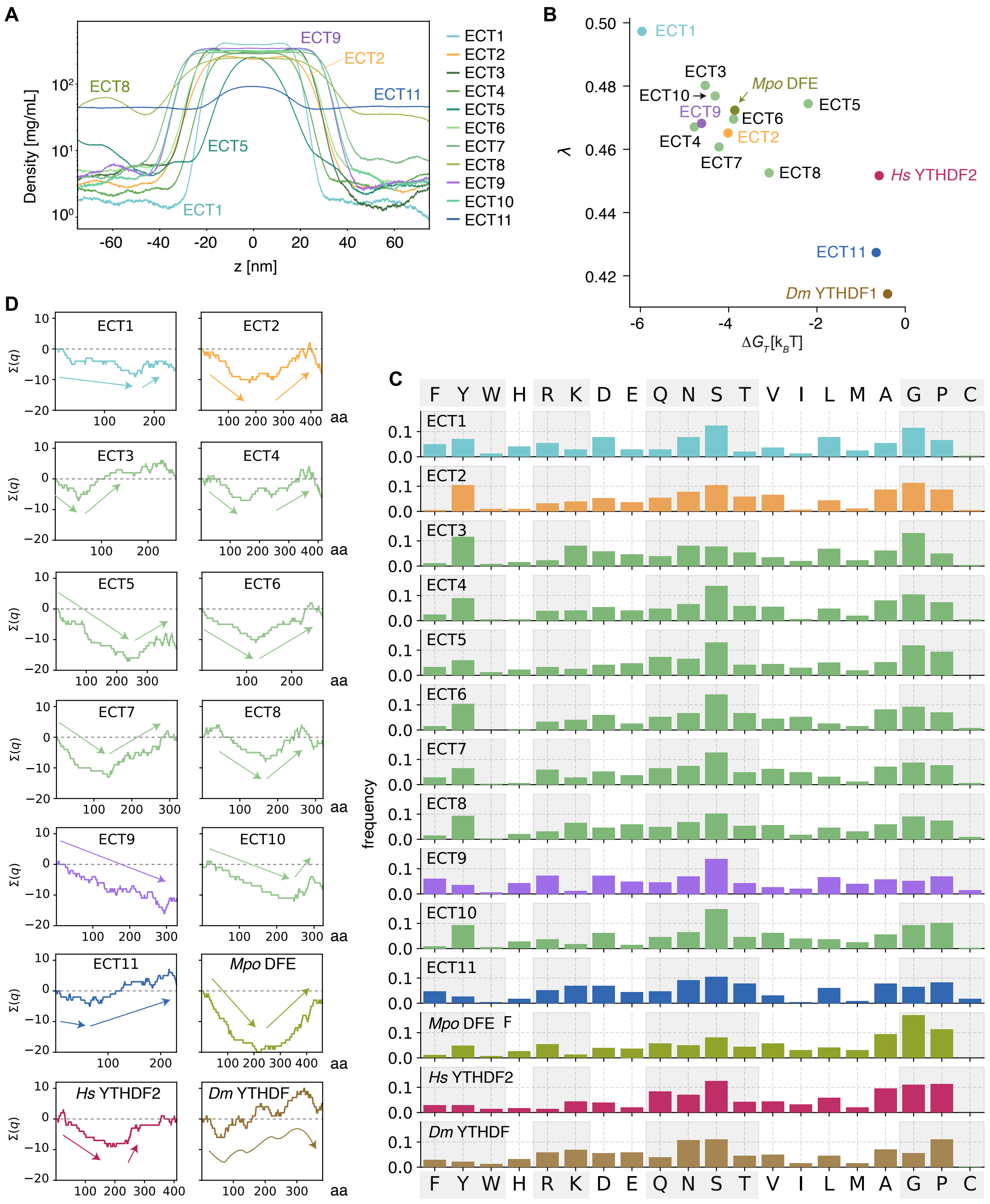


**S19 Fig. Analysis of the IDRs of the YTHDF proteins assayed in this study (extended data). (A)** Time-averaged density profiles of the slab simulations of the ECT proteins shown in Fig 5B. **(B)** Relationship between excess transfer free energy from dilute to dense phase ($\Delta G_{trans}$ = RT ln ($c_{dilute}$ / $c_{dense}$)) and average stickiness ($\lambda$) of the IDR residues. **(C-D)** Comparative amino acid composition (C) and representation of charges (D) along the IDR sequences of the different YTHDF proteins assayed in this study. In D, arrows highlight the overall trends in charge change along the IDRs. Notice that the length in amino acids (aa) is not at scale to simplify the representation.

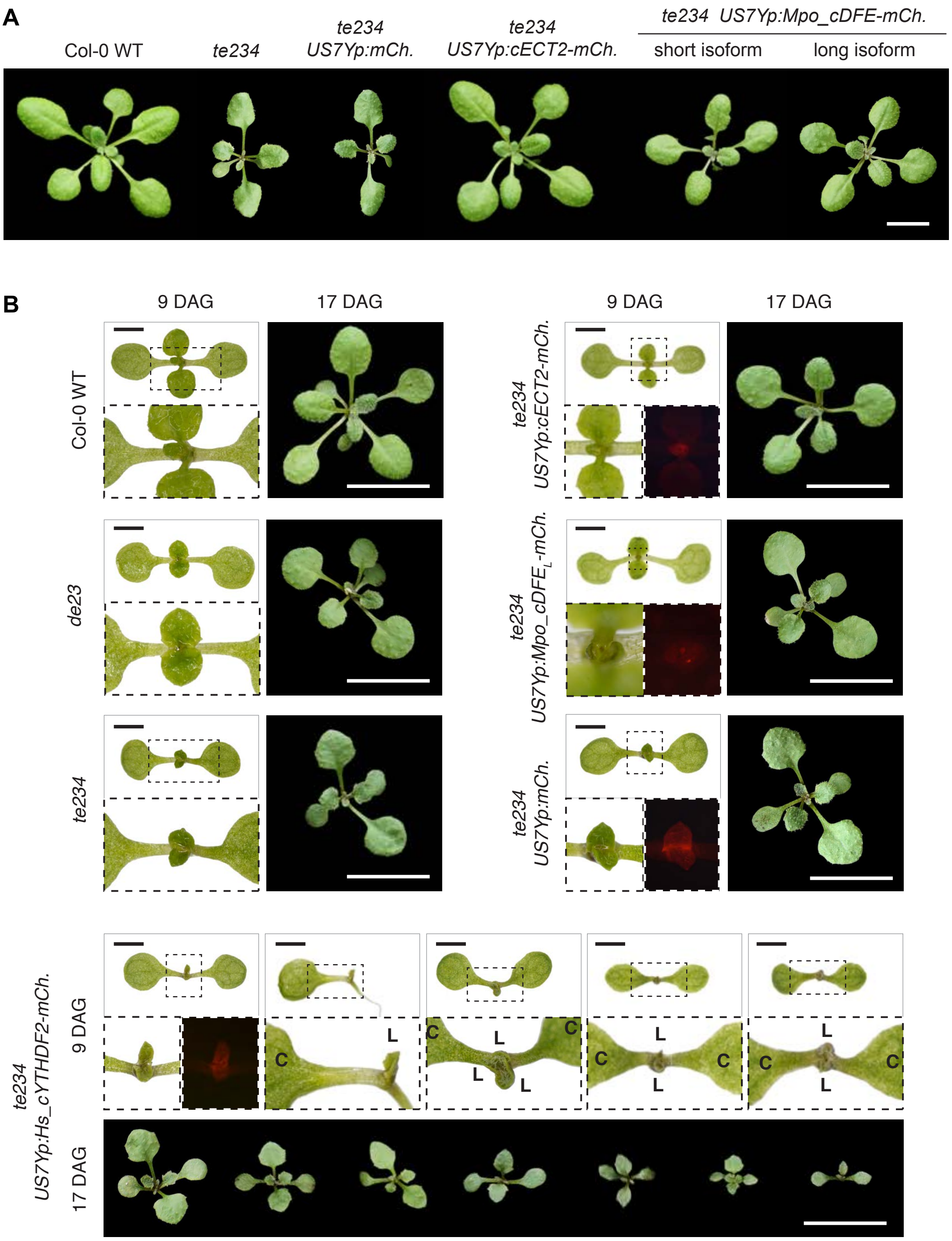


**S20 Fig. Heterologous expression of bryophyte and metazoan YTHDF proteins in arabidopsis (extended data). (A)** T1 plants expressing the long or short splice forms of *Marchantia polymorpha* (*Mpo*) DFE 19 days after germination (DAG). Additional genotypes are included as a reference. **(B)** Morphological appearance and fluorescence pattern of T2 plants expressing *Mpo* $DFE_L$ (L, long isoform) or *Homo sapiens* (*Hs*) YTHDF2 in the *te234* background 9 or 17 DAG. Additional genotypes are included as a reference. For *Hs* YTHDF2, several plants are shown to reflect the variability in phenotypic defects, which include enhanced aberrance in the shape or number of cotyledons and first true leaves, and slower overall growth. C, cotyledons; L, true leaves. White scale bars are 1 cm, and black are 1 mm.

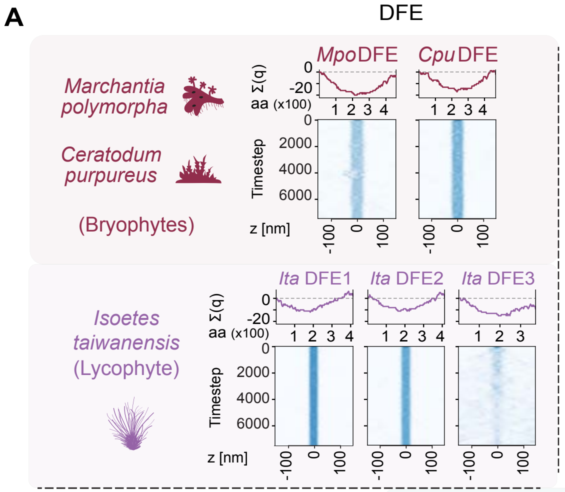


**S21 Fig. Analysis of the IDRs of YTHDF proteins from representative species of the main taxa of land plants. (A)** Representation of charges (top panels) and slab simulations (bottom panels) of the IDRs of YTHDF proteins from the main taxa of land plants. The proteins are sorted and the charge distributions coloured according to species and land plant evolution from top (basal group) to bottom, and the different plant DF clades are separated along the horizontal axis. *Isoetes taiwanensis* (*Ita*) DF-CD (Fig 1B) is excluded from the analysis because its N-terminus is very short (S1 Dataset) and does not behave like an IDR. **(B)** Relationship between excess transfer free energy from dilute to dense phase ($\Delta G_{trans}$ = RT ln [$c_{dilute}$ / $c_{dense}$]) and average stickiness (λ) of the IDR residues of the proteins in A. The proteins are color-coded according to plant DF clades as indicated. *Homo sapiens* (*Hs*) YTHDF2 is included as a reference. Same as for the *Arabidopsis thaliana* (*Ath*) ECT set in S19B Fig, the content of sticky residues defined by the CALVADOS model [78] mainly governs the different phase separation propensity among the IDRs. A clade-dependent separation of the proteins according to these properties is apparent.

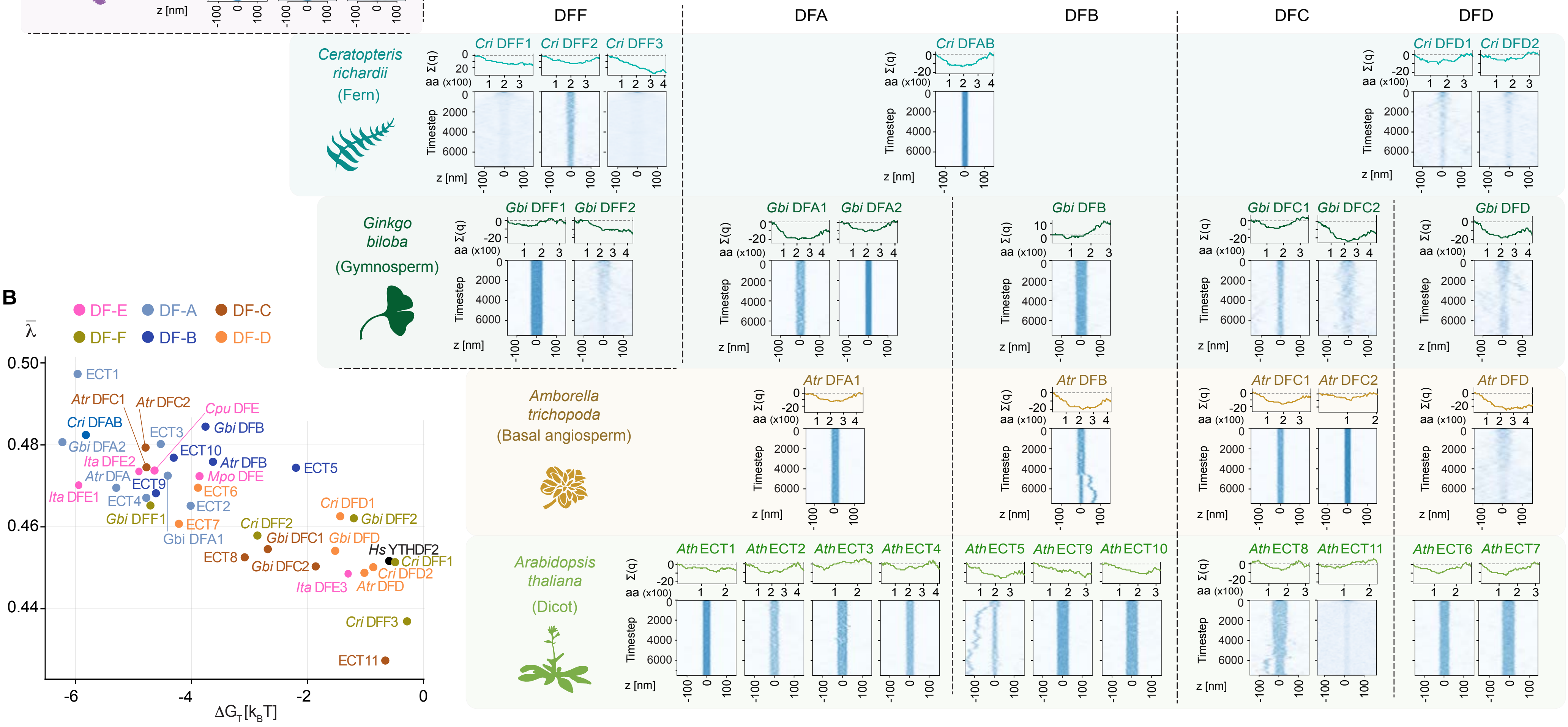

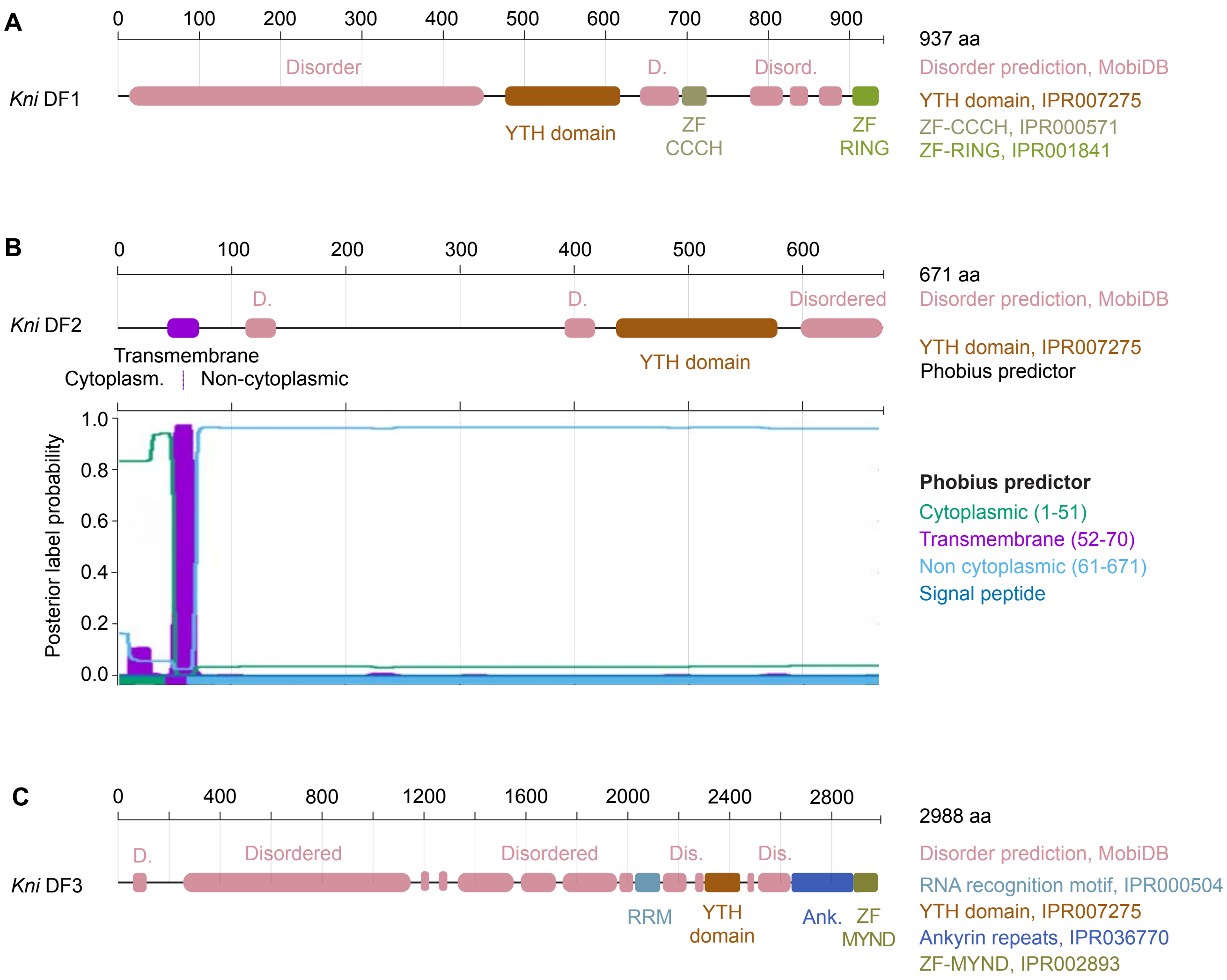


**S22 Fig. Analysis of domain composition of YTHDF proteins from the charophyte *Klebsormidium nitens* (*Kni*). (A-C)** Prediction of domains and other features in the protein sequences of *Kni* DF1-3 according to InterPro (former Pfam) (https://www.ebi.ac.uk/interpro/) [130]. Horizontal axes indicate protein length in amino acids (aa). InterPro entries (unique protein homologous superfamilies, families, domains, repeats or important sites based on one or more signatures) are indicated on the right side. Disorder prediction is engineered by MobiDB (https://mobidb.bio.unipd.it/) [76], and transmembrane domains are predicted by Phobius (https://phobius.sbc.su.se/) [131]. For *Kni* DF2 in B, the direct graphical output of Phobius Transmembrane Topology Predictor is shown.

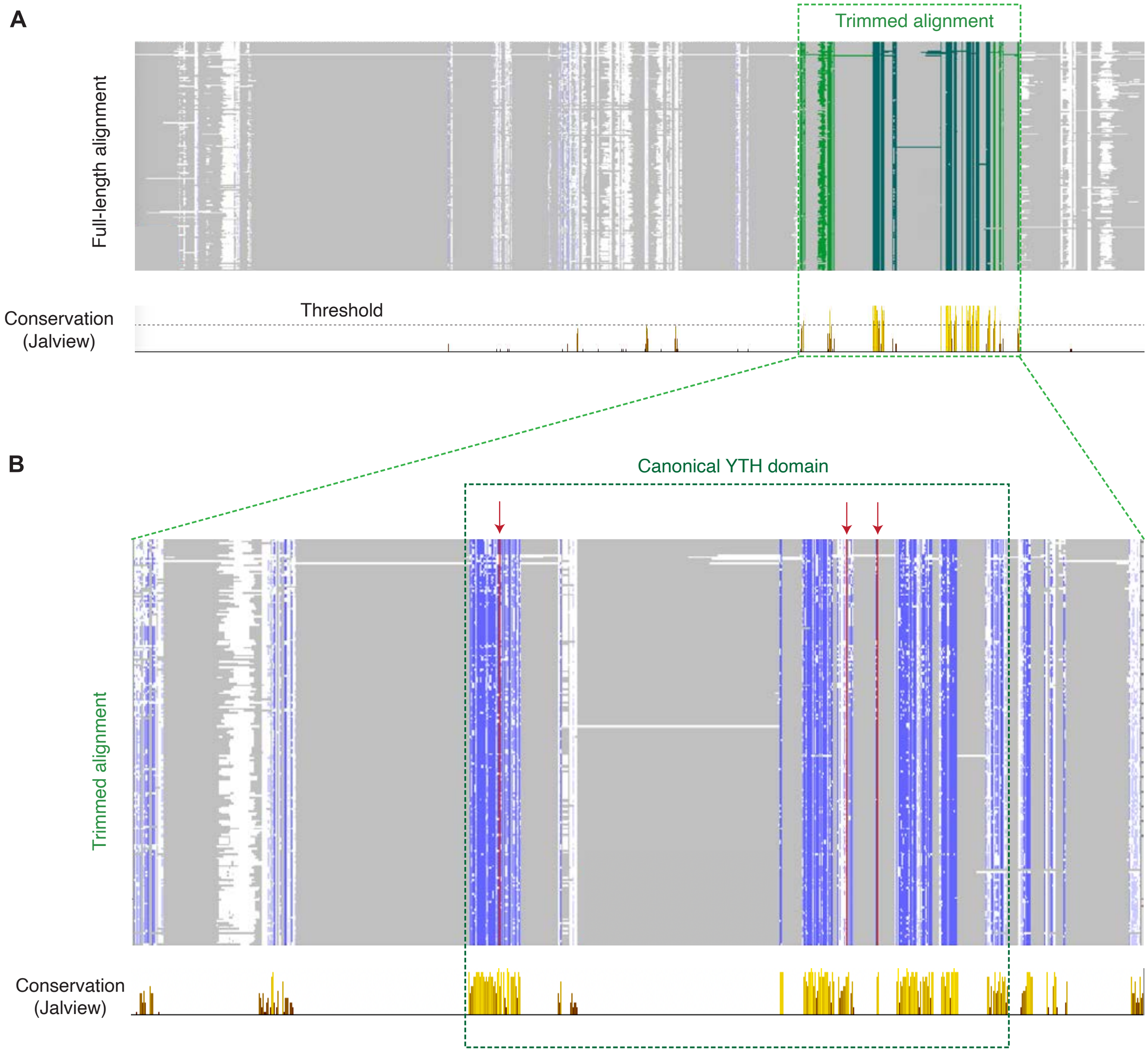


**S23 Fig. Trimming of the alignment of Viridiplantae YTHDF proteins according to conservation for phylogenetic analyses. (A-B)** Overview of the full-length amino acid sequence alignment of Viridiplantae YTHDF proteins (S2 Dataset) (A), and magnification of the trimmed region used for phylogenetic analyses (S3 Dataset) (B). The conservation score provided by Jalview [112] is shown below both panels, and the threshold used for trimming is marked with a horizontal dotted line in A. Blue-white colouring reflects percentage identity according to Jalview [112], with more intense shades of blue for the most highly conserved residues. Grey indicates gaps. In A, the trimmed region is coloured green, with a darker shade highlighting the position of the canonical YTH domain [8]. In B, the YTH domain is framed with a dashed dark green outline, and the aromatic residues that conform the m$^6$A-recognition cage are coloured red and marked with red arrows.
