## Supplemental Appendix (S1) for "Insights into the conservation and diversification of the molecular functions of YTHDF proteins"

S1 Appendix. Extended Materials and Methods

**Growth conditions.** For *in vitro* growth, seeds were sterilized by consecutive incubation in 70 % EtOH (2 min) and [1.5 % NaClO, 0.05 % Tween-20] (10 min) followed by two washes with sterile Milli-Q H_2_O, and spread on petri dishes containing Murashige & Skoog (MS) media (4.4 g/L salt mixture, 10 g/L sucrose, 8 g/L agar) supplemented with the appropriate antibiotics. Following 2-5 days of stratification at 4C in darkness, the plates were transferred to incubators at 21° C in long day conditions (16h-light/8h-dark photoperiod, 120 μmol m^-2^ s^-1^ light intensity). When further growth was needed, plants were transferred to soil and grown in incubators with the same conditions for phenotypic characterization, or standard greenhouse facilities for seed production.

**Cloning.** We employed the scar-free USER cloning method [1] to produce all *US7Yp:cYTHDF(X)-mCherry-OCSt* constructs by gluing four DNA fragments (*US7Yp*, *cYTHDF(X),* *mCherry, OCSt*) into the pCAMBIA3300-U plasmid [2]. The cloning strategy is analogous to the one described for the *ECT2p:gECT2-mCherry-ECT2t* construct [3]. This plasmid was also used as a template to PCR-amplify the *mCherry* sequence. *US7Yp* and *OCSt* fragments were amplified from the GreenGate Cloning System (Addgene) plasmids pGGA012 and pGGF005 respectively [4]. The specific fragments containing cDNA of the different YTHDF proteins (*cYTHDF(X)*) were amplified from commercial plasmids containing the cDNA clone of interest or, when unavailable, we used reverse-transcribed cDNA from RNA (see below). A list with all the templates used to amplify cDNA and the specific transcript isoform contained in the resulting constructs can be found in S1 Table. We designed the U-containing USER-primers in a way that the same three PCR fragments common to all constructs (*US7Yp*, *mCherry*, and *OCSt*) could be used in all ligations, and only the fragment encoding the cYTHDF gene had to be customized. For the chimeras, the strategy was comparable but gluing only two fragments: *US7Yp:cIDR_YTHDF(X)_* and *cYTH_YTHDF(Y)_-mCherry-OCSt*, amplified from the previously obtained *US7Yp:cYTHDF(X)-mCherry-OCSt* and *US7Yp:cYTHDF(Y)-mCherry-OCSt* constructs*.*

The construction of *ECT1p:gECT1-FLAG-TFP-ECT1t* was done by USER cloning [1] using gDNA from arabidopsis tissues to amplify the *ECT1* gene body and regulatory sequences, in identical way than described for *ECT2* in the construction of *ECT2p:gECT2-mCherry-ECT2ter* [3]. A plasmid containing *MTAp:gMTA-FLAG-TFP-MTAt* [5] was used as template to amplify the FLAG-TFP fragment.

Cloning of the trial *ECT2p:gECT1-mCherry-ECT2t* and *ECT2p:gECT4-mCherry-ECT2t* constructs was done by GreenGate [4]. In short, PCR fragments were amplified by PCR using Thermo Scientific Phusion High-Fidelity DNA Polymerase (NEB) and introduced into entry vectors by BsaI-restriction cloning, as specified in S2 Table. Of note, because none of our constructs carried an N-tag (fragment B-C), the fragment containing *ECT1/4* gDNA (from ATG to the last codon prior stop) was cloned using primers with B-to-D overhangs into pGEM-T Easy by A-tailing (Promega), thereby bypassing the need of a B-C element. The vectors containing the *ECT2* 5'UTR and upstream regulatory sequences (in pGGA000), the coding sequences (as gDNA) of *ECT1/4* (in pGEM-T Easy), linker-mCherry (pGGD003), the *ECT2* 3’UTR and downstream sequences (in pGGE000) and the D-AlaR cassette (pGGF003) were combined in a ‘Greengate reaction’ using BsaI-HF (NEB), T4 DNA-Ligase (Thermo Scientific) and pGGZ001 as destination vector [4]. All vectors were obtained from Addgene (plasmid kit #1000000036).

In all cases, the constructs were introduced into DH5α competent *Escherichia coli* cells (New England Biolabs) and Sanger-sequenced to discard clones with PCR-derived mutations. A list containing the sequences of all cloning primers, the fragments obtained with every primer set, and how these fragments were combined in the different constructs, can be found in S2 Table.

**Plant transformation and plant selection.** All final binary plasmids were introduced into *Agrobacterium tumefaciens* GV3101 to transform the appropriate plants (specified below) by floral dipping [6]. *ECT1p:gECT1-TFP-ECT1t* lines in the *ect1-2* background were selected on large MS-agar plates containing glufosinate ammonium (10 mg/L) for selection, and ampicillin (100 mg/L) to prevent *Agrobacterium* growth. Resistant primary transformants were transferred to soil at ~9 days after germination. For line selection, T2 seedlings were screened for single insertions according to the segregation of glufosinate-resistance, and visible fluorescence in a consistent pattern among many lines. Absence of free TFP was assessed by western blot, performed as described by Arribas-Hernández *et al.* [3]. *ECT2p:gECT1-mCherry-ECT2t* and *ECT2p:gECT4-mCherry-ECT2t* primary transformants in the *te234* background were selected in a similar manner but using 3 mM D-Alanine for selection.

**Complementation assay.** Due to the large amount of constructs tested, and the practical impossibility of simultaneous transformation of all constructs at once in our facilities, we performed repeated transformations of subsets of constructs to be compared, ensuring that each protein had at least two independent transformations done in parallel with the *US7Yp:cECT2-mCherry-OCSt* and *US7Yp:mCherry-OCSt* controls. The constructs dipped in each parallel transformation can be found in S7 Fig. To even out the inevitable pot-to-pot differences in transformation efficiency, we dipped between 2 and 4 pots, with 4-6 plants per pot, for each construct in every transformation, and harvested the seeds from different pots individually. To measure complementation, T1 seeds were sterilized and spread on large (135 mm) petri dishes (~2000 seeds (100 μL) / plate) containing MS-agar media supplemented with glufosinate ammonium (10 mg/L), sulfadiazine (4.83 mg/L) and ampicillin (100 mg/L). On average, we observed transformation efficiencies of ~0.8% (~15.3 transformants per plate) with roughly comparable values among constructs and transformation batches except for *ECT9, C-9/2* and *Hs YTHDF2*, for which we obtained a consistently lower amount of transformants (S9 Fig). Based on the transformation efficiency obtained in a first pilot transformation batch with only ECT2 and the mCherry control (TRF1, S9 Fig), we standardized a minimum of six plates per construct to be assessed in parallel in every independent assay, aiming to have >100 T1s of each type in any given independent comparison. Among the set of plates of each construct, we included repeats of the 2-4 independent seed batches originated from separate pots.

The seeds were stratified, germinated and grown as described above. Complementation was scored after 10 days of parallel growth for all genotypes included in the same batch. Due to the practical impossibility of measuring the exact length of more than a thousand leaves (~100 T1s x 12 constructs) within a time span shorter than that of measurable leaf growth, we decided to set a size threshold for fast assessment of complementation capacity. We considered seedlings with first true leaves of >0.5 mm (basal-to-apical length) as ‘complementing’, and those of that or inferior size as ‘not complementing’. The final score given to each construct is the average of the complementation percentages in all batches for which the construct was present, weighted by the amount of primary transformants recovered for that construct in each batch. For each score, the total amount of transformants (n) and independent transformations (I.T.) is indicated in the main figures, and the raw numbers can be found in S7 Fig and S6 Dataset.

**YTH domain modeling.** The alignment of YTH domains [7] of *Ath* ECTs with *Mpo* DFE, *Hs* YTHDF1/2, *Dm* YTHDF and *Sc* MRB1 was performed using Clustal Omega [8] on the EMBL-EBI webserver (https://www.ebi.ac.uk/Tools/msa/clustalo/) with default parameters. The overlay with the secondary structure of YTHDF1 (Protein Data Bank entry 4RCJ [9] was annotated using ESPript [10]. We manually searched the alignment for non-conservative amino acid changes between ECT1/9/11 and the rest of ECTs, and marked the most relevant positions on the topology diagram of *Hs* YTHDF1 [9].

We obtained the predicted structures of the apo YTH domains of *Ath* ECT1/2/9/11 from the AlphaFold Protein Structure Database [11, 12] and modeled surface electrostatic potential using the APBS 2.1 plugin [13] in the PyMOL Molecular Graphics System (Version 2.0 Schrödinger, LLC) on the predicted structures, with parameters 0.15 M ionic strength in monovalent salt, 310 K, protein dielectric 2, and solvent dielectric 78.

To infer the spatial position of the residues conforming the aromatic cage relative to the methylated adenosine, we modeled the structures of the YTH domains of *Ath* ECT1/2/9/11 on the crystal structure of the YTH domain of YTHDF1 in complex with m^6^A-containing RNA (Protein Data Bank entry 4RCJ) using the homology-modeling server SWISS-MODEL [14]. Sequence identity to the template was 56.6% for ECT1, 56.9% for ECT2, 61.3% for ECT9 and 53.7% for ECT11. Global Model Quality Estimations (GMQE scores) [0-1] were as follows: 0.87 for ECT1 and ECT2, 0.85 for ECT9, and 0.79 for ECT11. Higher numbers indicate higher reliability [14].

**Genotypic and phenotypic characterization.** DNA extraction and genotyping of *ect1, ect2, ect3* and *ect4* alleles for the construction of high order mutants, photographs of rosettes and seedlings, root growth characterization, and quantification of trichome branching were done with the same way and with the same equipment as described previously [3, 5]. The sequences of *ECT1*-specific primers (not used in the 2018 study) and the PCRs resulting from this genotyping can be found in S2 Table and S13 Fig.

**Fluorescence microscopy.** Stereo and confocal fluorescence microscopy were performed using the same equipment and methodology than Arribas-Hernández et al. [5]

**cDNA obtention.** cDNA was obtained using total RNA from *Arabidopsis thaliana* Col-0 wild type flowers, dissected gemma cups and apical notches *of Marchantia polymorpha* that grows spontaneously in humid areas of our greenhouses, human HepG2 cells (Sigma 85011430) or *Drosophila melanogaster* wild type larvae [15]. In all cases, the RNA was extracted using trizol, and reverse transcribed with oligo-dT primers to produce cDNA as previously described [3].

**Quantitative PCR.** Quantitative real-time RT-PCR was performed on the Bio-Rad CFX ConnectTM thermal cycler using the QuantiTect SYBR Green RT-PCR kit (Qiagen) following the instructions from the manufacturer. Expression analysis was performed following the ∆∆C_T_ method [16]. Samples were run in quadruplicates, and relative expression levels were normalized to *ACTIN* (AT3G18780) as housekeeping gene. A list of qPCR primers can be found in S2 Table.

**Western and Northern blotting.** Western blots to detect mCherry fusion proteins, and Northern blots to detect *ECT1* mRNA from total RNA of arabidopsis flowers was performed as described previously [3]. Primer sequences for the *ECT1*-specific probe are detailed in S2 Table.

**References**

1.	Bitinaite J, Nichols NM. DNA cloning and engineering by uracil excision. Curr Protoc Mol Biol. 2009;Chapter 3:Unit 3 21. doi: 10.1002/0471142727.mb0321s86. PubMed PMID: 19343708.

2.	Nour-Eldin HH, Hansen BG, Norholm MH, Jensen JK, Halkier BA. Advancing uracil-excision based cloning towards an ideal technique for cloning PCR fragments. Nucleic Acids Res. 2006;34(18):e122. doi: 10.1093/nar/gkl635. PubMed PMID: 17000637; PubMed Central PMCID: PMC1635280.

3.	Arribas-Hernández L, Bressendorff S, Hansen MH, Poulsen C, Erdmann S, Brodersen P. An m6A-YTH Module Controls Developmental Timing and Morphogenesis in Arabidopsis. Plant Cell. 2018;30(5):952-67.

4.	Lampropoulos A, Sutikovic Z, Wenzl C, Maegele I, Lohmann JU, Forner J. GreenGate - A Novel, Versatile, and Efficient Cloning System for Plant Transgenesis. PLOS ONE. 2013;8(12):e83043. doi: 10.1371/journal.pone.0083043.

5.	Arribas-Hernández L, Simonini S, Hansen MH, Paredes EB, Bressendorff S, Dong Y, et al. Recurrent requirement for the m6A-ECT2/ECT3/ECT4 axis in the control of cell proliferation during plant organogenesis. Development. 2020;147(14):dev189134. doi: 10.1242/dev.189134.

6.	Clough SJ, Bent AF. Floral dip: a simplified method for Agrobacterium-mediated transformation of Arabidopsis thaliana. Plant J. 1998;16(6):735-43. doi: 10.1046/j.1365-313x.1998.00343.x.

7.	Stoilov P, Rafalska I, Stamm S. YTH: a new domain in nuclear proteins. Trends Biochem Sci. 2002;27(10):495-7. doi: <http://dx.doi.org/10.1016/S0968-0004(02)02189-8>.

8.	Sievers F, Wilm A, Dineen D, Gibson TJ, Karplus K, Li W, et al. Fast, scalable generation of high-quality protein multiple sequence alignments using Clustal Omega. Mol Syst Biol. 2011;7(1):539.

9.	Xu C, Liu K, Ahmed H, Loppnau P, Schapira M, Min J. Structural Basis for the Discriminative Recognition of N6-Methyladenosine RNA by the Human YT521-B Homology Domain Family of Proteins. Journal of Biological Chemistry. 2015;290(41):24902-13. doi: 10.1074/jbc.M115.680389.

10.	Robert X, Gouet P. Deciphering key features in protein structures with the new ENDscript server. Nucleic Acids Res. 2014;42(Web Server issue):W320-4. doi: 10.1093/nar/gku316. PubMed PMID: 24753421; PubMed Central PMCID: PMC4086106.

11.	Varadi M, Anyango S, Deshpande M, Nair S, Natassia C, Yordanova G, et al. AlphaFold Protein Structure Database: massively expanding the structural coverage of protein-sequence space with high-accuracy models. Nucleic Acids Res. 2022;50(D1):D439-D44. doi: 10.1093/nar/gkab1061.

12.	Jumper J, Evans R, Pritzel A, Green T, Figurnov M, Ronneberger O, et al. Highly accurate protein structure prediction with AlphaFold. Nature. 2021;596(7873):583-9. doi: 10.1038/s41586-021-03819-2.

13.	Jurrus E, Engel D, Star K, Monson K, Brandi J, Felberg LE, et al. Improvements to the APBS biomolecular solvation software suite. Protein Sci. 2018;27(1):112-28. doi: <https://doi.org/10.1002/pro.3280>.

14.	Biasini M, Bienert S, Waterhouse A, Arnold K, Studer G, Schmidt T, et al. SWISS-MODEL: modelling protein tertiary and quaternary structure using evolutionary information. Nucleic Acids Res. 2014;42(W1):W252-W8. doi: 10.1093/nar/gku340.

15.	Arribas-Hernández L, Rennie S, Köster T, Porcelli C, Lewinski M, Staiger D, et al. Principles of mRNA targeting via the Arabidopsis m6A-binding protein ECT2. eLife. 2021;10:e72375. doi: 10.7554/eLife.72375.

16.	Pfaffl MW. A new mathematical model for relative quantification in real-time RT–PCR. Nucleic Acids Res. 2001;29(9):e45-e. PubMed PMID: PMC55695.
